# Finding needles in haystacks: identification of novel conserved PETase enzymes in *Streptomyces*

**DOI:** 10.1101/2023.11.08.566248

**Authors:** Jo-Anne Verschoor, Martijn R. J. Croese, Sven E. Lakemeier, Annemiek Mugge, Charlotte M. C. Burgers, Paolo Innocenti, Joost Willemse, Marjolein E. Crooijmans, Gilles P. van Wezel, Arthur F. J. Ram, Johannes H. de Winde

## Abstract

The rising use of plastic results in an appalling amount of waste which scatters into the environment affecting environmental, animal, and human health. One of these plastics is PET which is mainly used for bottles and textiles. In this research, we investigate the PET degrading ability of the *Is*PETase homolog *Sc*LipA from *Streptomyces coelicolor.* Of 96 different *Streptomyces* strains screened, 18 % were able to degrade the model substrate BHET. Three different variants of lipase A, named *Sc*LipA*, S2*LipA and *S92*LipA were identified and analyzed in detail. The *lipA* gene was deleted from *S. coelicolor* M145 using CRISPR/Cas9, resulting in reduced BHET degradation. *LipA* overexpression in the knock-out background significantly enhanced BHET degradation. All three enzymes were expressed in *E. coli* BL21 for protein purification and biochemical analysis, showing that enzymatic activity most likely resides in a dimeric form of the enzyme. The optimum pH and temperature were determined to be pH 7 and 25 °C for all three variants. Using these conditions, the activity on BHET and amorphous PET film was investigated. *S*2LipA efficiently degraded BHET and caused roughening and small indents on the surface of PET films, consistent with PET-degrading activity. The frequent occurrence of the *S2*LipA variant in *Streptomyces* suggests an environmental advantage towards the degradation of more hydrophobic substrates such as these polluting plastics in the environment.

## Introduction

Petroleum-derived plastics are among the most useful and widespread synthetic polymers in the modern world. They are relatively easy and cheap to produce, extremely versatile and durable materials, which makes them remarkably attractive for a wide range of applications [1], [2]. This has caused the demand and production of plastics to rise steadily to an estimated 9.2 billion metric tons (Mt) produced as of 2023 [1], [3], [4]. Of all plastic produced, 75 % (6.9 billion Mt) has been mismanaged or landfilled [4]. Durability and resistance to degradation of these polymers, combined with the increasing output of waste generated by humans, makes plastic accumulation and pollution a major global environmental concern [5]. In fact, commonly used plastics are generally not biodegradable, accumulate in landfills or leak into terrestrial and/or aquatic environments, where they disintegrate into micro- and nano-plastics, thereby posing a serious threat to many ecosystems [5].

Polyethylene terephthalate (PET) is extensively used to produce bottles, food packaging, clothing and films. The high demand for PET makes it one of the most common plastics that we encounter in everyday life with 18.8 million tons being produced worldwide in 2015 alone [6]. PET waste management in most countries today consists mainly of incineration and landfilling. Both practices are causing detrimental effects to the environment, such as leaching and the release of toxic compounds [7]. On the other hand, an increasing number of countries have recycling systems in place to prevent environmental accumulation and make renewed use of the PET waste produced. Despite such efforts, the recovery ratio is far less than 50 % and only a minor fraction is used to manufacture new products, with substantial amounts of PET still entering the environment [8].

With plastic waste accumulating in the environment, investigating the adaptation and response of soil and aquatic microorganisms to plastic exposure has proven eminent. Microbial response and adaptation mechanisms are likely to help battle plastic pollution and initiate remediation and recycling options. Enzymatic degradation already has proven to enable natural routes to depolymerize these recalcitrant polymeric materials. Several PET-hydrolyzing enzymes (PHEs) have been identified, initiating biodegradation of this polymer. Predominantly, these are esterases, lipases, cutinases and cutinase-like enzymes exhibiting hydrolysis of the ester bond between terephthalic acid (TPA) and ethylene glycol (EG) moieties within the PET polymer [9]–[14]. It was not until 2016 that Yoshida and colleagues reported the discovery of a novel bacterium, *Ideonella sakaiensis 201-F6* (hereafter *I. sakaiensis*), which had evolved towards the degradation and utilization of PET as a sole carbon source [15]. The enzyme identified as responsible for the first step in the catabolism of PET is the *Is*PETase (ISF6_4831), classified as an extracellular esterase. This enzyme hydrolyses PET into its monomer mono-(2-hydroxyethyl) terephthalic acid (MHET), with only trace amounts of bis(2-hydroxyethyl) terephthalate (BHET) and terephthalic acid (TPA) being observed. To complete the breakdown of PET, *I. sakaiensis* possesses another enzyme, the *Is*MHETase (ISF6_0224), which is capable of hydrolyzing MHET into TPA and ethylene glycol (EG), common starting chemicals to polymerize PET [15]. Compared to other PHEs which have a broader range of substrates, the *Is*PETase displays significantly higher activity against BHET, PET films, and commercial bottle-derived PET at 40 °C [15]. Currently, all PETase-like enzymes are described in the PAZY-database [16].

Many genera of soil and aquatic bacteria are exposed to accumulated plastics. An important phylum of bacteria remaining to be investigated is the *Actinobacteria.* This phylum consists of various complex genera such as *Thermobifida, Rhodococcus, Streptomyces* and many more. Several enzymes of actinobacterial origin such as the LCC, *Tf*Cut2 and many more have previously been identified and characterized to exhibit PET degrading abilities earning their place in the PAZY database [9]–[11], [13], [16]–[19]. However, for various other genera including *Streptomyces* only limited information is available [20]–[25]. Only one *Streptomyces* enzyme, SM14est has been described [21].

*Streptomycetes* are filamentous growing soil-dwelling bacteria with a characteristic lifecycle. They are renowned for their capability to produce antibiotics but are also known to excrete enzymes to degrade complex natural polymers, including plant biomass [26]–[28]. Depending on the soil pollution rate and the distance to the polluted area, it is possible to recover 7,5 x 10^4^ to 3,0 x 10^4^ plastic microparticles per kilogram (kg) of soil [29]. This equates to 2-20 milligrams (mg) of recoverable plastic particles per kg of soil [29]. Hence, the rhizosphere and its associated microbes, such as *Streptomyces* are extensively exposed to plastics. This exposure to plastics and their ability to degrade complex biomass with hydrolytic enzymes makes them a promising genus to explore for novel PHEs [20]–[25], [30], [31]. These possible PHEs are expected to be secreted enzymes. Expression of enzymes in *Streptomyces* is a highly regulated process and hence, such regulation will likely play an important role in putative PET degrading activity [32].

In this research, we identified Lipase A from *Streptomyces coelicolor* (*Sc*LipA) as a homolog to the *Is*PETase. Consequently, we investigated the ability of *S. coelicolor* to degrade the PET model compound BHET. Conservation of this BHET degrading ability was investigated by screening a collection of 96 previously isolated *Actinobacteria* (predominantly *Streptomyces*) under various conditions [33]. This yielded insight into the optimal conditions needed to induce BHET degradation as well as in the abundance of strains exhibiting BHET-degrading capability. In one or more of the provided conditions, 44 % of the strains could degrade BHET, 18 % of the evaluated strains were able to degrade BHET individually confirming a wider distribution of this ability in nature. The presence of LipA in these strains was explored and the corresponding genes were sequenced. The enzyme is highly conserved and can be classified into three different variants. The *S. coelicolor* variant further named *Sc*LipA, the MBT92 variant (*S*92LipA) which is only present in MBT92, and a conserved variant present in all other active strains (*S2*LipA). The functionality of all three LipA variants in the degradation ability of BHET and amorphous PET in *Streptomyces* species was investigated. A knock-out of the *lipA* gene was constructed in *S. coelicolor* M145, and overexpression constructs of the three variants were expressed in the knock-out to study the effect of the genes *in vivo.* The individual enzymes were expressed in *Escherichia coli* and purified for the analysis of substrate specificity, thermostability, and other properties. Enzyme characteristics were examined using enzyme assays and liquid chromatography-mass spectrometry (LC-MS). Additionally, the strains, as well as the purified enzymes, were incubated on amorphous PET films to investigate their true ability to adhere to and degrade PET.

In conclusion, we present a full comprehensive identification and characterization of a novel, conserved PET-degrading enzyme family of *Streptomyces*. Structural characteristics and *in vivo* and *in vitro* activity provide insight into the evolutionary and ecological adaptation of soil bacteria towards plastics in their environment.

## Results

### Scavenging the *Streptomyces* genome for candidate *Is*PETase homologs

Homologs of the *Is*PETase (Accession number A0A0K8P6T7, [34]) encoded by the genomes of *Streptomyces coelicolor*, *Streptomyces scabies* and *Streptomyces avermitilis* were identified using protein BLAST. The most promising enzyme was a *S. coelicolor* enzyme, annotated as a putatively secreted lipase A (LipA, accession number Q9L2J6, [35]), showed a query coverage of 91 % and an identity of 48 % with the *Is*PETase (Fig. 1A) [36]. The 3D structure of the *Is*PETase (Fig. 1B) and the AlphaFold model of *Sc*LipA (Fig. 1C) were superimposed, to compare the three-dimensional structures of the enzymes [37]. When overlayed, the overall structures looked very similar (Fig. 1D). The catalytic triad was conserved, as well as two of the three residues of the previously described substrate binding site (Fig. 1E and F) [36]. The difference in the binding site is a phenylalanine in LipA instead of a tyrosine in *Is*PETase, as shown in figure 1E. The high similarity and the presence of the important residues for PET degradation hint towards the putative PET degrading activity of the *Streptomyces coelicolor* lipase A (*Sc*LipA). On the N-terminal end of the protein, a twin-arginine translocation (TAT) signal is present indicating that the natural substrate of this enzyme is located outside of the cell [38]. Indeed, for plastics-degrading enzymes, extracellular activity appears to be essential.

**Figure 1:**
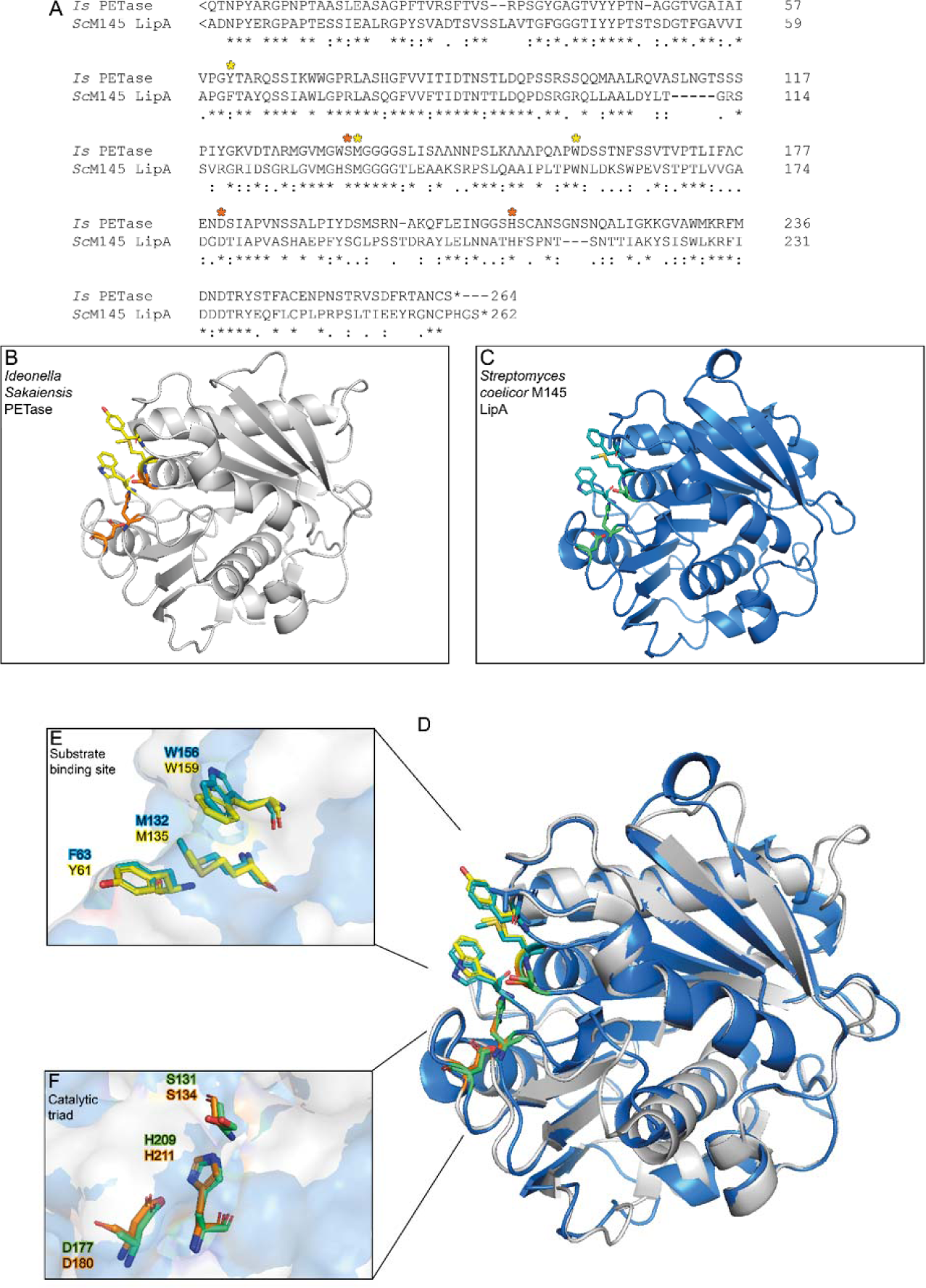
Sequence and structure comparison of IsPETase and ScLipA. A) sequence comparison of the IsPETase and ScLipA indicating the binding sites with yellow asterisks and the catalytic triad with orange asterisks. B) the structure of the IsPETase as provided by Uniprot (5XJH). The catalytic triad is displayed in orange, and the binding site is in yellow. C) Predicted model of the structure of ScLipA constructed with AlphaFold. The catalytic triad is shown in green and the binding domains are displayed in cyan. D) An overlay of the IsPETase and ScLipA. E) overlay of the binding domains, the tyrosine on position 63 of the IsPETase overlaps with a phenylalanine on position 61 if the ScLipA. F) overlay of the catalytic triads.

### BHET degrading activity of *S. coelicolor*

To further investigate the activity of *S. coelicolor* M145 on BHET, the strain was grown on *Streptomyces* minimal medium (StrepMM) agar plates containing BHET. Due to its limited solubility, the addition of BHET causes the plates to become turbid. When BHET is converted to MHET and/or TPA, a halo of clearance appears. The secreted enzyme activity was expected to be tightly regulated in response to environmental growth conditions [32]. Therefore, we tested *N*-acetyl glucosamine (GlcNAc) as a possible inducer of secreted enzyme activity. GlcNAc is a monomer of chitin and peptidoglycan, which under poor nutritional conditions induces development and antibiotic production in *Streptomyces coelicolor* [39], [40], whereas in nutrient-rich environments these responses are blocked [35]. We anticipated that GlcNAc may act as an inducer for the production of extracellular enzymes such as lipases, esterases and possibly BHET-degrading enzymes. Hence, *S. coelicolor* M145 was grown in the presence and absence of BHET [10 mM], mannitol [25 mM] and N-acetyl glucosamine [25 mM] in all possible combinations.

Interestingly, a clearance halo was observed at 18 days of growth as shown in figure 2A. Notably, figure 2A displays that the addition of GlcNAc as inducer seemed to induce the degradation of BHET. Additionally, *S. coelicolor* produces agarase, which resulted in ‘sinking’ halos which you can see in the controls without BHET (Fig. 2A, StrepMM) [41]. These ‘sinking halos’ looked very similar to clearance halos. Therefore, the ability *S. coelicolor* to degrade BHET in liquid culture was investigated using liquid chromatography-mass spectrometry (LC-MS). For the liquid analysis, strain M145 was precultured and inoculated in minimal liquid medium without polyethylene glycol (NMM) with and without BHET and GlcNAc. Samples were taken after 24 h and 72 h and analyzed using LC-MS. In the negative ionization mode, the mass of TPA (166 g/mol) could be observed around 4 min retention time showing a small peak on the UV spectrum, MHET (210 g/mol) appeared around 5.2 min with a strong signal on both the MS as well as the UV. Finally, BHET (255 g/mol) could be observed in the positive ionization mode between 5.5 and 6 min with a strong signal at 240 nm (Fig. 2B and 2C). Around 9.5 min, impurity was observed, which was present in the BHET stock (Fig. 2C). When grown in NMM medium containing BHET, 30-50 % of the BHET was degraded within 24 h, after 72 h, all BHET was degraded and only MHET was present. Indeed, the addition of GlcNAc enhanced the degradation of BHET (Fig. 2D).

**Figure 2:**
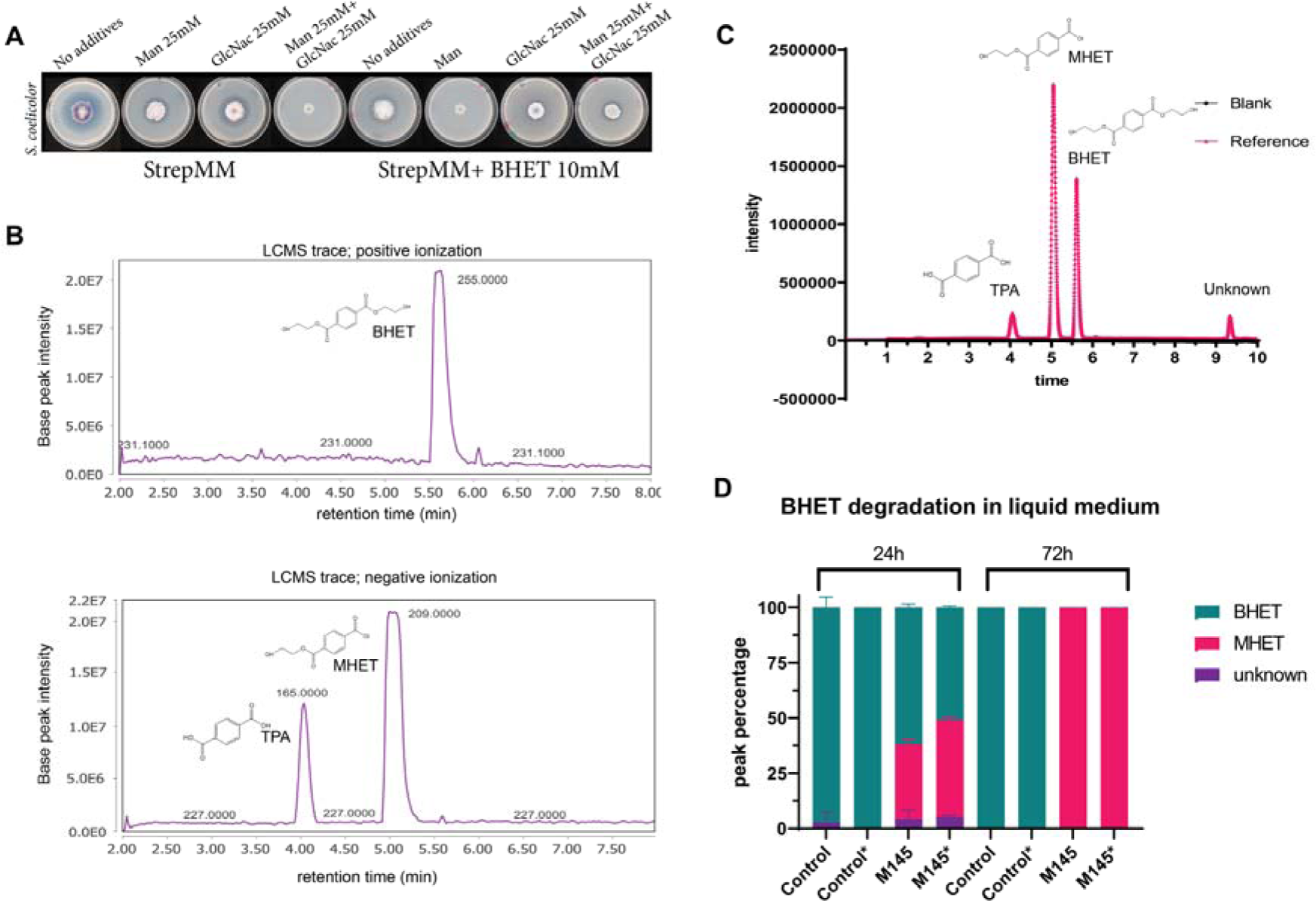
The ability of S. coelicolor M145 to degrade BHET. A) Degradation of BHET by S. coelicolor after 18 days of growth on Strep MM Difco agar with and without Mannitol, BHET and GlcNAc B) LC-MS trace of BHET in positive ionization, MHET and TPA in the negative ionization. C) The UV spectrum at 240 nm of TPA, MHET and BHET with corresponding retention times. D) The control samples are displayed as control and only contain NMM with BHET [10 mM]. The addition of GlcNAc [25 mM] to the cultures is indicated with an asterisk. Peak intensity and area percentage were calculated using GraphPad. The areas are presented in percentage compound present in culture. BHET is indicated in turquoise and MHET is indicated in magenta. Some impurities are always present as a peak around 9.5 min retention time, this compound is called unknown and presented in purple.

### Bulk screen and individual screening of *Actinobacteria* strain collection

With *S. coelicolor* M145 exhibiting a clear but modest BHET degrading activity on plates, we set out to screen a diverse collection of 96 Actinobacteria of our strain collection [42] to investigate the spread of this characteristic and identify additional BHET degrading bacteria. 36 strains were randomly chosen from this collection to perform an initial BHET toxicity test. Growth of these strains appeared to be delayed with 10 mM BHET, and impaired with 20 mM BHET and higher concentrations. Hence, a concentration of 10 mM of BHET was chosen for screening, with all tested strains able to grow and halos readily observed (Fig. 3A).

**Figure 3:**
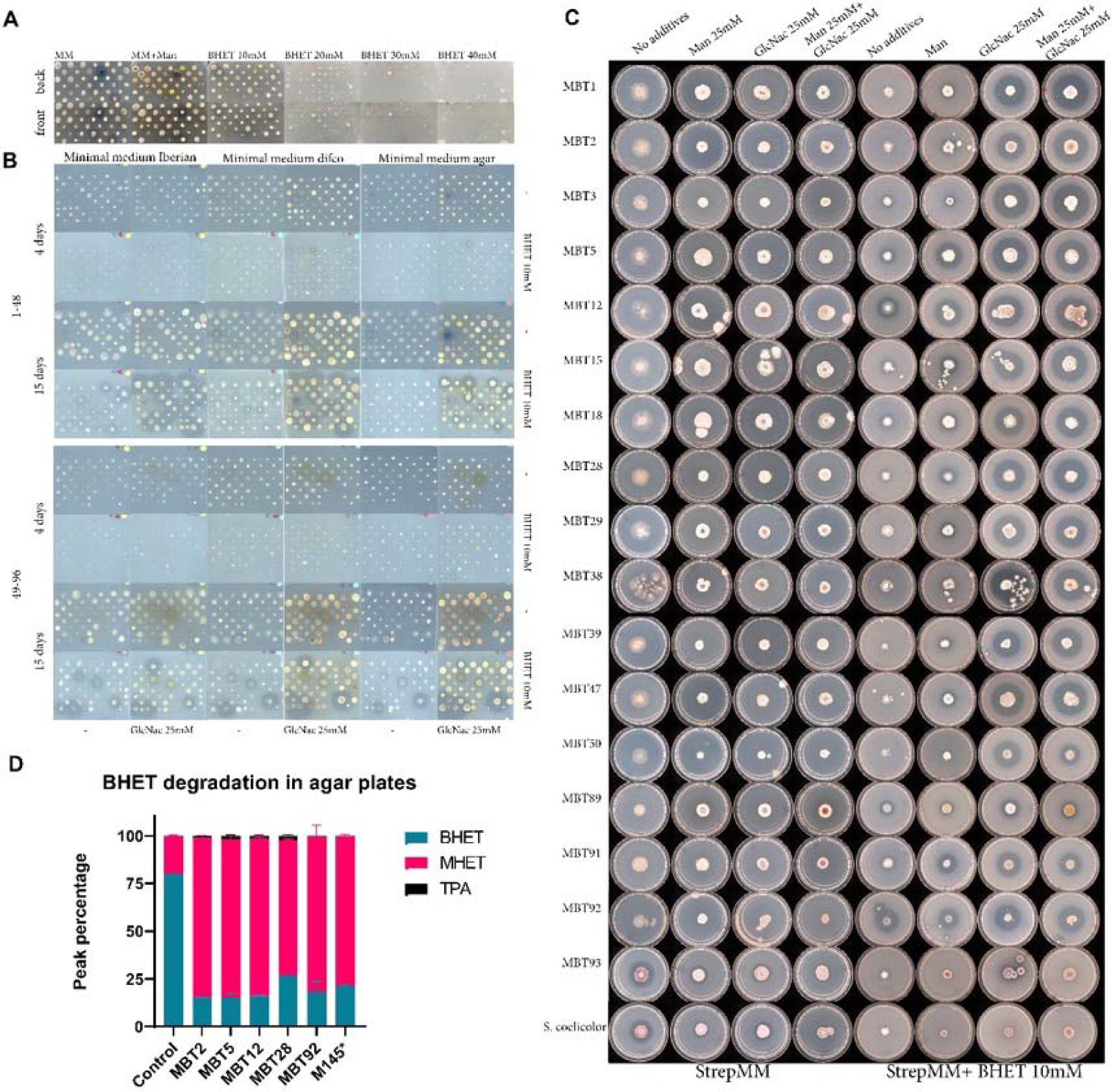
Actinobacteria screens on BHET. A) Toxicity screen of 36 Actinobacteria on BHET concentrations ranging from 0 to 40mM of BHET. B) Bulk screens of 96 Actinobacteria on StrepMM Iberian agar, Difco agar and agar agar with and without N-acetyl glucosamine as inducer at 4 and 15 days of growth. C) Individual screen of all active strains on Strep MM Difco agar with and without Mannitol, BHET and GlcNAc after 10 days of growth. D) Analysis of BHET degradation in agar plugs after 15 days of growth using LC-MS. The peak percentage was calculated using GraphPad. *The agar plug of M145 was taken after 18 days.

A bulk screen for BHET degradation was performed with all 96 strains split over two 96-well plates to provide them with enough space for development. Seven strains did not grow on the plates and were taken out of the screen. Spore suspensions were stamped onto plates containing StrepMM with different agar brands containing combinations of BHET [10 mM], Mannitol [25 mM] and GlcNAc [25 mM]. The type of agar used clearly impacted the observed degradation patterns. The agar brand seems to have a clear influence on the degradation pattern (Supplement 1). Difco agar was chosen as the agar source for all further experiments since most strains showed growth on Difco agar. The addition of GlcNAc resulted in a different halo pattern and more predominant halos during the bulk screen and induced BHET degradation in most strains (Fig 3B, Supplement 1).

*Actinobacteria,* especially *Streptomyces,* depend on environmental cues and interspecies interactions for the induction of enzymes and secondary metabolites [43]. Consequently, it was important to investigate their individual BHET degrading activity on separate plates. All strains showing halos in the bulk BHET screen were grown on individual plates (Supplement 2). Of 38 active strains, only 17 strains exhibited individual activity on BHET after 10 days of growth (Fig. 3C). Interestingly, most strains required the addition of GlcNAc to express the BHET degrading enzyme. Only 5 strains, MBT5, 12, 15, 91 and 92 degraded BHET in the absence of GlcNAc. MBT92 appeared to exhibit constitutive expression showing a similar-sized halo in all conditions. To substantiate a good correlation between halo size and biochemical degradation, strains MBT2, 5, 12, 28, 92 and *S. coelicolor* M145 were incubated with approximately 2*10^6^ spores on StrepMM Difco plates containing BHET and GlcNAc. Samples of agar were taken after 15 days using an agar excision tool to cut out part of the halo. Since *S. coelicolor* M145 displayed no activity after 15 days (see also above), halos were excised after 18 days. Samples were spun down using nylon spin columns, separating all liquids from the agar, and prepped for LC-MS analysis (Fig. 3D). In all strains, clear conversion from BHET to MHET was observed. MBT2, 5, 12, and 28 show traces of TPA suggesting further conversion of MHET.

### Identification of LipA in BHET-degrading strains

A PCR with degenerative primers was performed to investigate if *lipA* is present in the strains showing activity in the individual screens displayed in figure 3C. The PCR results are displayed in supplemental data S4. The PCR fragments were cloned into a linearized pJET2.1 and sequenced. Sequence alignment demonstrated substantial divergence within the signal sequence while presenting only minor variations in the overall enzyme coding region (S5). The enzyme sequences were aligned and could be clustered into three variants the *S. coelicolor* variant (*Sc*LipA), the MBT92 variant (*S*92LipA) which is only present in MBT92, and a conserved variant present in all other active strains (*S2*LipA). (S6). The enzyme structures predicted with AlphaFold showed minor changes in the enzyme structure with minor influence on the overall structure (Fig. 4A-H) [37]. The amino acid alignment showed some variations between the proteins especially surrounding the catalytic triad (Fig. 4I). The signal peptides, identified with SignalP 5.0, contained several differences [44]. The alignment containing the signal sequence is provided in the supplementary data (S6).

**Figure 4:**
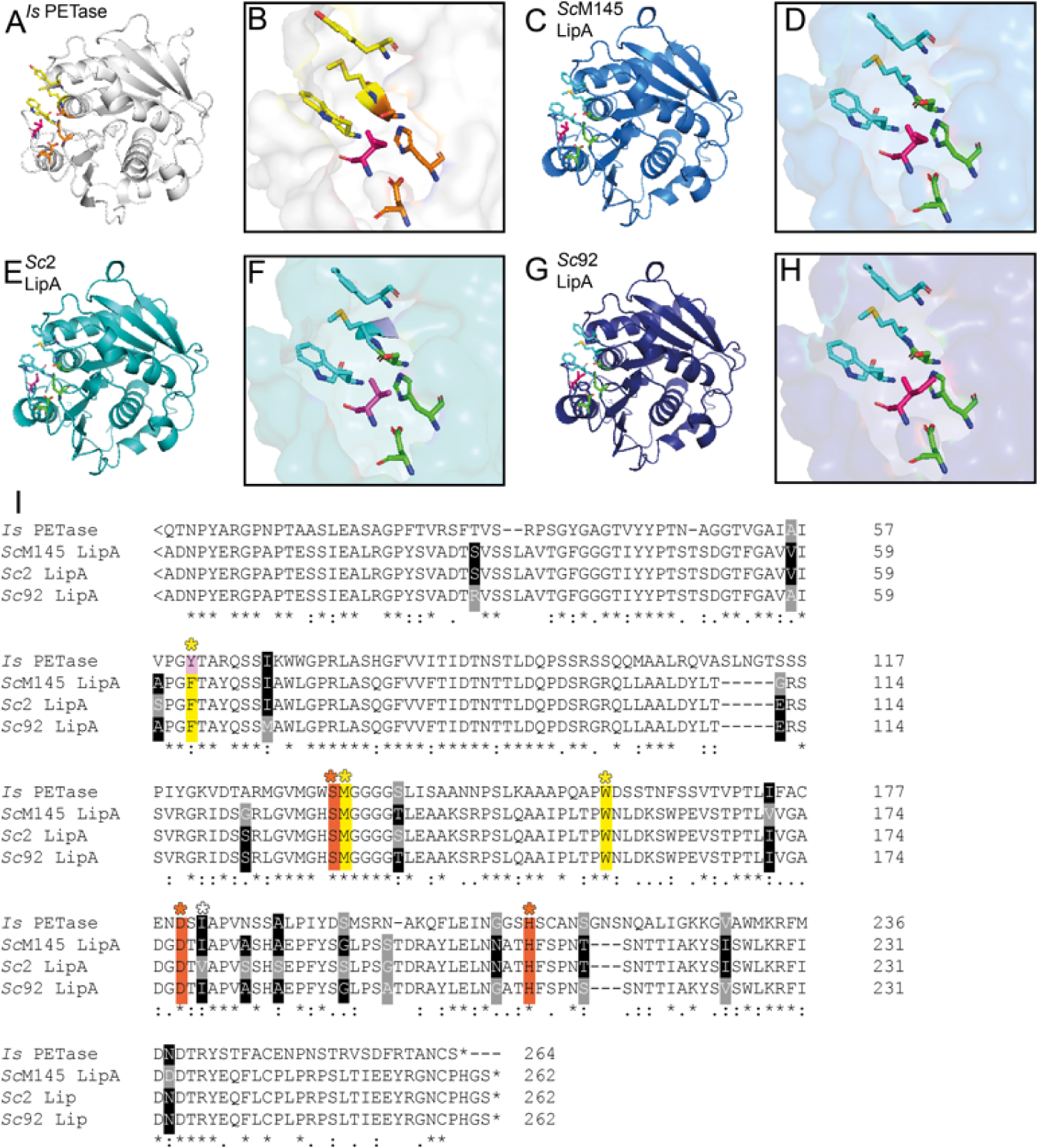
Comparison of Lipase A variants to the IsPETase. A) the structure of the IsPETase as provided by AlphaFold (5XJH) indicating the binding residues in yellow and the catalytic triad in orange, the pink residue that might influence binding. B-D) Predicted model of the structure of ScLipA (B), S2LipA (C) and S92LipA (D) constructed with AlphaFold. E-H) Enlarged image of the catalytic triad and binding site from IsPETase (E), ScLipA (F), S2LipA (G) and S92LipA (H). I) sequence comparison of the IsPETase and ScLipA, S2LipA and S92LipA indicating the binding residues with yellow asterisks/highlight and the catalytic triad with orange asterisks/highlight. The white asterisk shows a residue that might influence binding. Sequence differences from the different lipases are highlighted in grey, when only one sequence differs the conserved amino acids are highlighted in black.

### Scanning electron microscopy imaging of *Streptomyces* species and enzymes on amorphous PET film

MBT2, 5, 12, 28 and 92 were incubated in NMM with and without GlcNAc. All strains form a biofilm surrounding the edges of the plastic films, thereby fixing them to the 12 wells plates. After fixation, the samples were sputter-coated and visualized with scanning electron microscopy (SEM). All strains appeared to adhere well to the plastic films, forming hyphal structures colonizing the film (Fig. 5). Some areas were heavily overgrown and did not allow for visualization of the film. The less cultivated areas did not show significant damage over the control film suggesting that either more biomass is needed to observe degradation, or no PET degradation was achieved.

**Figure 5:**
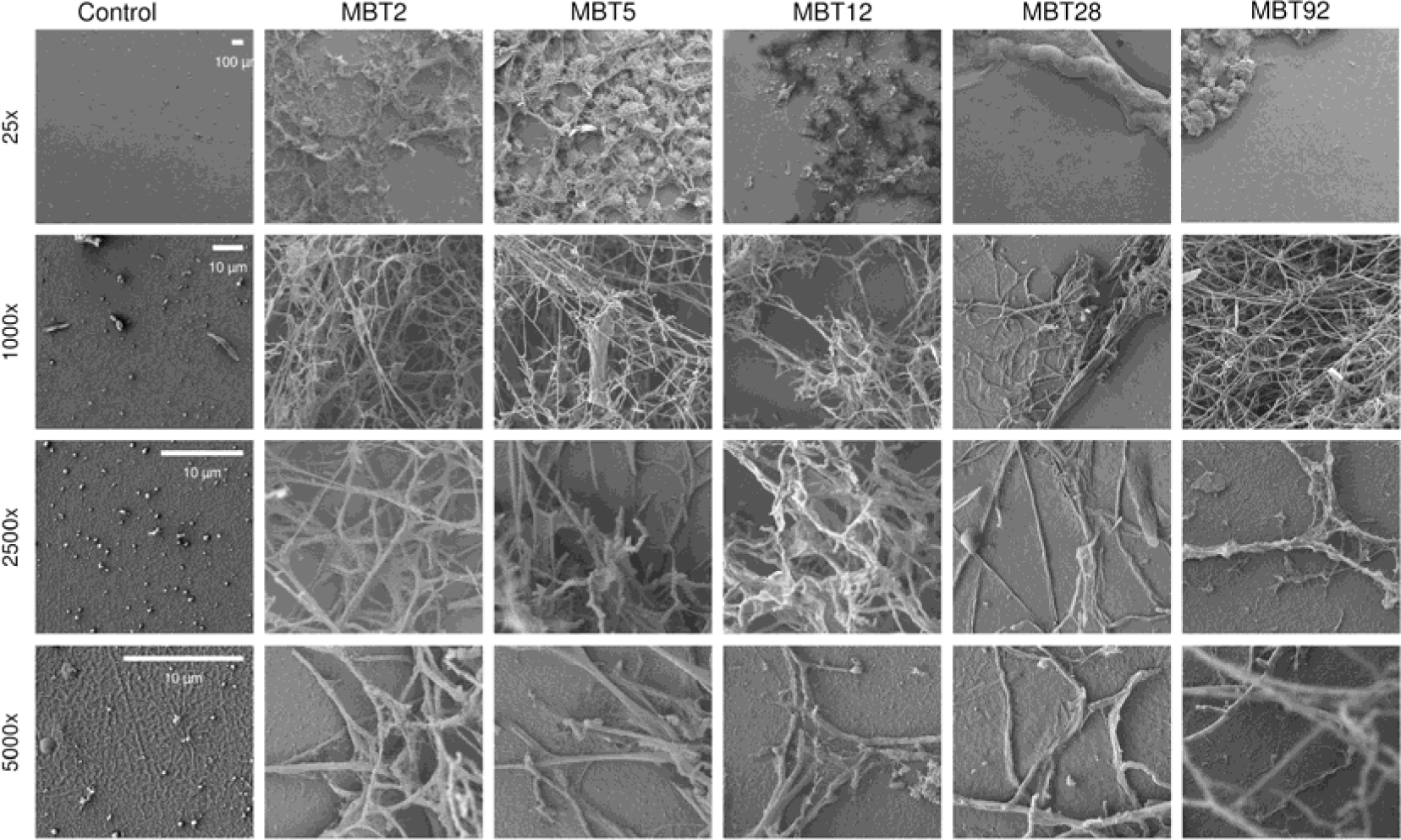
Scanning electron micrographs of MBT2, 5, 12, 28 and 92 on amorphous PET film. PET films incubated for 2 weeks at 30 °C in NMM with 0.05 % [w/v] glucose with 10^7^ spores and inoculated for 2 weeks at 30 °C. From left to right: control, strain MBT2, MBT5, MBT12, MBT28 and MBT92. From top to bottom the magnifications 25 x, 1000 x, 2500 x and 500 0x.

### Knock-out and overexpression of the three *lipA* variants in *S. coelicolor*

To obtain a better understanding of the role and function of LipA *in vivo,* a *lipA* knock-out mutant was constructed in *S. coelicolor* M145. The knock-out was validated via diagnostic PCR and sequencing (S 7.1).

Additionally, strains were constructed expressing one of the three different *lipA* variants under the control of the pGAP promotor (S 7.2). pGAP is a semi-constitutive promotor that is induced by simple sugar molecules [45]. To avoid interference of the native *Sc*LipA and other regulatory elements, all over-expression constructs were cloned in strain Δ*lipA*. This resulted in strain S3 expressing *SclipA,* S5 expressing *S2lipA* and S7 expressing *S92lipA*. Two expression media were chosen namely NMM as minimal medium with a defined carbon source and rich medium tryptic soy broth sucrose (TSBS) which is a protein expression medium. Expression was validated on both media using SDS page and Western blot (Fig. 6A-B and 6D-E). Both NMM with 5 % (w/v) glucose and TSBS show clear signal on western blot indicating that the enzyme is expressed.

**Figure 6:**
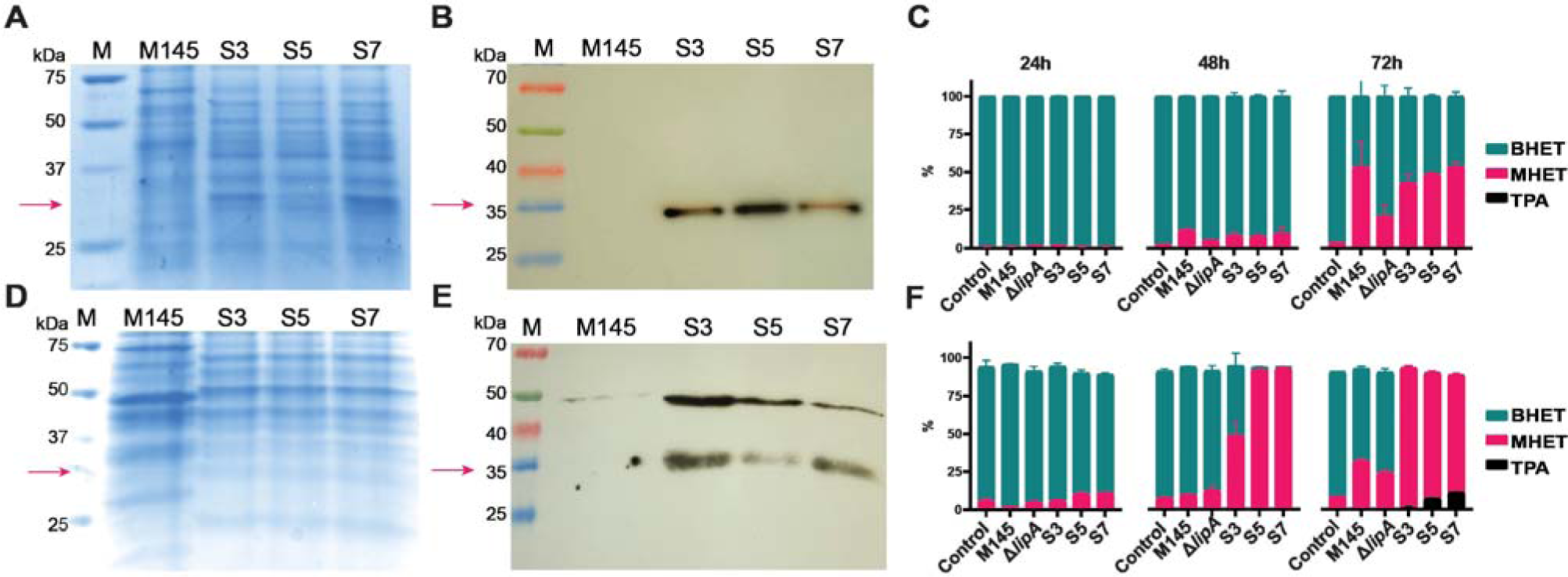
Expression of the LipA variants in S. coelicolor M145 ΔlipA in TSBS medium. A) SDS-PAGE of concentrated samples on NMM medium, faint band around ∼32 kDa in the overexpression strains. B) Western blot of NMM samples showing clear signal around 32 kDa C) Analysis of BHET degradation in NMM of the wildtype strain, ΔlipA, S3, S5 ans S7 using LC-MS. Samples were taken at 24 h, 48 h and 72 h. The percentage BHET is presented in turquoise, the percentage MHET in magenta and the concentration TPA in black. The area percentage was calculated using GraphPad. D) SDS-PAGE of concentrated samples on TSBS medium, faint band around ∼32 kDa in the overexpression strains. B) Western blot of TSBS samples showing clear signal around 32 kDa C) Analysis of BHET degradation in TSBS of the wildtype strain, ΔlipA, S3, S5 ans S7 using LC-MS. Samples were taken at 24 h, 48 h and 72 h. The percentage BHET is presented in turquoise, the percentage MHET in magenta and the concentration TPA in black.

BHET degradation was examined *in vivo* by inoculating 1,0*10^8^ spores in 50 ml of NMM with glucose [5 % (v/v)] and BHET or TSBS containing BHET. Samples were taken at 24, 48 and 72 h and analyzed using LC-MS. The percentages BHET, MHET and TPA were calculated over the total peak area. After 24 h, while the spores are still germinating, limited activity is observed. After 48 h, on NMM, the knock-out shows significantly less activity on BHET than the wildtype strain. After 72 h, there is a clear difference in the degradation pattern of the knock-out compared to the wildtype strain. The overexpression constructs show 40-50 % degradation, which is similar to the wildtype strain, although it is significantly higher than in the knock-out strain (parental strain) (Fig. 6C). Some residual activity, not related to LipA, was still observed in the knock-out strain.

After 48 h on TSBS, there was no difference observed in the BHET degrading ability of the wildtype strain compared to the knock-out strain; After 72 h, the wildtype strain degraded significantly more BHET than the knock-out strain. Unlike minimal medium, the strains containing the overexpressed *lipA* variants showed drastic BHET degradation on TSBS after 48 h, with some variation between the different enzyme variants (Fig. 6F). After 72 h, the knock-out converted approximately 23 % of the BHET to MHET, whereas the wild-type strain converted approximately 32 %, the overexpression strains degraded all BHET after 72 h and TPA could be observed in the medium. The knock-out still showed some activity suggesting the presence of at least one other enzyme with BHET degrading activity. A two-way ANOVA clearly indicated the statistical significance of the results (S8).

To observe physical changes within the cultures, automated time-lapse microscopy imaging was used to follow and visualize the development of the strains and degradation of the BHET. This microscope can make brightfield images at time intervals resulting in a timelapse video (S9). It takes all strains around 10 h to start developing (Fig. 7). Initially, no or only slight changes were observed in the presence and shape of the BHET crystals but over time the BHET crystals roughened and disappeared. The overexpression strains degraded the BHET crystals between 15 and 20 h, where the edges of the crystals started to roughen until the entire crystal fell apart (Fig. 7). The knock-out strain also degraded crystals, but complete degradation was only observed after 25 h. The wildtype strain degraded the BHET in approximately 23 h. For time-lapse videos, see supplementary data (S9).

**Figure 7:**
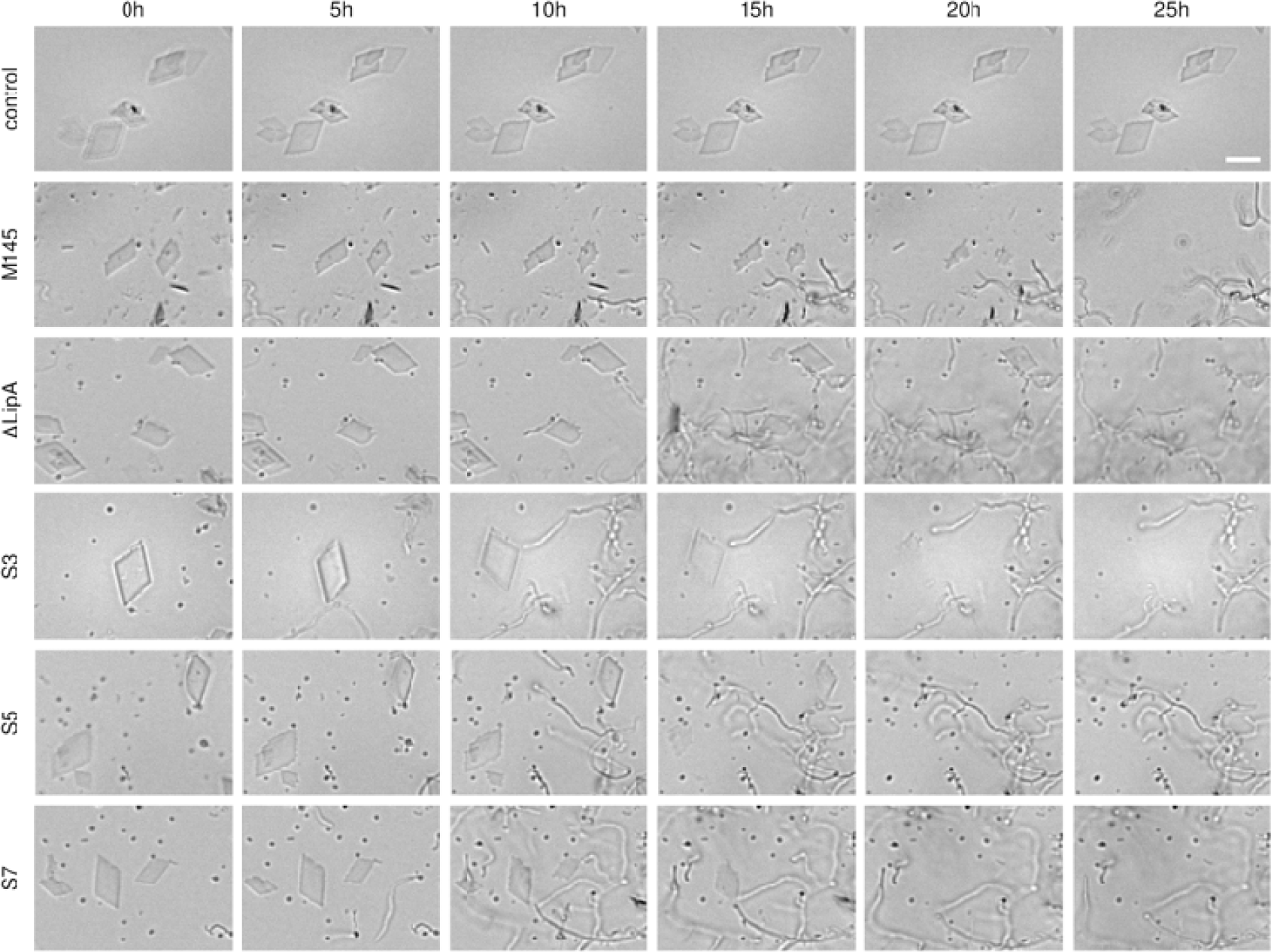
Time-lapse images of BHET degradation by S. coelicolor M145, ΔlipA, S3, S5 and S7. Time-lapse images at 5-hour intervals of wild type and ΔlipA mutant and overexpression strains S3, S5 and S7 in the presence of BHET particles (diamond-shaped). A negative control lacking bacterial inoculation was used to demonstrate that the BHET particles do not undergo natural degradation over time. The scale bar is 10 µm.

### Expression and in vitro characterization of Lipase A variants

For extensive *in vitro* analysis of the enzymes, the corresponding gene sequences were trimmed to remove their signal peptide, codon-optimized, synthesized, cloned into pET16b (adding an N-terminal His-tag to the protein) and expressed in *E. coli* BL21 A-I. As positive controls, the genes encoding for *Is*PETase and *Tf*Cut2 were synthesized and expressed in the same way. Both *Is*PETase and *Tf*Cut2 have PET degrading activity, however, their substrate preference, efficiency, optimal conditions, and stability vary. *Is*PETase has PET as the preferred substrate whereas *Tf*Cut2 is a cutinase that also displays high activity on cutinase-like substrates such as the para-nitrophenyl substrates and lipase-like substrates as tributyrin. Clear bands around 32 kDa indicated the presence of the corresponding enzymes (Fig. 8A), which was confirmed by Western blot (Fig. 8B). His-tag purification resulted in samples containing three bands for the LipA enzymes (Fig. 8C). The higher band (∼60 kDa) could also be observed in the western blot and most likely corresponded to a dimeric form or aggregate of the enzyme (Fig. 8C). Native zymogram analysis clearly showed that enzyme activity resided with the ∼60 kDa band for *S*2LipA, further indicating the dimeric form of the LipA enzymes as being active (Fig 8D). *Tf*Cut2 did not migrate through the gel, which might be due to the difference in isoelectric point.

**Figure 8:**
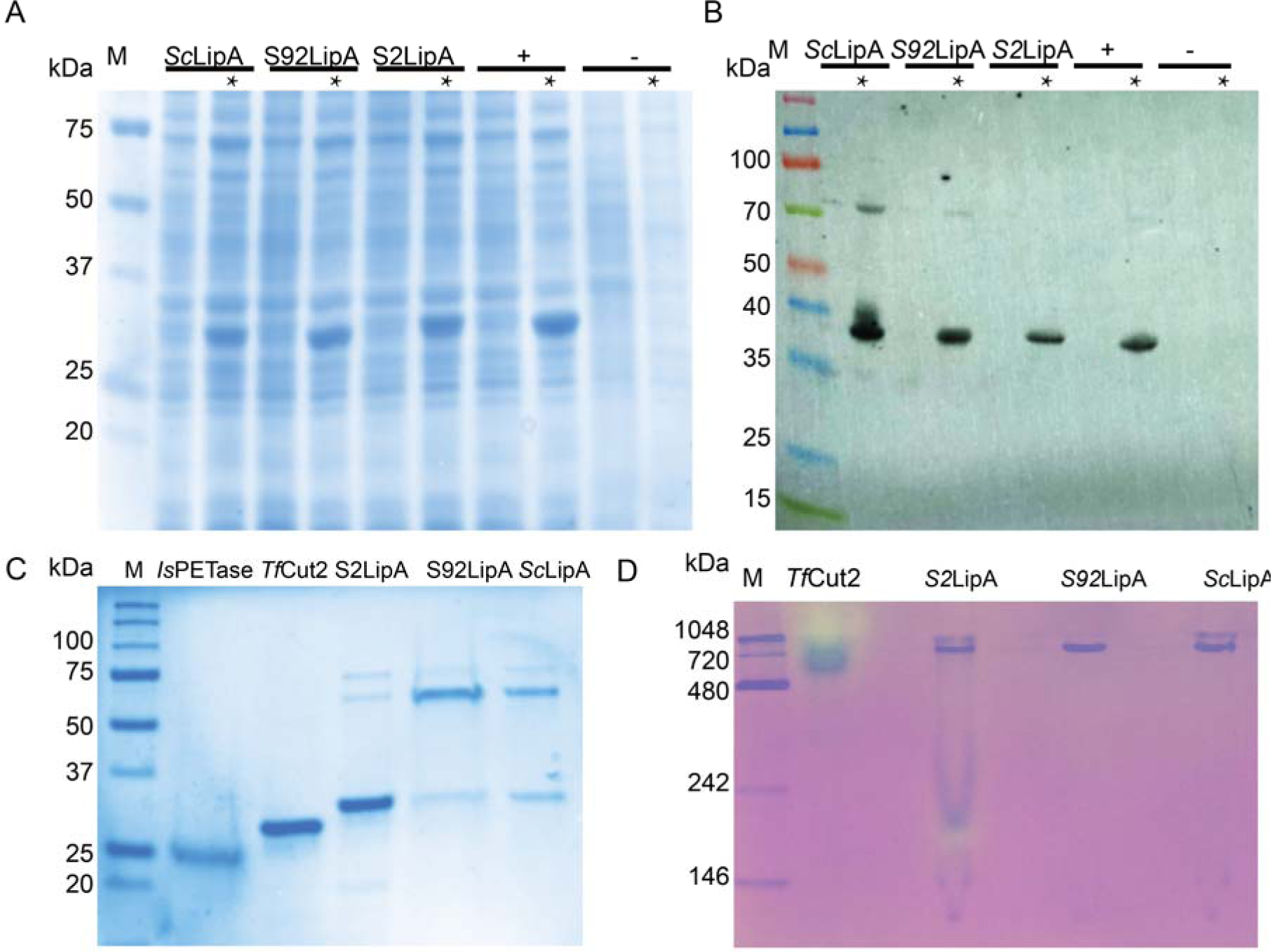
Expression and purification of Lipase A enzymes in E. coli. A); SDS-PAGE of uninduced and induced BL21 cultures. The Lipase A variants have a molecular weight of 28 kDa. The lanes with induced samples are indicated with an asterisk (*).The successful induction is evident in the induced samples, as indicated by the presence of thick bands around 28 kDa. 1: ladder, Biorad precision plus; 2&3: ScLipA; 4&5: S92LipA; 6&7: S2LipA; 8&9: Positive control pET16b with insert; 10&11: pET16b without insert. (B); Western blot was conducted to identify the induced proteins as the Lipase_A variants. In the induced samples clear black bands are visible around 38kDa, signifying the presence of the His-tagged Lipase A variants. 1: ladder, Spectra multicolor broad range protein marker; 2 till 11: same samples as for the SDS-PAGE. C); SDS-PAGE gel of the purified Lipase_A variants, along with the IsPETase and TfCut2; 1: Ladder, Biorad precision plus; 2: IsPETase sample; 3: TfCut2 sample; 4: S2LipA sample; 5: S92LipA; 6: ScLipA. Expected bands of the PETase is 25 kDa, TfCut2 (28 kDa), and the Lipase A variants 30 kDa. The Lipase variants show three bands one around 60 kDa and one around 70 kDa. D) Zymogram of TfCut2 and LipA variants on 1% tributyrin. TfCut2 and S2LipA show degradation of tributyrin (yellow spots) in one of the bands. Protein contents of other LipA variants it to low to observe activity. NativeMarker used as marker.

The optimal pH and temperature for enzyme activity were determined using paranitrophenyl dodecanoate as substrate according to the method of Altammar and colleagues [46]. A pH range from pH 3 to pH 8, was tested for one hour at 30 °C showing pH 7.0 as the optimal pH (Fig. 9A-C). A similar approach was taken to determine the optimal temperature. Enzymes were incubated using a temperature range of 25 °C-90 °C for 1 h. Thus, the optimal temperature appeared to be 25 °C (Fig. 9 A-C).

**Figure 9:**
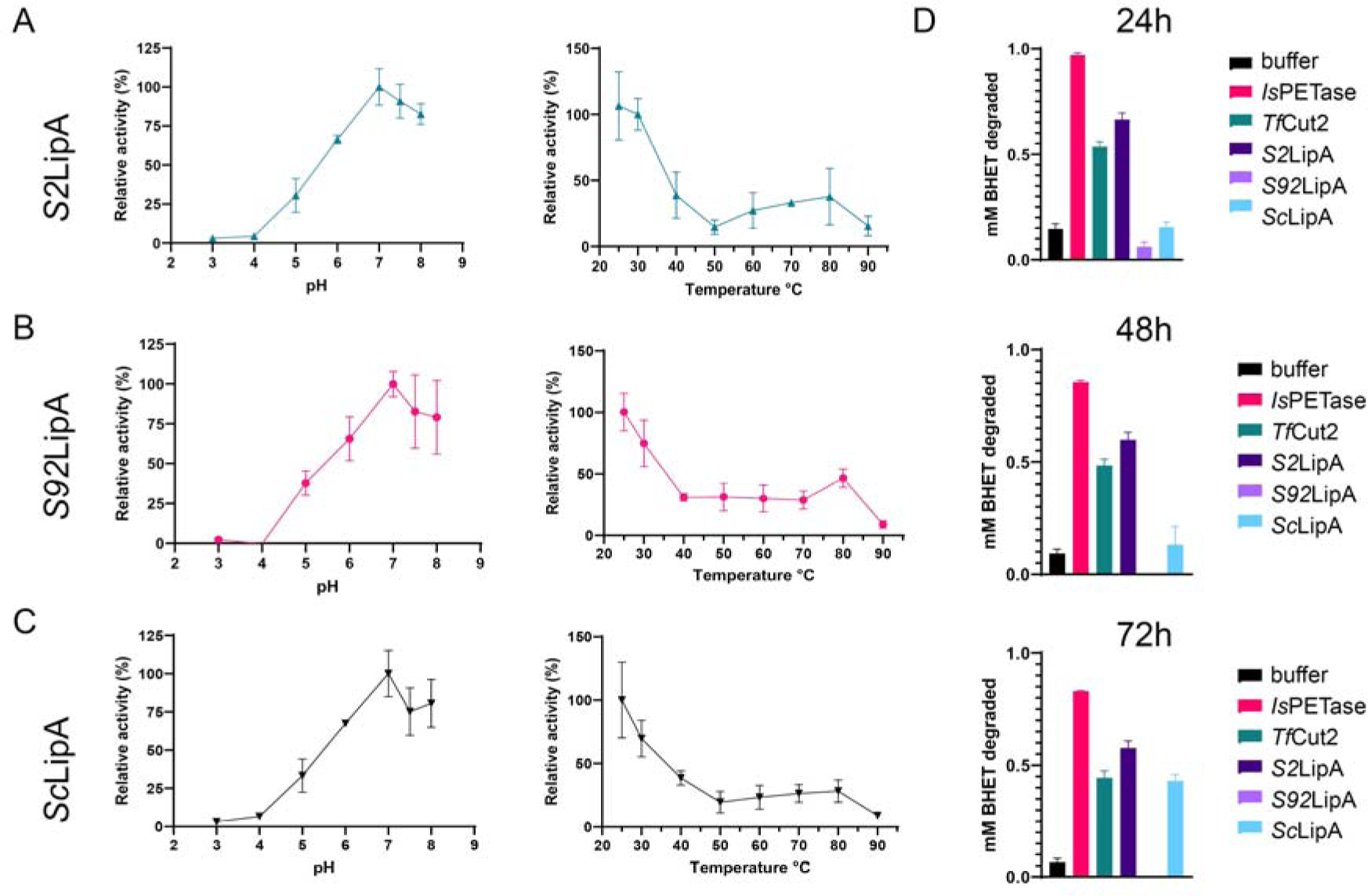
Determination of optimal enzyme conditions and BHET degradation under optimum conditions. A-C) relative activity on paranitrophenol dodecanoate at different pH for S2LipA (A), S92LipA(B) and ScLipA (C). D) Enzymatic BHET degradation using colorimetric assay after 24, 48 and 72 h. The activity of buffer control (black) as negative control, the IsPETase (magenta) and TfCut2 (turquoise) as positive controls and the samples S2LipA (dark purple,) S92LipA (lavender) and ScLipA (light blue) is displayed as the concentration BHET degraded in mM.

These conditions were then used to examine BHET degradation using a colorimetric assay. 2,5 ng/ml of enzyme was incubated with 1 mM of BHET for 24, 48 and 72 h. The positive controls *Is*PETase and *Tf*Cut2 show 50-100 % decrease in BHET (Fig. 9D). *S2*LipA, under these conditions, shows significantly more BHET degradation than *Tf*Cut2 after 24 h. *Sc*LipA shows delayed activity, after 72 h, BHET degrading activity is observed. The statistical analysis is provided in the supplement (S10). Simultaneously, amorphous PET films were incubated in pH 7.0 at 25 °C with 15 µg/ml of enzyme for 7 days and visualized using the SEM. *Is*PETase and *Tf*Cut2 were used as positive controls, whereas buffer without enzyme was used as negative control. The samples only incubated with buffer show a smooth clean surface. PET films incubated with the *Is*PETase show clear damage in the form of dents and cavities (Fig. 10). The *Tf*Cut2 showed roughening of the surface and at 10.000 x magnification small indents. *S2*LipA showed a similar pattern as *Tf*Cut2 only here it appeared that the top layer of the plastic was not completely degraded (2500 x-10.000 x), in places where this layer is degraded similar damage is observed as shown in *Tf*Cut2. Both *S92*LipA and *Sc*LipA show very limited to no activity (Fig. 10).

**Figure 10:**
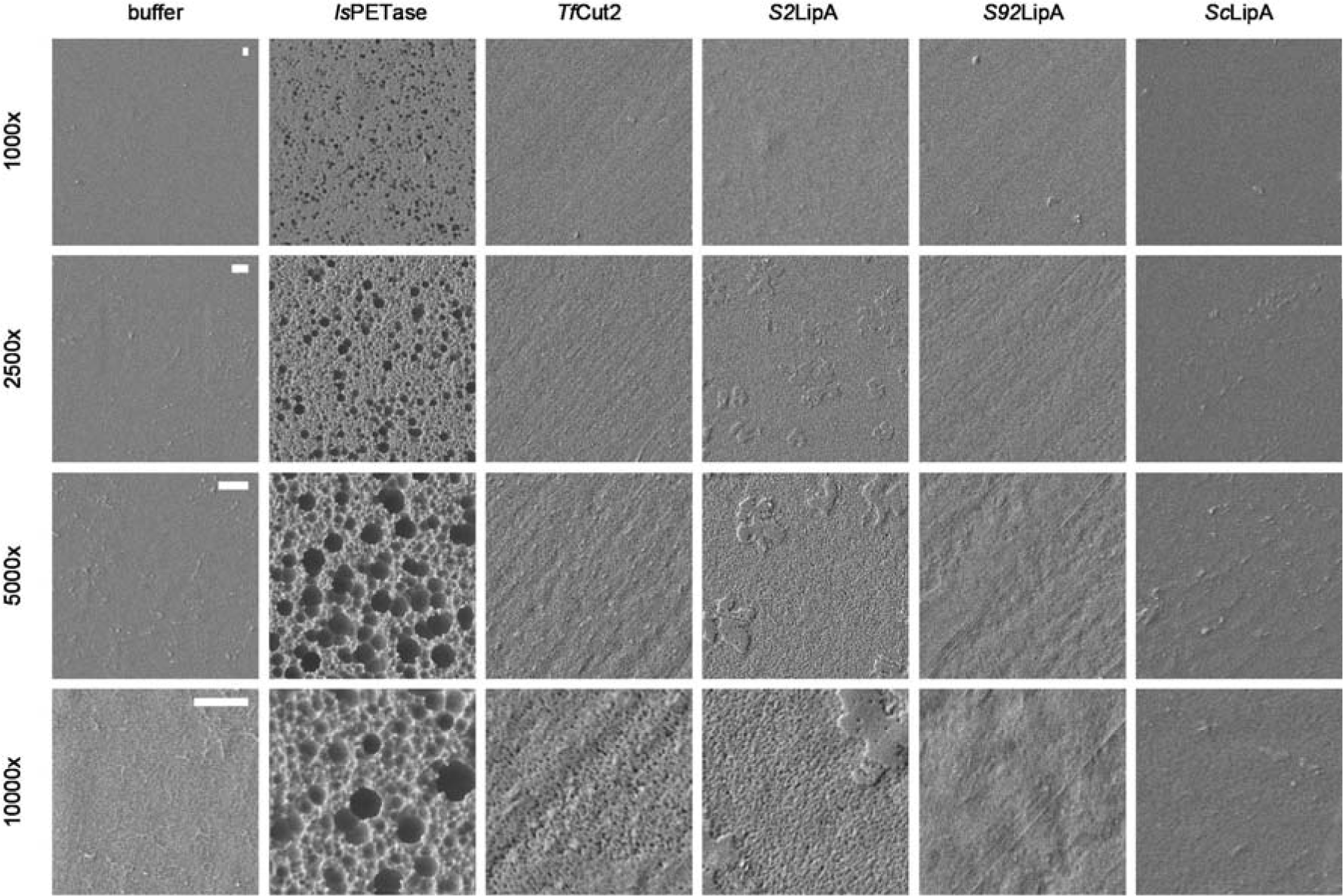
Effect of IsPETase, TfCut2, S2LipA, S92LipA and ScLipA on amorphous PET film. Amorphous PET films after 7 days of incubation with 15 µg/ml enzyme at 25 °C pH7 at 1000 – 10.000 x magnification. The scalebar is equal to 1 µm and applies to all images of the same magnification. Each row displays a different magnification whereas the columns show a different enzyme.

## Discussion

Plastics are very useful materials, simplifying everyday life. Unfortunately, poor waste management of plastics leads to polluting leakage into the environment. Understanding and harnessing nature’s response to this abundant plastics pollution may help find new solutions for sustainable plastics depolymerization and recycling. However, and importantly, the abundance of micro-organisms that have adapted to utilization and degradation of plastics is not known, and evolution and conservation of genes and proteins involved remains largely unexplored.

In this research, we have investigated plastics degrading activity of an *Is*PETase homolog Lipase A (LipA) from *Streptomyces coelicolor.* Of 96 *Streptomyces* strains 44 % was able to degrade BHET in at least one of the tested conditions, 18 % of the strains showed degrading activity when tested individually. Clearly, regulation of the induction of BHET-degrading activity by GlcNAc presented an important factor, however, the exact mechanism is still unknown. When strains MBT2, 5, 12, 38 and 92 were grown on amorphous PET films, they clearly showed adherence to the plastic while no immediate damage was observed.

We have shown the presence of *lipA* in 15 different Streptomyces strains as a highly conserved gene with some variations. The three variants *ScLipA* (present in *S. coelicolor*), S92LipA (present in MBT92 and S2LipA (present all other active strains) were further investigated. The *in vivo* activity of *Sc*LipA was investigated by comparing a knock-out strain with the wildtype strain, in both rich and minimal medium; the knock-out exhibited a significant decrease in its BHET degrading activity. Since some remaining BHET degradation was still observed in the knock-out strain, at least one other enzyme is likely to be involved in BHET degradation. However, this activity most likely resides with another type of enzyme since no other *Is*PETase homolog was found during the homology search. All three LipA variants, when expressed in the knock-out background under a semi-constitutive promoter, exhibited significantly enhanced degradation of BHET compared to the wildtype, on both rich and minimal medium. The variant enzymes were expressed in *E. coli*, purified and subjected to enzyme assays. Interestingly, after purification, two clear bands are visible at ∼30 kDa and ∼60 kDa. Native enzyme analysis via a zymogram indicated that activity only resided with the higher band, suggesting that the active form was a dimer. Indeed, 3D-structure prediction with AlphaFold-multimer suggested the formation of a dimeric enzyme with a probability of 90 % (S11) [47].

All enzyme variants were found to have an optimal pH of 7 and optimal temperature of 25 °C. Colorimetric BHET assays confirmed BHET degrading activity of *S2*LipA and *Sc*LipA under optimal conditions. Additionally, incubation of amorphous PET film with purified *S*2LipA caused the formation of indents and roughening of the surface, similar to the activity of PET-degrading cutinase *Tf*Cut2, further confirming PETase-like activity.

Our data suggests that PET is not the main substrate for these enzymes and that PET degradation merely appears as a moonlighting effect of the lipase activity, similar to the moonlighting effect described for *Tf*Cut2. *S92*LipA and *Sc*LipA showed less activity than *S2*LipA which most probably is due to structural variations in the enzymes caused by the various amino acid substitutions. For example, *S2*LipA contains a region rich in serines close to important substrate binding residues and the substrate binding cleft (Fig. 4I) as opposed to *S92*LipA and *Sc*LipA. This may result in enhanced formation of hydrogen bonds with the substrate and/or enhanced enzyme stability [48]–[50].

Additionally, serines have previously been found to make an enzyme more hydrophobic in neutral and alkaline conditions which is preferable for PET binding [51], [52]. Since all of the strains in our study are environmental isolates, we hypothesize that the abundance of *S2*LipA in these strains is initially caused by evolution towards a more stable and hydrophobic lipase and not toward plastic degradation per se. Yet, current pollution rates may push environmental evolution of those enzymes towards enhanced PET affinity. The current abundance of PET-degrading enzymes in nature indicates that nature can indeed take on the challenge of degradation and bioremediation. By further investigating and understanding this phenomenon, the microbial and enzyme characteristics involved, and applying those features, we may ultimately move towards less pollution and a more sustainable future.

## Materials and methods

### Strains and culturing conditions

For the initial tests, *Streptomyces coelicolor* M145 [28] and Actinobacteria from the MBT strain collection of the Institute of Biology Leiden were used for screening. These strains were isolated from the Himalaya mountains, Qinling mountains, Cheverny and Grenouillere France and the Netherlands [42]. The *Streptomyces* plates were incubated at 30 °C for 10 days or longer before investigating the BHET degradation. *E. coli* was grown on Luria-Burtani (LB) agar or in LB cultures overnight at 37 °C. Routine *Streptomyces* manipulation, growth and preparation of spore stocks was performed according to the *Streptomyces* manual [53]. Spore suspensions were obtained by growing the strains on Soy Flour Mannitol agar (SFM [54]) until sporulation.

### Homology search

Potential homologs of the PETase *Ideonella sakaiensis* (Uniprot A0A0K8P6T7 (PETH_IDESA)) in the Actinobacterial genomes were identified by performing a BLASTp for the genome of *Streptomyces coelicolor* (NCBI 100226) [35]. BLASTp was run in the default setting. The hits were ranked based on their BLAST score, their query cover, their percent identity and the presence of the catalytic triad and substrate binding site.

### Bulk screens

The strains are inoculated on the plates using the stamping method. A stamp fitting a 96-wells plate was autoclaved, placed in the stamping device and placed in the 96 wells plate, containing 100 μl of each spore stock in separate wells, and stamped on the plates of interest. Each spot contained approximately 2 μl of spore solution. The plates were grown for 10 days at 30 °C. the spore plates were stored at −20 °C.

### Plate assays

Plate assays were performed using minimal medium plates with different types of agars. Upon media selection, *Streptomyces* minimal medium (StrepMM [53]) Difco agar was chosen as standard condition [55]. Plates and additives are described in table 1. Bis(2-hydroxyethyl) terephthalate (BHET) provided by Sigma (Cas: 959-26-2, PN 465151), N-acetyl glucosamine (GlcNAc) provided by Sigma (Cas 7512-17-6, PN A4106).

**Table 1:** Plate compositions for screening.

| Experiment | Medium | Agar | Addition |
| --- | --- | --- | --- |
| Toxicity screen | StrepMM | Iberian Agar | Mannitol 25 mM +<br>BHET 0, 10 mM, 20<br>mM, 30 mM and 40 mM |
| Bulk screens | StrepMM | Iberian Agar<br>Difco agar<br>Agar-Agar | +/- BHET 10 mM<br>+/- GlcNAc 25 mM |
| Individual screens<br>*S. coelicolor* and active<br>strains | StrepMM | Difco Agar | +/- Mannitol 25 mM<br>+/- BHET 10 mM<br>+/- GlcNAc 25 mM |

### Individual screens

For the individual screens 2 μl of uncorrected spore solution was spotted on 5cm agar plates. And grown at 30 °C for 10 days.

### Agar excision agar samples

*S. coelicolor* M145, and *Streptomyces* species MBT2, MBT5, MBT12, MBT38 and MBT92 with 2*10^6^ spores were inoculated on StrepMM Difco with GlcNAc [25 mM] and BHET [10 mM]. After 14 days of growth, a part of the halo was excised using an agar excision tool. The agar and liquids were separated using clear spin-filtered microtubes (Sorenson BioSciences) spun at 10.000 rpm for 15 min. The flow-through was stored at −20 °C and prepared for LC-MS analysis according to the sample preparation protocol.

### Identification variants Strep strains

For the identification of the Lipase A variants in the *Actinobacteria* collection genomic DNA of the active strains was isolated. A PCR was performed using degenerative primers (Table 2) based on the sequence of the *2lipA* gene. PCR was performed with Phusion according to the Phusion protocol provide by Thermofischer scientific (F531S). The blunt-end PCR products were cloned in pJET2.1 using the CloneJET PCR cloning kit from Thermofischer scientific (K1231).

**Table 2:**
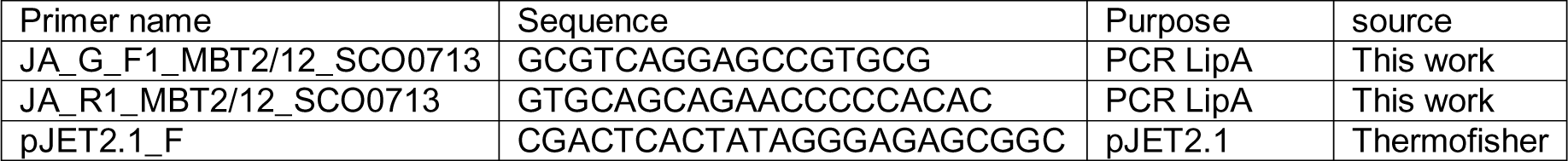
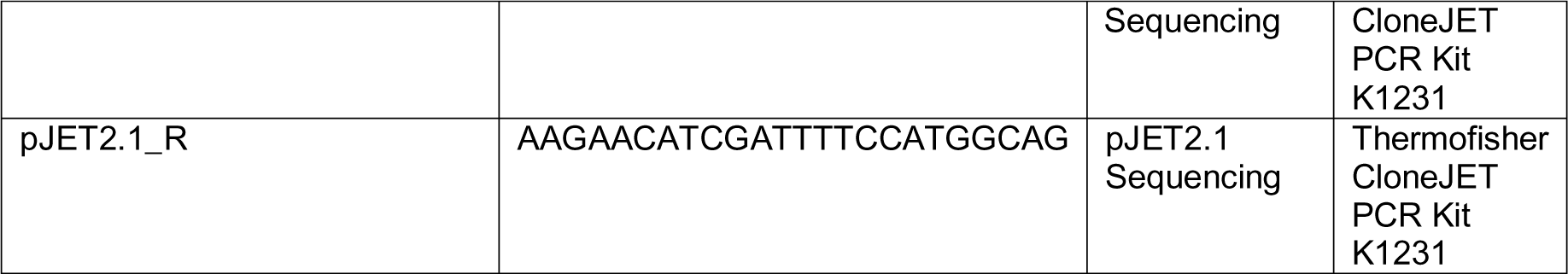
Primers for amplification of Lipase A variants.

**Table 3:**
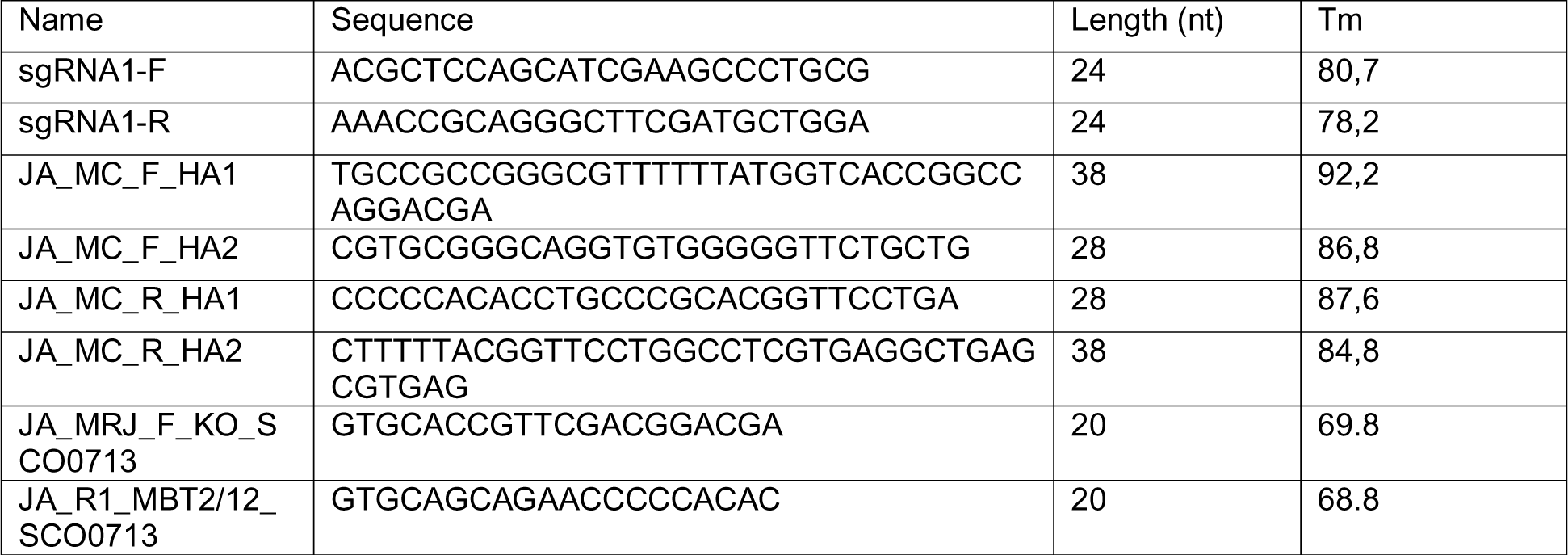
sgRNA and primers for homologous arms and diagnostic PCR.

### CRISPR/Cas knock-out

To obtain a LipA knock-out strain, the CRISPR-Cas9-based pCRISPomyces was used [56], [57]. In this system, the CRISPR-Cas9 mediated double stranded break will be repaired via homology-directed repair. Two homologous arms of approximately 1000 bp with a 40 nt overlap at both sides of *lipA* have been amplified via PCR on genomic DNA of M145. These homologous arms can be digested into the pCRISPomyces-2 via three-piece Gibson Assembly using the XbaI site. Additionally, a sgRNA will be annealed into the plasmid using Golden Gate cloning. Primers for these homologous arms bordering LipA and sgRNA were made using the protocol of Cobb and colleagues. Assembled plasmids were confirmed via sequencing. Assembled plasmids were transferred to *Streptomyces coelicolor* M145 via conjugation with *E. coli* ET12567 harboring the pUZ8002 plasmid [58], [59]. Single colonies were streaked on SFM with apramycin [50 μg/ml] to check for true apramycin resistance. Resistant colonies were picked to TSBS for gDNA isolation. Correct gene knock-outs were confirmed via diagnostic PCR and sequencing. Plasmid loss was achieved by restreaking strains for several generations on SFM agar without antibiotic pressure at 37 °C and checking for the loss of apramycin resistance [50], [51].

Overexpression strains were obtained by using the pSET152 integrative plasmid [60]. To create overexpression strains for LipA and its variants, the genes were transferred to a pSET152 integrative plasmid. Genomic DNA of *S. coelicolor* M145, *Streptomyces* sp. MBT2 and *Streptomyces* sp. MBT92 was isolated and the *lipA* PCR amplified with overhangs containing a NdeI site on the 5’ and a His-tag followed by a BamHI site on the 3’ end of the product. pSET152 was linearized with BamHI and NdeI and PCR amplified *lipA* genes were digested into the plasmid using T4 ligase. Assembled plasmids were confirmed via sequencing. *Sc*LipA and its variations were cloned into pSET152 under the regulation of strong semi-constitutive promotor pGAP [45]. pSET152 was transferred to *Streptomyces coelicolor* M145 via conjugation as described in [58], [59]. Single colonies were streaked on SFM with apramycin to select for correct integration of the plasmid. All strains obtained are described in table 4.

**Table 4:**
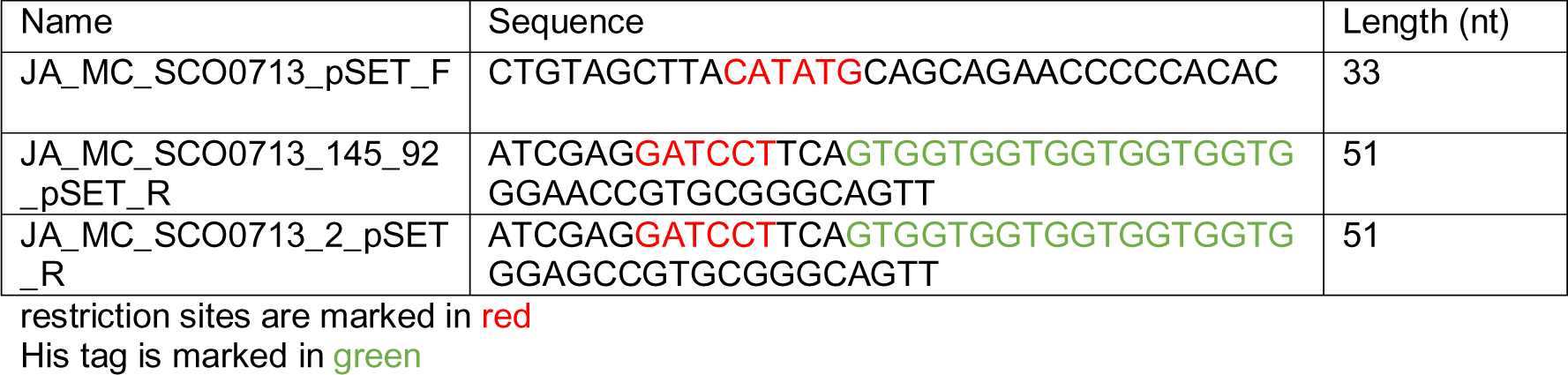
Primers amplification lipA variants for expression in pSET152.

**Table 5:** Obtained Streptomyces strains.

| Organism | Construct | Function | Working name |
| --- | --- | --- | --- |
| *S. coelicolor* M145 | - | WT | WT |
| *S. coelicolor* M145<br>*ΔlipA* | - | Knock-out | KO |
| *S. coelicolor* M145<br>*ΔlipA* | pSET152_pGAP_*lipA*<br>M145 | Overexpression | S3 |
| *S. coelicolor* M145<br>*ΔlipA* | pSET152_pGAP_*lipA*<br>MBT2 | Overexpression | S5 |
| *S. coelicolor* M145<br>*ΔlipA* | pSET152_pGAP_*lipA*<br>MBT92 | Overexpression | S7 |

### Liquid cultures for enzyme expression

For liquid chromatography-mass spectrometry (LC-MS) analysis, 1,0 x 10^8^ spores were inoculated in 50 ml liquid minimal medium without PEG (NMM [53]) + 5 % glucose (w/v) and Tryptic soy broth sucrose (TSBS [53]) containing 10 mM BHET. Samples of 1 ml were taken after 24, 48 and 72 h.

### Liquid chromatography-mass spectrometry (LC-MS)

Upon thawing, cell-free media was prepared for the LC-MS run by dilution in 1:1 in acetonitrile at a final volume of 1 ml and passed through 0.2 μm filters (Sartorius). Analysis of TPA, MHET, and BHET. LC-MS analyses were performed on a Shimadzu LC-20AD system with a Shimadzu Shim-Pack GIST-HP C18-AQ column (3.0 x 150 mm, 3 μm) at 40 °C and equipped with a UV detector monitoring at 240 and 260 nm. The following solvent system, at a flow rate of 0.5 ml/min, was used: solvent A, 0.1 % formic acid in water; solvent B, acetonitrile. Gradient elution was as follows: 80:20 (A/B) for 1 min, 80:20 to 45:55 (A/B) over 6 min, 45:55 to 0:100 (A/B) over 1 min, 0:100 (A/B) for 2 min, then reversion back to 80:20 (A/B) over 1 min and 80:20 (A/B) for 2 min. This system was connected to a Shimadzu 8040 triple quadrupole mass spectrometer (ESI ionisation).

The reference chromatogram used for the identification of BHET and its monomers [∼1 mg/ml each]. The area percentage was calculated using GraphPad Prism. The statistical analysis consists of a two-way ANOVA using the default setting of GraphPad Prism (p=0,05). A multiple comparison was performed providing insights into the significance of each compound present in each culture. The outcomes of the statistical analysis are provided in Supplement S8.

### Microscopy

The Lionheart FX automated microscope (BioTek) was used for the time-lapse imaging of BHET particle degradation. In total, 3.5*10^4^ spores were precultured in TSBS + nalidixic acid [50 µg/ml] for 48 h. Spores were precultured in TSBS containing nalidixic acid [50 µg/ml] for 48 h. Of this solution, 5 µl was inoculated in 200 µl TSBS containing ampicillin [50 µg/ml] and BHET [0.25 mM]. This solution was added to a Greiner Bio-One SensoPlate 96-well, non-treated black plate with a clear bottom and spun down to settle the mixture. Bright-field pictures were taken every 15 min over the course of 60 h on a 40 x magnification at 30 °C.

### Expression in *E. coli*

The used *S2*LipA, *S92*LipA and *Sc*LipA were codon optimized for *E. coli* and ordered with GeneART from ThermoFisher Scientific (Netherlands). The genes arrived in the Thermofisher plasmid pMA containing an ampicillin resistance.

The plasmids were transformed to chemical competent *E. coli DH5*α cells for amplification and isolated using the “GeneJET plasmid miniprep Kit" (Thermofischer, K0503). After plasmid isolation, the *lipA* variants were cloned into the pET16b plasmid using restriction/ligation with restriction enzymes BamHI and NdeI [10 U/ μl] (Thermofischer Scientific). BamHI was added half an hour after incubation with NdeI to achieve higher efficiency. Following restriction, the restriction enzymes were inactivated by incubating the samples for 20 min at 80 °C in a water bath. The *lipA* genes and the linearized pET16b plasmids were extracted from gel using the “GeneJET GEL extraction Kit” (Thermofischer, K0692). T4 ligase was used according to the user manual of the manufacturer. See Table 6 for an overview of the used plasmids in this study.

**Table 6:** Plasmids for expression Lip A variants in E. coli.

| Plasmids | Characteristics |
| --- | --- |
| pMA_LipA_MBT2 | Amp<sup>R</sup>, LipA_MBT2 |
| pMA_LipA_MBT92 | Amp<sup>R</sup>, LipA_MBT92 |
| pMA_LipA_M145 | Amp<sup>R</sup>, LipA_M145 |
| pET16b_LipA_MBT2 | Amp<sup>R</sup>, His-tag, LipA_MBT2 |
| pET16b_LipA_MBT92 | Amp<sup>R</sup>, His-tag, LipA_MBT92 |
| pET16b_LipA_M145 | amp<sup>r</sup>, His-tag, LipA_M145 |

**Table 7:** E. coli strains used in this study.

| Strain | Plasmid | Purpose |
| --- | --- | --- |
| *E. coli* BL21 | pET16b | Negative control |
|  | pET16b_PETase | Positive control<br>PETase activity |
|  | pET16b_TfCut2 | Positive control<br>Cutinase activity |
|  | pET16b_LipA M145 |  |
|  | pET16b_LipA MBT2 |  |
|  | pET16b_LipA MBT92 |  |

Transformed *E. coli BL21 A-I (*Invitrogen C607003) strains were grown on agar plates with ampicillin [100 μg/mL] at 37 °C or in LB liquid medium at 37 °C on 200 rpm.

### Enzyme purification

For protein production, 20 ml precultures with ampicillin (100 µg/ml) of the transformed *E. coli* BL21 A-I were made following the standard culture conditions. 5 ml of these precultures was transferred to 500 ml TB containing ampicillin. When an OD value between 0.6-0.8 was reached, the cultures were induced using of arabinose [0.2 % (w/v)] and IPTG [0.5 mM]. The induced cultures were incubated overnight along with a 1 ml non-induced sample at 16 °C with 200 rpm shaking.

The pellet was harvested from 1 ml of the induced and non-induced samples, by spinning down for 5 min at 13000 rpm and removing the supernatant. Subsequently, this pellet was resuspended in the 300 μl of MilliQ water. The samples were sonicated (Bandelin, Sonopuls) for 2x 30 sec at 15 % Amplitude with the MS73 probe to release to proteins. From the cell lysate, 3 μl was loaded on SDS-PAGE to evaluate if the induction was successful.

Pellet the 500 ml cultures were harvested by ultracentrifugation (Himac) for 30 min, 6000 rpm at 4 °C. The harvested pellet was transferred to 50 ml falcon tubes and frozen in liquid nitrogen. The frozen pellets were stored at −20 °C.

Next, defrosted pellet was resuspended in 15 ml Buffer A (table 8). These solutions were sonicated (Bandelin, Sonopuls) 3 times for 30 sec at 15 % Amplitude. Subsequently, to remove the insoluble faction’s ultracentrifugation (Himac) took place for 60 min, 10.000 rpm at 4 °C. The Sample was filtered through a 0.2 µm filter (Filtropur S 0.2 µm, Sarstedt).

**Table 8:** Buffers for the His – affinity purification.

| Buffer | Type of Buffer | Composition |
| --- | --- | --- |
| Buffer A | Equilibration buffer | 25mM Tris-HCl and 300 mM NaCl pH 7.5 |
| Buffer B | Elution Buffer | 25mM Tris-HCl, 300 mM NaCl and 500 mM imidazole pH 7.5 |
| Buffer C | Desalting Buffer | 25 mM Tris-HCl and 150 mM NaCl pH 7.5 |

To obtain purified Lipase A variants, His – affinity purification on an AKTA system was performed, following the AKTA start manual. In the purification process, two additional buffers were employed (Tabel 8).

During the purification, buffer B was used to first wash the non-binding proteins (15 % buffer B) and then to elute the Lipase A variants from the column (100 % buffer B). After purification, the fractions were desalted using buffer C desalting columns (Cytiva, PD-10) according to the protocol of Cytiva columns. The resulting purified Lipase A variants were stored at −20 °C in a 1:1 mixture of Buffer with 40 % glycerol.

### SDS-PAGE, Western blot and Zymogram

For SDS and Western-Blot, 50 ml of TSBS / NMM + 5 % glucose (v/v) were inoculated with 50 µl dense spore prep and incubated for 24 h. Cultures were centrifuged at 4000 g for 10 min and supernatant was filtered through a 0.2 μm filter. The supernatant was concentrated using a Viaspin column (Sartorius 5 kDas). Protein concentrations were estimated and normalized to 200 ng/μl by performing a Bradford assay.

Overall, all SDS pages contained 12 % acrylamide and were run for 20 min at 70 V to stack the proteins on the gels. Further, the gel was run at 150 V until the loading dye reached the bottom of the gel. SDS-page gels were stained with Coomassie-blue staining. For western blot, the gels were transferred using a BioRad Trans-blot Turbo and the corresponding transfer packs (1704157EDU) according to the mixed gel protocol of BioRad. Gel was washed using Tris buffered saline (TBS) buffer and blocked using Tris buffered saline with 0.5 % Tween 20 (TBST) buffer containing 1 % Elk milk. The blot was blocked between 60-90 min. His-antibody was added to a final concentration of 1 μg/ml and incubated overnight (K953-01). The blot was rinsed with water and washed 4 times with TBST, the blot is then incubated with luminol for 1 min, (product number) dried, and developed on X-ray film.

For the zymogram, an 8 % native gel, was run for 30 min at 70 V and 2 h at 150 V. The gel was rinsed with demi water and washed with demi water and equilibrated by incubating 3 times for 30 min with 25 mM NaCl. After, the gel is placed on a 1 % tributyrin, 0.5 % agarose plate and incubated at 30 °C for 1.5 h [61].

### Enzyme assays

#### Standard enzyme assays

The concentration of enzyme was estimated using the Bradford method (ref and company). The esterase/cutinase activity was tested using para-nitrophenyl dodecanoate (sigma) The protocol followed was based on the protocol of Altammar and colleagues with minor adjustments [46]. For the optimal pH test, 50 mM citrate buffers ranging from pH 3 to pH 7 were used, for pH 7.5 and 8 50 mM Tris-HCl buffer was used. The incubation step of 10 min was prolonged to 1 h. The reaction was terminated using 0.1 M sodium carbonate [46]

### Colorimetric assay BHET degradation

The colorimetric assay was performed according to the methods of Beech and colleagues [62]. 0.5 ng of enzyme was incubated with 1mM of BHET for 24, 48 and 72 h and measured at 615 nm in the Tecan M Spark. A reference line was made by adding BHET in the concentrations of 1 mM to 0 mM in steps of 0.1 mM an excess of *Tf*Cut2 was added to convert all BHET to MHET.

The amount of BHET degraded was calculated with GraphPad using the above-mentioned reference. The statistical analysis consists of a one-way ANOVA using the default setting of GraphPad Prism (p = 0.05). A comparison was made between the means of each enzyme treatment providing insights into the significance of the BHET degrading activity of each enzyme. The outcomes of the statistical analysis are provided in Supplemental data S10.

### SEM

The effect of Streptomyces on amorphous plastic films was investigated by incubating 107 spores in 3 ml of NMM with amorphous PET for two weeks at 30 °C. The samples were fixed with 1.5 % glutaraldehyde (30 min). Subsequently, samples were dehydrated using series of increasing ethanol percentages (70 %, 80 %, 90 %, 96 % and 100 %, each step 30 minutes) and critical point dried (Baltec CPD-030). Hereafter the samples were coated with 10 nm Platinum palladium using a sputter coater, and directly imaged using a JEOL JSM6700F. When investigating enzymes on PET film, 15 µg/ml of enzyme was incubated for 7 days, the films were washed with water and 70 % ethanol, air dried, sputter coated with 10 nm Platinum palladium and visualized using a JEOL JSM6700F.

## Supporting information

all Supplemental data

S9_control

S9_dLipA

## Contributions

JAV: conceptualization, data acquisition (including LC-MS, SEM and Lionheart) data analysis, figure design and writing. MC: Data acquisition (including LC-MS), data analysis and writing; SL homology search, 3D-structure modeling, PyMOL analysis and figure design; AM: Data acquisition; CB: Data acquisition; PI: development LC-MS methods and acquisition LC-MS data; JW: development SEM methods and acquiring SEM images; MEC: development Lionheart methods and acquisition Lionheart videos and pictures. GvW providing *Streptomyces* strains and proofreading; AR: conceptualization, supervision and reviewing. JdW: conceptualization, supervision, writing and reviewing.

## Acknowledgements

We thank Lennart Schada von Borzyskowski for providing BL21 AI and the pET16b. Additionally, Chaoxian Bai for providing the CRISPR system for *Streptomyces* and his advice on using it. Erik Vijgenboom and Jean Richard Quant for providing pSET152pGAP. Kees van den Hondel for this introduction into the lipase substrates. Marco Blasioli for his background work on LipA. Mia Urem for suggesting GlcNAc as possible inducer. Somayah Elsayed for explaining the MZ-mine software. Nathaniel Martin for his advice and support on setting up the LC-MS methods. Finally, Davy de Witt for preparing all the media.

