## Supplementary material for "Finding needles in haystacks: identification of novel conserved PETase enzymes in *Streptomyces*": all Supplemental data

### Supplementary data

#### S1. Bulk screen

| Agar | StrepMM BHET | StrepMM +Glcnac | Difco | StrepMM BHET | StrepMM +Glcnac | IB | StrepMM BHET | StrepMM +Glcnac |
|---|---|---|---|---|---|---|---|---|
| MBT1 | + | + | | + | + | | - | + |
| MBT2 | + | + | | + | + | | - | + |
| MBT3 | + | + | | + | + | | - | + |
| MBT4 | - | - | | - | - | | - | - |
| MBT5 | - | + | | + | + | | + | + |
| MBT6 | - | - | | + | + | | - | + |
| MBT7 | - | - | | - | - | | - | - |
| MBT8 | - | - | | - | - | | - | + |
| MBT9 | - | - | | - | - | | - | + |
| MBT10 | ng | - | | - | - | | - | - |
| MBT11 | + | - | | - | - | | - | + |
| MBT12 | + | + | | + | + | | + | + |
| MBT13 | - | - | | - | - | | - | - |
| MBT14 | - | - | | - | - | | - | ng |
| MBT15 | - | + | | + | + | | + | + |
| MBT16 | - | - | | - | - | | - | - |
| MBT17 | - | - | | - | - | | - | - |
| MBT18 | + | + | | + | + | | + | + |
| MBT19 | - | - | | - | - | | - | - |
| MBT20 | - | - | | - | - | | - | ng |
| MBT21 | - | - | | - | - | | - | + |
| MBT22 | - | - | | - | - | | - | + |
| MBT23 | - | - | | - | - | | - | + |
| MBT24 | - | - | | - | - | | - | + |
| MBT25 | - | - | | - | - | | - | - |
| MBT26 | - | - | | - | - | | - | - |
| MBT27 | - | - | | - | - | | - | - |
| MBT28 | + | + | | + | + | | + | + |
| MBT29 | + | + | | + | + | | + | + |
| MBT30 | - | - | | - | - | | - | - |
| MBT31 | - | - | | - | ~ | | - | ~ |
| MBT32 | - | - | | - | - | | - | - |

| | | | | | | |
|---|---|---|---|---|---|---|
| MBT33 | + | ~ | + | ~ | - | + |
| MBT34 | - | - | - | - | - | + |
| MBT35 | - | - | - | - | - | - |
| MBT36 | ng | ng | ng | ng | ng | ng |
| MBT37 | - | - | - | - | - | - |
| MBT38 | + | + | + | + | - | + |
| MBT39 | + | + | + | + | - | + |
| MBT40 | - | - | - | - | - | - |
| MBT41 | - | - | - | - | - | ~ |
| MBT42 | - | - | - | - | - | - |
| MBT43 | - | - | - | - | - | ~ |
| MBT44 | - | - | - | - | - | - |
| MBT45 | + | - | + | - | - | - |
| MBT46 | + | - | + | - | - | - |
| MBT47 | + | + | + | + | + | + |
| MBT48 | - | - | - | - | - | - |
| MBT49 | - | - | - | - | - | - |
| MBT50 | ~ | + | - | + | - | + |
| MBT51 | ~ | - | - | - | - | - |
| MBT52 | - | - | - | - | - | - |
| MBT53 | - | - | - | - | - | - |
| MBT54 | - | - | - | - | - | - |
| MBT55 | - | - | - | - | - | - |
| MBT56 | - | - | - | - | - | - |
| MBT57 | - | - | - | - | - | - |
| MBT58 | - | - | - | - | - | - |
| MBT59 | - | - | - | - | - | - |
| MBT60 | - | - | - | - | - | - |
| MBT61 | - | - | - | - | - | - |
| MBT62 | + | + | + | + | + | + |
| MBT63 | - | - | ~ | - | - | - |
| MBT64 | ~ | - | + | - | - | - |
| MBT65 | + | + | + | + | - | - |
| MBT66 | + | - | + | - | - | - |
| MBT67 | - | - | - | - | - | - |
| MBT68 | ng | - | - | - | - | - |
| MBT69 | - | - | - | - | ng | ng |

| | | | | | | |
|---|---|---|---|---|---|---|
| MBT70 | + | + | + | + | + | + |
| MBT71 | - | - | - | + | - | - |
| MBT72 | - | - | - | - | - | - |
| MBT73 | - | - | - | - | - | - |
| MBT74 | - | ~ | - | - | - | - |
| MBT75 | - | - | - | ~ | - | - |
| MBT76 | ng | ng | ng | ng | ng | ng |
| MBT77 | - | - | ~ | - | - | - |
| MBT78 | - | - | - | - | - | - |
| MBT79 | + | + | - | ~ | - | ~ |
| MBT80 | + | - | ~ | - | - | - |
| MBT81 | - | + | ~ | - | - | - |
| MBT82 | - | - | - | ~ | - | - |
| MBT83 | ng | ng | ng | ng | ng | ng |
| MBT84 | ng | ng | ng | ng | ng | ng |
| MBT85 | + | + | ~ | + | ~ | ~ |
| MBT86 | ~ | ~ | ~ | ~ | - | - |
| MBT87 | - | - | - | - | - | - |
| MBT89 | + | + | + | + | + | + |
| MBT90 | - | - | - | - | - | - |
| MBT91 | + | + | + | + | + | + |
| MBT92 | + | + | + | + | + | + |
| MBT93 | + | + | + | + | ~ | ~ |
| MBT95 | ng | ng | ng | ng | ng | ng |
| MBT96 | - | - | - | - | - | - |

#### S2. Individual screen

| strain | StrepMM BHET | BHET+man | BHET+GlcNAc | BHET glcnac+man |
|---|---|---|---|---|
| 1 | - | + | + | + |
| 2 | - | + | + | + |
| 3 | + | + | + | + |
| 5 | + | + | + | + |
| 6 | - | - | - | - |
| 8 | - | - | - | - |
| 11 | - | - | - | - |
| 12 | + | + | + | + |

| | | | | |
|---|---|---|---|---|
| 13 | - | - | - | - |
| 15 | + | + | + | + |
| 18 | + | + | + | + |
| 21 | - | - | - | - |
| 25 | - | - | - | - |
| 28 | - | + | + | + |
| 29 | - | + | + | + |
| 31 | - | - | - | - |
| 33 | - | - | - | - |
| 37 | - | - | - | - |
| 38 | - | + | + | + |
| 39 | - | - | + | + |
| 45 | - | - | - | - |
| 47 | - | + | + | + |
| 50 | - | - | + | + |
| 62 | - | - | - | - |
| 65 | - | - | - | - |
| 70 | - | - | - | - |
| 71 | - | - | - | - |
| 76 | - | - | - | - |
| 79 | - | - | - | - |
| 80 | - | - | - | - |
| 81 | - | - | - | - |
| 85 | - | - | - | - |
| 86 | - | - | - | - |
| 89 | + | + | + | + |
| 91 | - | + | + | + |
| 92 | + | + | + | + |
| 93 | - | - | - | - |
| 96 | - | - | - | - |
| S.coelicolor | - | - | - | - |

after 14 days*

#### S3. Pictures of all individual (images)

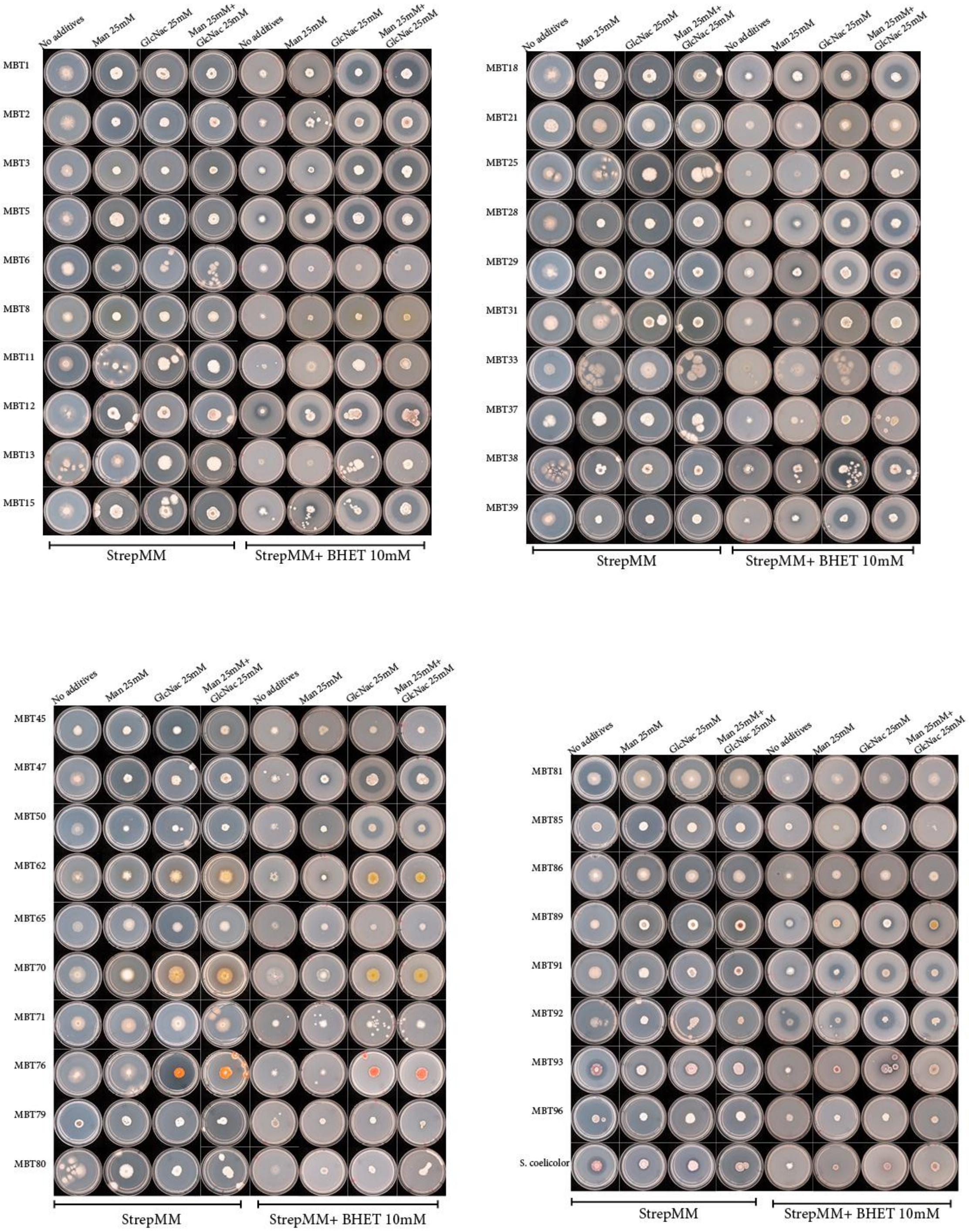

*Figure 1: Individual screen ofl active strains on Strep MM difco agar with and without Mannitol, BHET and GlcNac after 10 days of growth.*

#### S4. PCR & sequencing variants

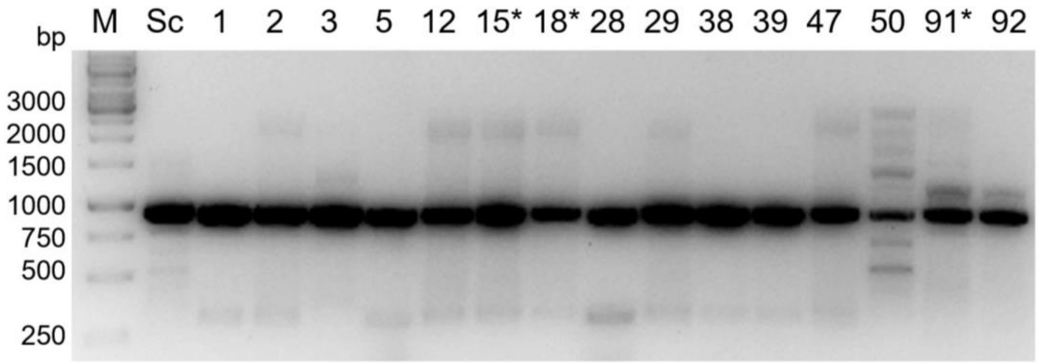

*Figure 2:* ***PCR of LipA gene in the active strains***

*Table 1:* ***Primers for PCR and sequencing LipA variants***

| Primer name | Sequence | Purpose | source |
|---|---|---|---|
| JA_G_F1_MBT2/12_SCO0713 | GCGTCAGGAGCCGTGCG | PCR LipA | This work |
| JA_R1_MBT2/12_SCO0713 | GTGCAGCAGAACCCCCACAC | PCR LipA | This work |
| pJET2.1_F | CGACTCACTATAGGGAGAGCGGC | pJET2.1 Sequencing | Thermofisher CloneJET PCR Kit K1231 |
| pJET2.1_R | AAGAACATCGATTTTCCATGGCAG | pJET2.1 Sequencing | Thermofisher CloneJET PCR Kit K1231 |

#### S5 DNA sequence alignment

| | | |
|---|---|---|
| MBT1 | GTGCAGCAGAACCCCCACAC------------CCACGCGGCGCGTCCGGCGTTCCGTGGA | 48 |
| MBT2 | GTGCAGCAGAACCCCCACAC------------CCACGCGGCGCGTCCGGCGTTCCGTGGA | 48 |
| MBT5 | GTGCAGCAGAACCCCCACAC------------CCACGCGGCGCGTCCGGCGTTCCGTGGA | 48 |
| MBT12 | GTGCAGCAGAACCCCCACAC------------CCACGCGGCGCGTCCGGCGTTCCGTGGA | 48 |
| MBT28 | GTGCAGCAGAACCCCCACAC------------CCACGCGGCGCGTCCGGCGTTCCGTGGA | 48 |
| MBT29 | GTGCAGCAGAACCCCCACAC------------CCACGCGGCGCGTCCGGCGTTCCGTGGA | 48 |
| MBT38 | GTGCAGCAGAACCCCCACAC------------CCACGCGGCGCGTCCGGCGTTCCGTGGA | 48 |
| MBT39 | GTGCAGCAGAACCCCCACAC------------CCACGCGGCGCGTCCGGCGTTCCGTGGA | 48 |
| MBT47 | GTGCAGCAGAACCCCCACAC------------CCACGCGGCGCGTCCGGCGTTCCGTGGA | 48 |
| MBT50 | GTGCAGCAGAACCCCCACAC------------CCACGCGGCGCGTCCGGCGTTCCGTGGA | 48 |
| MBT3 | GTGCAGCAGAACCCCCACAC------------CCACGCGGCGCGTCCGGCGTTCCGTGGA | 48 |
| M145 | GTGCAGCAGAACCCCCACACCCACGCCGCCCCAGGCGCCGCGCGCCCCGTCCTCCGGGGC | 60 |
| MBT92 | GTGCAGCAGAACCCCCACACCCACGCGGCCCCCGGGGCCGCGCGCCCCGTCTTCCGGGGC | 60 |

```
MBT1    CCCCGCCGGCGGCTCGCCGCTCTCACGGCCGCCGTGGCCGCCGCCGTCGCGCTCACCACC   108
MBT2    CCCCGCCGGCGGCTCGCCGCTCTCACGGCCGCCGTGGCCGCCGCCGTCGCGCTCACCACC   108
MBT5    CCCCGCCGGCGGCTCGCCGCTCTCACGGCCGCCGTGGCCGCCGCCGTCGCGCTCACCACC   108
MBT12   CCCCGCCGGCGGCTCGCCGCTCTCACGGCCGCCGTGGCCGCCGCCGTCGCGCTCACCACC   108
MBT28   CCCCGCCGGCGGCTCGCCGCTCTCACGGCCGCCGTGGCCGCCGCCGTCGCGCTCACCACC   108
MBT29   CCCCGCCGGCGGCTCGCCGCTCTCACGGCCGCCGTGGCCGCCGCCGTCGCGCTCACCACC   108
MBT38   CCCCGCCGGCGGCTCGCCGCTCTCACGGCCGCCGTGGCCGCCGCCGTCGCGCTCACCACC   108
MBT39   CCCCGCCGGCGGCTCGCCGCTCTCACGGCCGCCGTGGCCGCCGCCGTCGCGCTCACCACC   108
MBT47   CCCCGCCGGCGGCTCGCCGCTCTCACGGCCGCCGTGGCCGCCGCCGTCGCGCTCACCACC   108
MBT50   CCCCGCCGGCGGCTCGCCGCTCTCACGGCCGCCGTGGCCGCCGCCGTCGCGCTCACCACC   108
MBT3    CCCCGCCGGCGGCTCGCCGCTCTCACGGCCGCCGTGGCCGCCGCCGTCGCGCTCACCACC   108
M145    GTCCGGCGGCGGCTGGCCGCAGTGACGGCGGCCGTGGCCGCGGTCCTCGTGCTCGGCACC   120
MBT92   GTCCGGCGGCGGCTGGCCGGCGTGACGGCGGCCGTGGCCGCGGCCCTCGTGCTCGGCACC   120

MBT1    CTCACCGGCCCGGGCGCCCAGGCCGCCGACAACCCGTACGAGCGCGGCCCGGCGCCCACC   168
MBT2    CTCACCGGCCCGGGCGCCCAGGCCGCCGACAACCCGTACGAGCGCGGCCCGGCGCCCACC   168
MBT5    CTCACCGGCCCGGGCGCCCAGGCCGCCGACAACCCGTACGAGCGCGGCCCGGCGCCCACC   168
MBT12   CTCACCGGCCCGGGCGCCCAGGCCGCCGACAACCCGTACGAGCGCGGCCCGGCGCCCACC   168
MBT28   CTCACCGGCCCGGGCGCCCAGGCCGCCGACAACCCGTACGAGCGCGGCCCGGCGCCCACC   168
MBT29   CTCACCGGCCCGGGCGCCCAGGCCGCCGACAACCCGTACGAGCGCGGCCCGGCGCCCACC   168
MBT38   CTCACCGGCCCGGGCGCCCAGGCCGCCGACAACCCGTACGAGCGCGGCCCGGCGCCCACC   168
MBT39   CTCACCGGCCCGGGCGCCCAGGCCGCCGACAACCCGTACGAGCGCGGCCCGGCGCCCACC   168
MBT47   CTCACCGGCCCGGGCGCCCAGGCCGCCGACAACCCGTACGAGCGCGGCCCGGCGCCCACC   168
MBT50   CTCACCGGCCCGGGCGCCCAGGCCGCCGACAACCCGTACGAGCGCGGCCCGGCGCCCACC   168
MBT3    CTCACCGGCCCGGGCGCCCAGGCCGCCGACAACCCGTACGAGCGCGGCCCGGCGCCCACC   168
M145    CTCACCGGCCCGGGAGCCCAGGCCGCCGACAACCCGTACGAGCGCGGCCCGGCGCCCACC   180
MBT92   CTCACCGGCCCGGGCGCCCAGGCCGCGGACAACCCGTACGAGCGCGGCCCGGCGCCCACC   180

MBT1    GAGTCCAGCATCGAGGCCCTGCGCGGTCCGTACTCCGTGGCCGACACCAGCGTCTCCTCG   228
MBT2    GAGTCCAGCATCGAGGCCCTGCGCGGTCCGTACTCCGTGGCCGACACCAGCGTCTCCTCG   228
MBT5    GAGTCCAGCATCGAGGCCCTGCGCGGTCCGTACTCCGTGGCCGACACCAGCGTCTCCTCG   228
MBT12   GAGTCCAGCATCGAGGCCCTGCGCGGTCCGTACTCCGTGGCCGACACCAGCGTCTCCTCG   228
MBT28   GAGTCCAGCATCGAGGCCCTGCGCGGTCCGTACTCCGTGGCCGACACCAGCGTCTCCTCG   228
MBT29   GAGTCCAGCATCGAGGCCCTGCGCGGTCCGTACTCCGTGGCCGACACCAGCGTCTCCTCG   228
MBT38   GAGTCCAGCATCGAGGCCCTGCGCGGTCCGTACTCCGTGGCCGACACCAGCGTCTCCTCG   228
MBT39   GAGTCCAGCATCGAGGCCCTGCGCGGTCCGTACTCCGTGGCCGACACCAGCGTCTCCTCG   228
MBT47   GAGTCCAGCATCGAGGCCCTGCGCGGTCCGTACTCCGTGGCCGACACCAGCGTCTCCTCG   228
MBT50   GAGTCCAGCATCGAGGCCCTGCGCGGTCCGTACTCCGTGGCCGACACCAGCGTCTCCTCG   228
MBT3    GAGTCCAGCATCGAGGCCCTGCGCGGTCCGTACTCCGTGGCCGACACCAGCGTCTCCTCG   228
M145    GAGTCCAGCATCGAAGCCCTGCGCGGTCCCTACTCCGTGGCCGACACCAGCGTCTCCTCG   240
MBT92   GAGTCCAGCATCGAGGCCCTGCGCGGCCCCTACTCCGTGGCCGACACCAGGGTCTCCTCG   240

MBT1    CTCGCCGTCACCGGCTTCGGCGGCGGCACCATCTACTACCCGACCAGCACCAGCGACGGC   288
MBT2    CTCGCCGTCACCGGCTTCGGCGGCGGCACCATCTACTACCCGACCAGCACCAGCGACGGC   288
MBT5    CTCGCCGTCACCGGCTTCGGCGGCGGCACCATCTACTACCCGACCAGCACCAGCGACGGC   288
MBT12   CTCGCCGTCACCGGCTTCGGCGGCGGCACCATCTACTACCCGACCAGCACCAGCGACGGC   288
MBT28   CTCGCCGTCACCGGCTTCGGCGGCGGCACCATCTACTACCCGACCAGCACCAGCGACGGC   288
MBT29   CTCGCCGTCACCGGCTTCGGCGGCGGCACCATCTACTACCCGACCAGCACCAGCGACGGC   288
MBT38   CTCGCCGTCACCGGCTTCGGCGGCGGCACCATCTACTACCCGACCAGCACCAGCGACGGC   288
```

```
MBT39   CTCGCCGTCACCGGCTTCGGCGGCACCATCTACTACCCGACCAGCACCAGCGACGGC   288
MBT47   CTCGCCGTCACCGGCTTCGGCGGCACCATCTACTACCCGACCAGCACCAGCGACGGC   288
MBT50   CTCGCCGTCACCGGCTTCGGCGGCACCATCTACTACCCGACCAGCACCAGCGACGGC   288
MBT3    CTCGCCGTCACCGGCTTCGGCGGCACCATCTACTACCCGACCAGCACCAGCGACGGC   288
M145    CTCGCGGTCACCGGATTCGGCGGCACCATCTACTACCCGACCAGCACCAGCGACGGC   300
MBT92   CTCGCGGTCACCGGATTCGGCGGCACCATCTACTACCCGACCAGCACCAGCGACGGC   300

MBT1    ACCTTCGGCGCCGTGGTGATCTCGCCCGGCTTCACCGCGTACCAGTCGTCCATCGCCTGG   348
MBT2    ACCTTCGGCGCCGTGGTGATCTCGCCCGGCTTCACCGCGTACCAGTCGTCCATCGCCTGG   348
MBT5    ACCTTCGGCGCCGTGGTGATCTCGCCCGGCTTCACCGCGTACCAGTCGTCCATCGCCTGG   348
MBT12   ACCTTCGGCGCCGTGGTGATCTCGCCCGGCTTCACCGCGTACCAGTCGTCCATCGCCTGG   348
MBT28   ACCTTCGGCGCCGTGGTGATCTCGCCCGGCTTCACCGCGTACCAGTCGTCCATCGCCTGG   348
MBT29   ACCTTCGGCGCCGTGGTGATCTCGCCCGGCTTCACCGCGTACCAGTCGTCCATCGCCTGG   348
MBT38   ACCTTCGGCGCCGTGGTGATCTCGCCCGGCTTCACCGCGTACCAGTCGTCCATCGCCTGG   348
MBT39   ACCTTCGGCGCCGTGGTGATCTCGCCCGGCTTCACCGCGTACCAGTCGTCCATCGCCTGG   348
MBT47   ACCTTCGGCGCCGTGGTGATCTCGCCCGGCTTCACCGCGTACCAGTCGTCCATCGCCTGG   348
MBT50   ACCTTCGGCGCCGTGGTGATCTCGCCCGGCTTCACCGCGTACCAGTCGTCCATCGCCTGG   348
MBT3    ACCTTCGGCGCCGTGGTGATCTCGCCCGGCTTCACCGCGTACCAGTCGTCCATCGCCTGG   348
M145    ACGTTCGGCGCCGTCGTCATCGCGCCCGGGTTCACCGCGTACCAGTCGTCCATCGCCTGG   360
MBT92   ACGTTCGGCGCCGTGGCGATCGCACCCGGCTTCACCGCGTACCAGTCCTCCATGGCCTGG   360

MBT1    CTCGGTCCGCGGCTGGCCTCGCAGGGCTTCGTGGTCTTCACCATCGACACCAACACCACG   408
MBT2    CTCGGTCCGCGGCTGGCCTCGCAGGGCTTCGTGGTCTTCACCATCGACACCAACACCACG   408
MBT5    CTCGGTCCGCGGCTGGCCTCGCAGGGCTTCGTGGTCTTCACCATCGACACCAACACCACG   408
MBT12   CTCGGTCCGCGGCTGGCCTCGCAGGGCTTCGTGGTCTTCACCATCGACACCAACACCACG   408
MBT28   CTCGGTCCGCGGCTGGCCTCGCAGGGCTTCGTGGTCTTCACCATCGACACCAACACCACG   408
MBT29   CTCGGTCCGCGGCTGGCCTCGCAGGGCTTCGTGGTCTTCACCATCGACACCAACACCACG   408
MBT38   CTCGGTCCGCGGCTGGCCTCGCAGGGCTTCGTGGTCTTCACCATCGACACCAACACCACG   408
MBT39   CTCGGTCCGCGGCTGGCCTCGCAGGGCTTCGTGGTCTTCACCATCGACACCAACACCACG   408
MBT47   CTCGGTCCGCGGCTGGCCTCGCAGGGCTTCGTGGTCTTCACCATCGACACCAACACCACG   408
MBT50   CTCGGTCCGCGGCTGGCCTCGCAGGGCTTCGTGGTCTTCACCATCGACACCAACACCACG   408
MBT3    CTCGGTCCGCGGCTGGCCTCGCAGGGCTTCGTGGTCTTCACCATCGACACCAACACCACG   408
M145    CTCGGCCCGCGCCTGGCCTCCCAGGGCTTCGTGGTCTTCACCATCGACACCAACACCACG   420
MBT92   CTCGGTCCGCGGCTGGCCTCCCAGGGCTTCGTGGTCTTCACCATCGACACCAACACCACG   420

MBT1    CTGGACCAGCCCGACTCCCGAGGCCGGCAACTGCTGGCCGCCCTGGACTACCTGACCGAG   468
MBT2    CTGGACCAGCCCGACTCCCGAGGCCGGCAACTGCTGGCCGCCCTGGACTACCTGACCGAG   468
MBT5    CTGGACCAGCCCGACTCCCGAGGCCGGCAACTGCTGGCCGCCCTGGACTACCTGACCGAG   468
MBT12   CTGGACCAGCCCGACTCCCGAGGCCGGCAACTGCTGGCCGCCCTGGACTACCTGACCGAG   468
MBT28   CTGGACCAGCCCGACTCCCGAGGCCGGCAACTGCTGGCCGCCCTGGACTACCTGACCGAG   468
MBT29   CTGGACCAGCCCGACTCCCGAGGCCGGCAACTGCTGGCCGCCCTGGACTACCTGACCGAG   468
MBT38   CTGGACCAGCCCGACTCCCGAGGCCGGCAACTGCTGGCCGCCCTGGACTACCTGACCGAG   468
MBT39   CTGGACCAGCCCGACTCCCGAGGCCGGCAACTGCTGGCCGCCCTGGACTACCTGACCGAG   468
MBT47   CTGGACCAGCCCGACTCCCGAGGCCGGCAACTGCTGGCCGCCCTGGACTACCTGACCGAG   468
MBT50   CTGGACCAGCCCGACTCCCGAGGCCGGCAACTGCTGGCCGCCCTGGACTACCTGACCGAG   468
MBT3    CTGGACCAGCCCGACTCCCGAGGCCGGCAACTGCTGGCCGCCCTGGACTACCTGACCGAG   468
M145    CTGGACCAGCCCGACTCCCGCGGGCGGCAACTGCTGGCCGCCCTGGACTACCTGACCGGG   480
MBT92   CTGGACCAGCCCGACTCCCGCGGCCGCCAACTGCTGGCCGCCCTGGACTACCTCACCGAG   480
```

```
MBT1    CGCAGCTCCGTCCGGGGACGGATCGACAGCAGCCGGCTCGGCGTCATGGGCCACTCCATG   528
MBT2    CGCAGCTCCGTCCGGGGACGGATCGACAGCAGCCGGCTCGGCGTCATGGGCCACTCCATG   528
MBT5    CGCAGCTCCGTCCGGGGACGGATCGACAGCAGCCGGCTCGGCGTCATGGGCCACTCCATG   528
MBT12   CGCAGCTCCGTCCGGGGACGGATCGACAGCAGCCGGCTCGGCGTCATGGGCCACTCCATG   528
MBT28   CGCAGCTCCGTCCGGGGACGGATCGACAGCAGCCGGCTCGGCGTCATGGGCCACTCCATG   528
MBT29   CGCAGCTCCGTCCGGGGACGGATCGACAGCAGCCGGCTCGGCGTCATGGGCCACTCCATG   528
MBT38   CGCAGCTCCGTCCGGGGACGGATCGACAGCAGCCGGCTCGGCGTCATGGGCCACTCCATG   528
MBT39   CGCAGCTCCGTCCGGGGACGGATCGACAGCAGCCGGCTCGGCGTCATGGGCCACTCCATG   528
MBT47   CGCAGCTCCGTCCGGGGACGGATCGACAGCAGCCGGCTCGGCGTCATGGGCCACTCCATG   528
MBT50   CGCAGCTCCGTCCGGGGACGGATCGACAGCAGCCGGCTCGGCGTCATGGGCCACTCCATG   528
MBT3    CGCAGCTCCGTCCGGGGACGGATCGACAGCAGCCGGCTCGGCGTCATGGGCCACTCCATG   528
M145    CGCAGCTCCGTCCGCGGCCGGATCGACAGCGGCCGACTCGGCGTCATGGGCCACTCCATG   540
MBT92   CGCAGCTCCGTCCGCGGCCGGATCGACAGCAGCCGCCTCGGCGTGATGGGCCACTCCATG   540

MBT1    GGCGGCGGAGGCTCGCTGGAGGCCGCCAAGTCCCGTCCGTCGCTCCAGGCGGCGATCCCG   588
MBT2    GGCGGCGGAGGCTCGCTGGAGGCCGCCAAGTCCCGTCCGTCGCTCCAGGCGGCGATCCCG   588
MBT5    GGCGGCGGAGGCTCGCTGGAGGCCGCCAAGTCCCGTCCGTCGCTCCAGGCGGCGATCCCG   588
MBT12   GGCGGCGGAGGCTCGCTGGAGGCCGCCAAGTCCCGTCCGTCGCTCCAGGCGGCGATCCCG   588
MBT28   GGCGGCGGAGGCTCGCTGGAGGCCGCCAAGTCCCGTCCGTCGCTCCAGGCGGCGATCCCG   588
MBT29   GGCGGCGGAGGCTCGCTGGAGGCCGCCAAGTCCCGTCCGTCGCTCCAGGCGGCGATCCCG   588
MBT38   GGCGGCGGAGGCTCGCTGGAGGCCGCCAAGTCCCGTCCGTCGCTCCAGGCGGCGATCCCG   588
MBT39   GGCGGCGGAGGCTCGCTGGAGGCCGCCAAGTCCCGTCCGTCGCTCCAGGCGGCGATCCCG   588
MBT47   GGCGGCGGAGGCTCGCTGGAGGCCGCCAAGTCCCGTCCGTCGCTCCAGGCGGCGATCCCG   588
MBT50   GGCGGCGGAGGCTCGCTGGAGGCCGCCAAGTCCCGTCCGTCGCTCCAGGCGGCGATCCCG   588
MBT3    GGCGGCGGAGGCTCGCTGGAGGCCGCCAAGTCCCGTCCGTCGCTCCAGGCGGCGATCCCG   588
M145    GGCGGCGGCGGCACCCTGGAGGCCGCCAAGTCCCGGCCGTCGCTCCAGGCGGCGATCCCC   600
MBT92   GGCGGCGGCGGCACTTTGGAGGCCGCCAAGTCCCGGCCGTCCCTCCAGGCGGCGATCCCC   600

MBT1    CTCACCCCCTGGAACCTGGACAAGAGCTGGCCGGAGGTCAGCACCCCGACCCTGATCGTG   648
MBT2    CTCACCCCCTGGAACCTGGACAAGAGCTGGCCGGAGGTCAGCACCCCGACCCTGATCGTG   648
MBT5    CTCACCCCCTGGAACCTGGACAAGAGCTGGCCGGAGGTCAGCACCCCGACCCTGATCGTG   648
MBT12   CTCACCCCCTGGAACCTGGACAAGAGCTGGCCGGAGGTCAGCACCCCGACCCTGATCGTG   648
MBT28   CTCACCCCCTGGAACCTGGACAAGAGCTGGCCGGAGGTCAGCACCCCGACCCTGATCGTG   648
MBT29   CTCACCCCCTGGAACCTGGACAAGAGCTGGCCGGAGGTCAGCACCCCGACCCTGATCGTG   648
MBT38   CTCACCCCCTGGAACCTGGACAAGAGCTGGCCGGAGGTCAGCACCCCGACCCTGATCGTG   648
MBT39   CTCACCCCCTGGAACCTGGACAAGAGCTGGCCGGAGGTCAGCACCCCGACCCTGATCGTG   648
MBT47   CTCACCCCCTGGAACCTGGACAAGAGCTGGCCGGAGGTCAGCACCCCGACCCTGATCGTG   648
MBT50   CTCACCCCCTGGAACCTGGACAAGAGCTGGCCGGAGGTCAGCACCCCGACCCTGATCGTG   648
MBT3    CTCACCCCCTGGAACCTGGACAAGAGCTGGCCGGAGGTCAGCACCCCGACCCTGATCGTG   648
M145    CTCACCCCCTGGAACCTGGACAAGAGCTGGCCGGAGGTCAGCACGCCGACGCTGGTCGTC   660
MBT92   CTCACGCCCTGGAACCTGGACAAGAGCTGGCCCGAGGTCAGCACCCCGACGCTGATCGTG   660

MBT1    GGGGCCGACGGCGACACGGTCGCGCCCGTCTCCTCGCACTCCGAGCCTTTCTACTCCAGC   708
MBT2    GGGGCCGACGGCGACACGGTCGCGCCCGTCTCCTCGCACTCCGAGCCTTTCTACTCCAGC   708
MBT5    GGGGCCGACGGCGACACGGTCGCGCCCGTCTCCTCGCACTCCGAGCCTTTCTACTCCAGC   708
MBT12   GGGGCCGACGGCGACACGGTCGCGCCCGTCTCCTCGCACTCCGAGCCTTTCTACTCCAGC   708
MBT28   GGGGCCGACGGCGACACGGTCGCGCCCGTCTCCTCGCACTCCGAGCCTTTCTACTCCAGC   708
MBT29   GGGGCCGACGGCGACACGGTCGCGCCCGTCTCCTCGCACTCCGAGCCTTTCTACTCCAGC   708
```

| | | |
|---|---|---|
| MBT38 | GGGGCCGACGGCGACACGGTCGCGCCCGTCTCCTCGCACTCCGAGCCTTTCTACTCCAGC | 708 |
| MBT39 | GGGGCCGACGGCGACACGGTCGCGCCCGTCTCCTCGCACTCCGAGCCTTTCTACTCCAGC | 708 |
| MBT47 | GGGGCCGACGGCGACACGGTCGCGCCCGTCTCCTCGCACTCCGAGCCTTTCTACTCCAGC | 708 |
| MBT50 | GGGGCCGACGGCGACACGGTCGCGCCCGTCTCCTCGCACTCCGAGCCTTTCTACTCCAGC | 708 |
| MBT3 | GGTGCCGACGGCGACACGGTCGCGCCCGTCTCCTCGCACTCCGAGCCGTTCTACTCCAGC | 708 |
| M145 | GGGGCCGACGGCGACACGATCGCCCCCGTGGCCTCGCACGCCGAACCGTTCTACTCCGGC | 720 |
| MBT92 | GGGGCCGACGGCGACACGATCGCCCCCGTGGCCTCGCACGCCGAGCCGTTCTACTCCGGC | 720 |
| MBT1 | CTGCCGTCCGGAACGGACCGCGCCTACCTGGAGCTGAACAACGCGACCCACTTCTCGCCG | 768 |
| MBT2 | CTGCCGTCCGGAACGGACCGCGCCTACCTGGAGCTGAACAACGCGACCCACTTCTCGCCG | 768 |
| MBT5 | CTGCCGTCCGGAACGGACCGCGCCTACCTGGAGCTGAACAACGCGACCCACTTCTCGCCG | 768 |
| MBT12 | CTGCCGTCCGGAACGGACCGCGCCTACCTGGAGCTGAACAACGCGACCCACTTCTCGCCG | 768 |
| MBT28 | CTGCCGTCCGGAACGGACCGCGCCTACCTGGAGCTGAACAACGCGACCCACTTCTCGCCG | 768 |
| MBT29 | CTGCCGTCCGGAACGGACCGCGCCTACCTGGAGCTGAACAACGCGACCCACTTCTCGCCG | 768 |
| MBT38 | CTGCCGTCCGGAACGGACCGCGCCTACCTGGAGCTGAACAACGCGACCCACTTCTCGCCG | 768 |
| MBT39 | CTGCCGTCCGGAACGGACCGCGCCTACCTGGAGCTGAACAACGCGACCCACTTCTCGCCG | 768 |
| MBT47 | CTGCCGTCCGGAACGGACCGCGCCTACCTGGAGCTGAACAACGCGACCCACTTCTCGCCG | 768 |
| MBT50 | CTGCCGTCCGGAACGGACCGCGCCTACCTGGAGCTGAACAACGCGACCCACTTCTCGCCG | 768 |
| MBT3 | CTGCCGTCCGGAACGGACCGCGCCTACCTGGAGCTGAACAACGCGACCCACTTCTCGCCG | 768 |
| M145 | CTGCCCTCGTCGACCGACCGGGCCTATCTGGAGCTGAACAACGCGACCCACTTCTCGCCC | 780 |
| MBT92 | CTGCCCTCCGCGACCGACCGGGCCTATCTGGAGCTGAACGGCGCGACGCACTTCTCGCCC | 780 |
| MBT1 | AACACGTCGAACACCACGATCGCGAAGTACAGCATCTCCTGGCTCAAGCGGTTCATCGAC | 828 |
| MBT2 | AACACGTCGAACACCACGATCGCGAAGTACAGCATCTCCTGGCTCAAGCGGTTCATCGAC | 828 |
| MBT5 | AACACGTCGAACACCACGATCGCGAAGTACAGCATCTCCTGGCTCAAGCGGTTCATCGAC | 828 |
| MBT12 | AACACGTCGAACACCACGATCGCGAAGTACAGCATCTCCTGGCTCAAGCGGTTCATCGAC | 828 |
| MBT28 | AACACGTCGAACACCACGATCGCGAAGTACAGCATCTCCTGGCTCAAGCGGTTCATCGAC | 828 |
| MBT29 | AACACGTCGAACACCACGATCGCGAAGTACAGCATCTCCTGGCTCAAGCGGTTCATCGAC | 828 |
| MBT38 | AACACGTCGAACACCACGATCGCGAAGTACAGCATCTCCTGGCTCAAGCGGTTCATCGAC | 828 |
| MBT39 | AACACGTCGAACACCACGATCGCGAAGTACAGCATCTCCTGGCTCAAGCGGTTCATCGAC | 828 |
| MBT47 | AACACGTCGAACACCACGATCGCGAAGTACAGCATCTCCTGGCTCAAGCGGTTCATCGAC | 828 |
| MBT50 | AACACGTCGAACACCACGATCGCGAAGTACAGCATCTCCTGGCTCAAGCGGTTCATCGAC | 828 |
| MBT3 | AACACGTCGAACACCACGATCGCGAAGTACAGCATCTCCTGGCTCAAGCGGTTCATCGAC | 828 |
| M145 | AACACGTCCAACACGACGATCGCGAAGTACAGCATCTCCTGGCTCAAGCGCTTCATCGAC | 840 |
| MBT92 | AACTCGTCCAACACGACGATCGCGAAGTACAGCGTCTCCTGGCTCAAGCGGTTCATCGAC | 840 |

```
MBT1     AACGACACCCGCTACGAGCAGTTCCTGTGCCCGCTGCCCCGGCCGAGCCTGACCATCGAG    888
MBT2     AACGACACCCGCTACGAGCAGTTCCTGTGCCCGCTGCCCCGGCCGAGCCTGACCATCGAG    888
MBT5     AACGACACCCGCTACGAGCAGTTCCTGTGCCCGCTGCCCCGGCCGAGCCTGACCATCGAG    888
MBT12    AACGACACCCGCTACGAGCAGTTCCTGTGCCCGCTGCCCCGGCCGAGCCTGACCATCGAG    888
MBT28    AACGACACCCGCTACGAGCAGTTCCTGTGCCCGCTGCCCCGGCCGAGCCTGACCATCGAG    888
MBT29    AACGACACCCGCTACGAGCAGTTCCTGTGCCCGCTGCCCCGGCCGAGCCTGACCATCGAG    888
MBT38    AACGACACCCGCTACGAGCAGTTCCTGTGCCCGCTGCCCCGGCCGAGCCTGACCATCGAG    888
MBT39    AACGACACCCGCTACGAGCAGTTCCTGTGCCCGCTGCCCCGGCCGAGCCTGACCATCGAG    888
MBT47    AACGACACCCGCTACGAGCAGTTCCTGTGCCCGCTGCCCCGGCCGAGCCTGACCATCGAG    888
MBT50    AACGACACCCGCTACGAGCAGTTCCTGTGCCCGCTGCCCCGGCCGAGCCTGACCATCGAG    888
MBT3     AACGACACCCGCTACGAGCAGTTCCTGTGCCCGCTGCCCCGGCCGAGCCTGACCATCGAG    888
M145     GACGACACCCGCTACGAGCAGTTCCTGTGCCCGCTGCCGCGGCCGAGCCTGACGATCGAG    900
MBT92    AACGACACCCGCTACGAGCAGTTCCTGTGCCCGCTGCCGCGGCCGAGTCTGACGATCGAG    900

MBT1     GAGTACCGGGGCAACTGCCCGCACGGCTCCTGA    921
MBT2     GAGTACCGGGGCAACTGCCCGCACGGCTCCTGA    921
MBT5     GAGTACCGGGGCAACTGCCCGCACGGCTCCTGA    921
MBT12    GAGTACCGGGGCAACTGCCCGCACGGCTCCTGA    921
MBT28    GAGTACCGGGGCAACTGCCCGCACGGCTCCTGA    921
MBT29    GAGTACCGGGGCAACTGCCCGCACGGCTCCTGA    921
MBT38    GAGTACCGGGGCAACTGCCCGCACGGCTCCTGA    921
MBT39    GAGTACCGGGGCAACTGCCCGCACGGCTCCTGA    921
MBT47    GAGTACCGGGGCAACTGCCCGCACGGCTCCTGA    921
MBT50    GAGTACCGGGGCAACTGCCCGCACGGCTCCTGA    921
MBT3     GAGTACCGGGGCAACTGCCCGCACGGCTCCTGA    921
M145     GAGTACCGGGGCAACTGCCCGCACGGTTCCTGA    933
MBT92    GAGTACCGGGGCAACTGCCCGCACGGCTCCTGA    933
```

#### S6 Protein alignment

```
MBT1,2,3,5,12,28,29,38,39,47,50    MQQNPHTHA----ARPAFRGPRRRLAALTAAVAAAVALTTLTGPGAQAADNPYERGPAPT   56
M145                               MQQNPHTHAAPGAARPVLRGVRRRLAAVTAAVAAVLVLGTLTGPGAQAADNPYERGPAPT   60
MBT92                              MQQNPHTHAAPGAARPVFRGVRRRLAGVTAAVAAALVLGTLTGPGAQAADNPYERGPAPT   60
                                            ****    *  *     **      *** *

MBT1,2,3,5,12,28,29,38,39,47,50    ESSIEALRGPYSVADTSVSSLAVTGFGGGTIYYPTSTSDGTFGAVVISPGFTAYQSSIAW   116
M145                               ESSIEALRGPYSVADTSVSSLAVTGFGGGTIYYPTSTSDGTFGAVVIAPGFTAYQSSIAW   120
MBT92                              ESSIEALRGPYSVADTRVSSLAVTGFGGGTIYYPTSTSDGTFGAVAIAPGFTAYQSSMAW   120
                                                   *                            *           *

MBT1,2,3,5,12,28,29,38,39,47,50    LGPRLASQGFVVFTIDTNTTLDQPDSRGRQLLAALDYLTERSSVRGRIDSSRLGVMGHSM   176
M145                               LGPRLASQGFVVFTIDTNTTLDQPDSRGRQLLAALDYLTGRSSVRGRIDSGRLGVMGHSM   180
MBT92                              LGPRLASQGFVVFTIDTNTTLDQPDSRGRQLLAALDYLTERSSVRGRIDSSRLGVMGHSM   180
                                                                          *          *

MBT1,2,3,5,12,28,29,38,39,47,50    GGGGSLEAAKSRPSLQAAIPLTPWNLDKSWPEVSTPTLIVGADGDTVAPVSSHSEPFYSS   236
M145                               GGGGTLEAAKSRPSLQAAIPLTPWNLDKSWPEVSTPTLVVGADGDTIAPVASHAEPFYSG   240
MBT92                              GGGGTLEAAKSRPSLQAAIPLTPWNLDKSWPEVSTPTLIVGADGDTIAPVASHAEPFYSG   240
                                       *                                 *       *   *  *     *

MBT1,2,3,5,12,28,29,38,39,47,50    LPSGTDRAYLELNNATHFSPNTSNTTIAKYSISWLKRFIDNDTRYEQFLCPLPRPSLTIE   296
M145                               LPSSTDRAYLELNNATHFSPNTSNTTIAKYSISWLKRFIDDDTRYEQFLCPLPRPSLTIE   300
MBT92                              LPSATDRAYLELNGATHFSPNSSNTTIAKYSVSWLKRFIDNDTRYEQFLCPLPRPSLTIE   300
                                      *         *       *         *        *

MBT1,2,3,5,12,28,29,38,39,47,50    EYRGNCPHGS*     306
M145                               EYRGNCPHGS*     310
MBT92                              EYRGNCPHGS*     310
```

#### S7.1 Knock-out (Martijn)

A LipA knock-out was created via CRISPR-Cas9 using homology-directed repair. Two homologous arms of approximately 1000bp with a 40nt overlap at both sides of LipA have been amplified via PCR on genomic DNA of M145. These homologous arms were cloned into the pCRISPomyces-2 via three-piece Gibson Assembly using a XbaI site. Additionally a sgRNA was annealed into the plasmid using Golden Gate cloning. Primers for these homologous arms bordering LipA and sgRNA were made using the protocol of Cobb and colleagues. Assembled plasmids were confirmed via sequencing.

*Table 2:* ***sgRNA and primers for homologous arms, diagnostic PCR and sequencing***

| Name | Sequence | Length (nt) | Purpose |
|---|---|---|---|
| sgRNA1-F | ACGCTCCAGCATCGAAGCCCTGCG | 24 | sgRNA |
| sgRNA1-R | AAACCGCAGGGCTTCGATGCTGGA | 24 | sgRNA |
| JA_MC_F_HA1 | TGCCGCCGGGCGTTTTTTATGGTCACCGGCCAGGACGA | 38 | Amplification Homologous Arm |
| JA_MC_F_HA2 | CGTGCGGGCAGGTGTGGGGGTTCTGCTG | 28 | Amplification Homologous Arm |
| JA_MC_R_HA1 | CCCCCACACCTGCCCGCACGGTTCCTGA | 28 | Amplification Homologous Arm |
| JA_MC_R_HA2* | CTTTTTACGGTTCCTGGCCTCGTGAGGCTGAGCGTGAG | 38 | Amplification Homologous Arm |
| JA_MRJ_F_KO_SCO0713 | GTGCACCGTTCGACGGACGA | 20 | Diagnostic PCR and forward sequencing |
| JA_R1_MBT2/12_SCO0713 | GTGCAGCAGAACCCCCACAC | 20 | Diagnostic PCR and reverse sequencing |

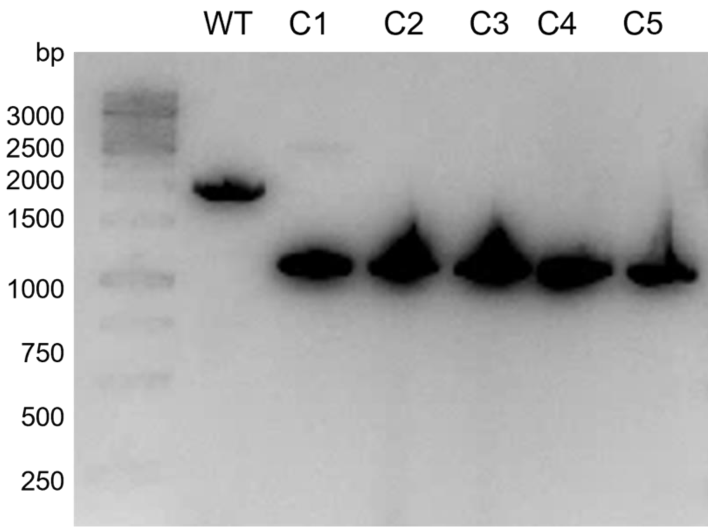

*Figure 3:* ***Diagnostic PCR of the WT and 5 knock-out colonies***
*Expected size WT is 2000bp (HA1+LipA), Expected size knock-out 1000bp (HA1).*

#### S7.2 . pSET152 cloning (Martijn)

To create overexpression strains for SCO0713 and its variants, the genes were transferred to a pSET152 integrative plasmid. This plasmid contains a phage $\varphi C31$ integrase gene which has been proven to mediate integration of plasmids into the genomic DNA of *Streptomyces* species with high efficiency. LipA of M145, MBT2 and MBT 92 was PCR amplified with overhangs containing a NdeI site on the 5' and BamHI site on the 3' end of the product. Additionally, a 6xHIS tag was added to the PCR product. pSET152 was linearized with BamHI and NdeI and PCR amplified LipA genes were cloned into the plasmid using T4 ligase. Assembled plasmids were confirmed via sequencing.

*Table 2:* ***Primers for construction pSET152_XLipA***

| Name | Sequence |
|---|---|
| JA_MC_SCO0713_pSET_F | CTGTAGCTTACATATGCAGCAGAACCCCCACAC |
| JA_MC_SCO0713_145_92_pSET_R | ATCGAGGATCCTTCAGTGGTGGTGGTGGTGGTGGGAACCGTGCGGGCAGTT |
| JA_MC_SCO0713_2_pSET_R | ATCGAGGATCCTTCAGTGGTGGTGGTGGTGGTGGGAGCCGTGCGGGCAGTT |

NdeI site is marked in RED
BamHI site is marked in Blue
His tag is marked in GREEN

### S8. Statistical analysis of BHET degradation on LC-MS

| | | | | | |
|---|---|---|---|---|---|
| Table Analyzed | All_TSBS_24h | | | | |
| Two-way ANOVA | Ordinary | | | | |
| Alpha | 0,05 | | | | |
| Source of Variation | % of total variation | P value | P value summary | Significant? | |
| Interaction | 0,8437 | <0,0001 | **** | Yes | |
| Strain | 0,04566 | 0,0205 | * | Yes | |
| Compound | 99 | <0,0001 | **** | Yes | |
| ANOVA table | SS | DF | MS | F (DFn, DFd) | P value |
| Interaction | 692,3 | 10 | 69,23 | F (10, 36) = 28,45 | P<0,0001 |
| Strain | 37,47 | 5 | 7,494 | F (5, 36) = 3,080 | P=0,0205 |
| Compound | 81244 | 2 | 40622 | F (2, 36) = 16696 | P<0,0001 |
| Residual | 87,59 | 36 | 2,433 | | |
| Data summary | | | | | |
| Number of columns (Com | 3 | | | | |
| Number of rows (Strain) | 6 | | | | |
| Number of values | 54 | | | | |
| Within each column, compare rows (simple effects within columns) | | | | | |
| Number of families | 3 | | | | |
| Number of comparisons p | 15 | | | | |
| Alpha | 0,05 | | | | |
| Tukey's multiple comparis | Mean Diff, | 95,00% CI of diff, | Below threshold? | Summary | Adjusted P Value |
| TPA | | | | | |
| Control vs. M145 | 0 | -3,832 to 3,832 | No | ns | >0,9999 |
| Control vs. ΔSCO0713 | 0 | -3,832 to 3,832 | No | ns | >0,9999 |

| | | | | | |
|---|---|---|---|---|---|
| Control vs. S3 | 0 | -3,832 to 3,832 | No | ns | >0,9999 |
| Control vs. S5 | 0 | -3,832 to 3,832 | No | ns | >0,9999 |
| Control vs. S7 | 0 | -3,832 to 3,832 | No | ns | >0,9999 |
| M145 vs. ΔSCO0713 | 0 | -3,832 to 3,832 | No | ns | >0,9999 |
| M145 vs. S3 | 0 | -3,832 to 3,832 | No | ns | >0,9999 |
| M145 vs. S5 | 0 | -3,832 to 3,832 | No | ns | >0,9999 |
| M145 vs. S7 | 0 | -3,832 to 3,832 | No | ns | >0,9999 |
| ΔSCO0713 vs. S3 | 0 | -3,832 to 3,832 | No | ns | >0,9999 |
| ΔSCO0713 vs. S5 | 0 | -3,832 to 3,832 | No | ns | >0,9999 |
| ΔSCO0713 vs. S7 | 0 | -3,832 to 3,832 | No | ns | >0,9999 |
| S3 vs. S5 | 0 | -3,832 to 3,832 | No | ns | >0,9999 |
| S3 vs. S7 | 0 | -3,832 to 3,832 | No | ns | >0,9999 |
| S5 vs. S7 | 0 | -3,832 to 3,832 | No | ns | >0,9999 |
| MHET | | | | | |
| Control vs. M145 | 3,863 | 0,03093 to 7,694 | Yes | * | 0,0472 |
| Control vs. ΔSCO0713 | 1,066 | -2,765 to 4,898 | No | ns | 0,9584 |
| Control vs. S3 | -0,001333 | -3,833 to 3,830 | No | ns | >0,9999 |
| Control vs. S5 | -4,631 | -8,463 to -0,7996 | Yes | * | 0,0103 |
| Control vs. S7 | -5,347 | -9,178 to -1,515 | Yes | ** | 0,0022 |
| M145 vs. ΔSCO0713 | -2,796 | -6,628 to 1,035 | No | ns | 0,2647 |
| M145 vs. S3 | -3,864 | -7,696 to -0,03226 | Yes | * | 0,0471 |
| M145 vs. S5 | -8,494 | -12,33 to -4,662 | Yes | **** | <0,0001 |
| M145 vs. S7 | -9,209 | -13,04 to -5,378 | Yes | **** | <0,0001 |
| ΔSCO0713 vs. S3 | -1,068 | -4,899 to 2,764 | No | ns | 0,9582 |
| ΔSCO0713 vs. S5 | -5,698 | -9,529 to -1,866 | Yes | *** | 0,001 |
| ΔSCO0713 vs. S7 | -6,413 | -10,24 to -2,581 | Yes | *** | 0,0002 |
| S3 vs. S5 | -4,63 | -8,462 to -0,7983 | Yes | * | 0,0103 |
| S3 vs. S7 | -5,345 | -9,177 to -1,514 | Yes | ** | 0,0022 |
| S5 vs. S7 | -0,7153 | -4,547 to 3,116 | No | ns | 0,9929 |
| BHET | | | | | |
| Control vs. M145 | -5,613 | -9,445 to -1,782 | Yes | ** | 0,0012 |

| | | | | | |
|---|---|---|---|---|---|
| Control vs. ΔSCO0713 | 1,847 | -1,985 to 5,678 | No | ns | 0,697 |
| Control vs. S3 | -0,46 | -4,292 to 3,372 | No | ns | 0,9991 |
| Control vs. S5 | 8,717 | 4,885 to 12,55 | Yes | **** | <0,0001 |
| Control vs. S7 | 10,34 | 6,512 to 14,18 | Yes | **** | <0,0001 |
| M145 vs. ΔSCO0713 | 7,46 | 3,628 to 11,29 | Yes | **** | <0,0001 |
| M145 vs. S3 | 5,153 | 1,322 to 8,985 | Yes | ** | 0,0033 |
| M145 vs. S5 | 14,33 | 10,50 to 18,16 | Yes | **** | <0,0001 |
| M145 vs. S7 | 15,96 | 12,12 to 19,79 | Yes | **** | <0,0001 |
| ΔSCO0713 vs. S3 | -2,307 | -6,138 to 1,525 | No | ns | 0,4717 |
| ΔSCO0713 vs. S5 | 6,87 | 3,038 to 10,70 | Yes | **** | <0,0001 |
| ΔSCO0713 vs. S7 | 8,497 | 4,665 to 12,33 | Yes | **** | <0,0001 |
| S3 vs. S5 | 9,177 | 5,345 to 13,01 | Yes | **** | <0,0001 |
| S3 vs. S7 | 10,8 | 6,972 to 14,64 | Yes | **** | <0,0001 |
| S5 vs. S7 | 1,627 | -2,205 to 5,458 | No | ns | 0,7952 |

| Test details | Mean 1 | Mean 2 | Mean Diff, | SE of diff, | N1 | N2 | q | DF |
|---|---|---|---|---|---|---|---|---|
| TPA | | | | | | | | |
| Control vs. M145 | 0 | 0 | 0 | 1,274 | 3 | 3 | 0 | 36 |
| Control vs. ΔSCO0713 | 0 | 0 | 0 | 1,274 | 3 | 3 | 0 | 36 |
| Control vs. S3 | 0 | 0 | 0 | 1,274 | 3 | 3 | 0 | 36 |
| Control vs. S5 | 0 | 0 | 0 | 1,274 | 3 | 3 | 0 | 36 |
| Control vs. S7 | 0 | 0 | 0 | 1,274 | 3 | 3 | 0 | 36 |
| M145 vs. ΔSCO0713 | 0 | 0 | 0 | 1,274 | 3 | 3 | 0 | 36 |
| M145 vs. S3 | 0 | 0 | 0 | 1,274 | 3 | 3 | 0 | 36 |
| M145 vs. S5 | 0 | 0 | 0 | 1,274 | 3 | 3 | 0 | 36 |
| M145 vs. S7 | 0 | 0 | 0 | 1,274 | 3 | 3 | 0 | 36 |
| ΔSCO0713 vs. S3 | 0 | 0 | 0 | 1,274 | 3 | 3 | 0 | 36 |
| ΔSCO0713 vs. S5 | 0 | 0 | 0 | 1,274 | 3 | 3 | 0 | 36 |
| ΔSCO0713 vs. S7 | 0 | 0 | 0 | 1,274 | 3 | 3 | 0 | 36 |
| S3 vs. S5 | 0 | 0 | 0 | 1,274 | 3 | 3 | 0 | 36 |
| S3 vs. S7 | 0 | 0 | 0 | 1,274 | 3 | 3 | 0 | 36 |

| | | | | | | | | |
|---|---|---|---|---|---|---|---|---|
| S5 vs. S7 | 0 | 0 | 0 | 1,274 | 3 | 3 | 0 | 36 |
| MHET | | | | | | | | |
| Control vs. M145 | 5,867 | 2,004 | 3,863 | 1,274 | 3 | 3 | 4,289 | 36 |
| Control vs. ΔSCO0713 | 5,867 | 4,8 | 1,066 | 1,274 | 3 | 3 | 1,184 | 36 |
| Control vs. S3 | 5,867 | 5,868 | -0,001333 | 1,274 | 3 | 3 | 0,001481 | 36 |
| Control vs. S5 | 5,867 | 10,5 | -4,631 | 1,274 | 3 | 3 | 5,143 | 36 |
| Control vs. S7 | 5,867 | 11,21 | -5,347 | 1,274 | 3 | 3 | 5,937 | 36 |
| M145 vs. ΔSCO0713 | 2,004 | 4,8 | -2,796 | 1,274 | 3 | 3 | 3,105 | 36 |
| M145 vs. S3 | 2,004 | 5,868 | -3,864 | 1,274 | 3 | 3 | 4,291 | 36 |
| M145 vs. S5 | 2,004 | 10,5 | -8,494 | 1,274 | 3 | 3 | 9,432 | 36 |
| M145 vs. S7 | 2,004 | 11,21 | -9,209 | 1,274 | 3 | 3 | 10,23 | 36 |
| ΔSCO0713 vs. S3 | 4,8 | 5,868 | -1,068 | 1,274 | 3 | 3 | 1,186 | 36 |
| ΔSCO0713 vs. S5 | 4,8 | 10,5 | -5,698 | 1,274 | 3 | 3 | 6,327 | 36 |
| ΔSCO0713 vs. S7 | 4,8 | 11,21 | -6,413 | 1,274 | 3 | 3 | 7,121 | 36 |
| S3 vs. S5 | 5,868 | 10,5 | -4,63 | 1,274 | 3 | 3 | 5,141 | 36 |
| S3 vs. S7 | 5,868 | 11,21 | -5,345 | 1,274 | 3 | 3 | 5,935 | 36 |
| S5 vs. S7 | 10,5 | 11,21 | -0,7153 | 1,274 | 3 | 3 | 0,7943 | 36 |
| BHET | | | | | | | | |
| Control vs. M145 | 87,9 | 93,52 | -5,613 | 1,274 | 3 | 3 | 6,233 | 36 |
| Control vs. ΔSCO0713 | 87,9 | 86,06 | 1,847 | 1,274 | 3 | 3 | 2,051 | 36 |
| Control vs. S3 | 87,9 | 88,36 | -0,46 | 1,274 | 3 | 3 | 0,5108 | 36 |
| Control vs. S5 | 87,9 | 79,19 | 8,717 | 1,274 | 3 | 3 | 9,679 | 36 |
| Control vs. S7 | 87,9 | 77,56 | 10,34 | 1,274 | 3 | 3 | 11,49 | 36 |
| M145 vs. ΔSCO0713 | 93,52 | 86,06 | 7,46 | 1,274 | 3 | 3 | 8,284 | 36 |
| M145 vs. S3 | 93,52 | 88,36 | 5,153 | 1,274 | 3 | 3 | 5,722 | 36 |
| M145 vs. S5 | 93,52 | 79,19 | 14,33 | 1,274 | 3 | 3 | 15,91 | 36 |
| M145 vs. S7 | 93,52 | 77,56 | 15,96 | 1,274 | 3 | 3 | 17,72 | 36 |
| ΔSCO0713 vs. S3 | 86,06 | 88,36 | -2,307 | 1,274 | 3 | 3 | 2,561 | 36 |
| ΔSCO0713 vs. S5 | 86,06 | 79,19 | 6,87 | 1,274 | 3 | 3 | 7,628 | 36 |
| ΔSCO0713 vs. S7 | 86,06 | 77,56 | 8,497 | 1,274 | 3 | 3 | 9,435 | 36 |
| S3 vs. S5 | 88,36 | 79,19 | 9,177 | 1,274 | 3 | 3 | 10,19 | 36 |

| | | | | | | | | |
|---|---|---|---|---|---|---|---|---|
| S3 vs. S7 | 88,36 | 77,56 | 10,8 | 1,274 | 3 | 3 | 12 | 36 |
| S5 vs. S7 | 79,19 | 77,56 | 1,627 | 1,274 | 3 | 3 | 1,806 | 36 |

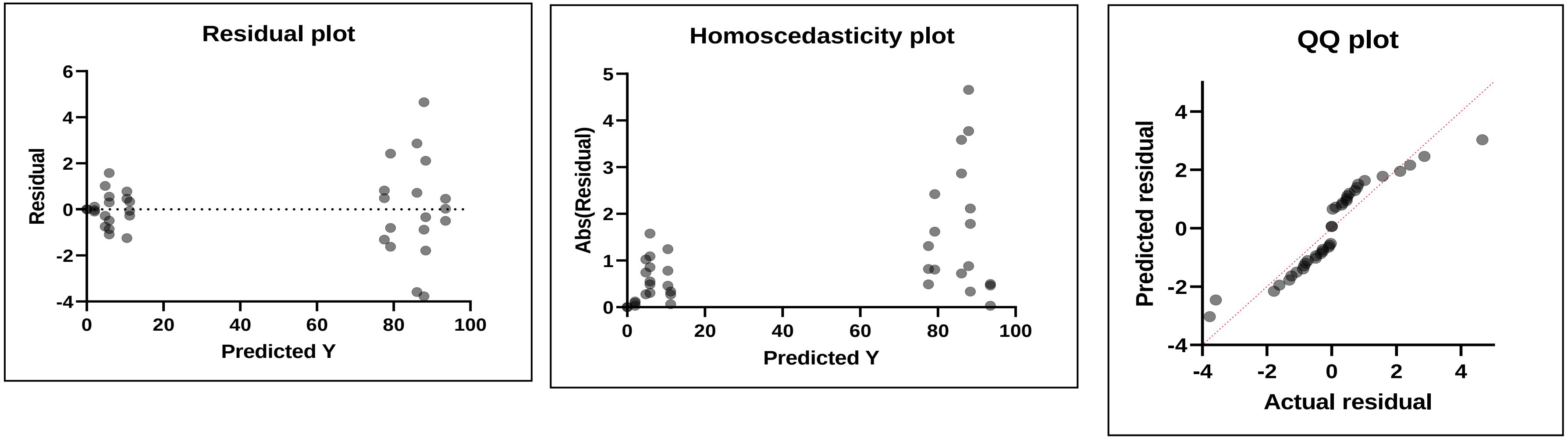

| | |
|---|---|
| Table Analyzed | **All_TSBS_48h** |

| | |
|---|---|
| Two-way ANOVA | Ordinary |
| Alpha | 0,05 |

| Source of Variation | % of total variation | P value | P value sur | Significant? |
|---|---|---|---|---|
| Interaction | 56,8 | <0,0001 | **** | Yes |
| strain | 0,01644 | 0,9437 | ns | No |
| compound | 33,96 | <0,0001 | **** | Yes |

| ANOVA table | SS (Type III) | DF | MS | F (DFn, DF | P value |
|---|---|---|---|---|---|
| Interaction | 40955 | 10 | 4096 | F (10, 34) = | P<0,0001 |
| strain | 11,86 | 5 | 2,371 | F (5, 34) = | P=0,9437 |
| compound | 24484 | 2 | 12242 | F (2, 34) = | P<0,0001 |
| Residual | 341 | 34 | 10,03 | | |

| Data summary | |
|---|---|
| Number of columns (compound) | 3 |
| Number of rows (strain) | 6 |
| Number of values | 52 |

Within each column, compare rows (simple effects within columns)

| | |
|---|---|
| Number of families | 3 |
| Number of comparisons per family | 15 |
| Alpha | 0,05 |

| Tukey's multiple comparisons test | Predicted (LS) mear | 95,00% CI | Below thres | Summary | Adjusted P Value |
|---|---|---|---|---|---|
| TPA | | | | | |
| Control vs. M145 | 1,776E-14 | -7,805 to 7, | No | ns | >0,9999 |
| Control vs. ΔSCO0713 | 1,421E-14 | -7,805 to 7, | No | ns | >0,9999 |

| | | | | | |
|---|---|---|---|---|---|
| Control vs. S3 | -1,421E-14 | -7,805 to 7, | No | ns | >0,9999 |
| Control vs. S5 | -1,421E-14 | -7,805 to 7, | No | ns | >0,9999 |
| Control vs. S7 | -3,553E-14 | -7,805 to 7, | No | ns | >0,9999 |
| M145 vs. ΔSCO0713 | -3,553E-15 | -7,805 to 7, | No | ns | >0,9999 |
| M145 vs. S3 | -3,197E-14 | -7,805 to 7, | No | ns | >0,9999 |
| M145 vs. S5 | -3,197E-14 | -7,805 to 7, | No | ns | >0,9999 |
| M145 vs. S7 | -5,329E-14 | -7,805 to 7, | No | ns | >0,9999 |
| ΔSCO0713 vs. S3 | -2,842E-14 | -7,805 to 7, | No | ns | >0,9999 |
| ΔSCO0713 vs. S5 | -2,842E-14 | -7,805 to 7, | No | ns | >0,9999 |
| ΔSCO0713 vs. S7 | -4,974E-14 | -7,805 to 7, | No | ns | >0,9999 |
| S3 vs. S5 | 0 | -7,805 to 7, | No | ns | >0,9999 |
| S3 vs. S7 | -2,132E-14 | -7,805 to 7, | No | ns | >0,9999 |
| S5 vs. S7 | -2,132E-14 | -7,805 to 7, | No | ns | >0,9999 |
| MHET | | | | | |
| Control vs. M145 | -2,238 | -10,04 to 5, | No | ns | 0,9521 |
| Control vs. ΔSCO0713 | -4,993 | -12,80 to 2, | No | ns | 0,4017 |
| Control vs. S3 | -41,14 | -48,95 to -3 | Yes | **** | <0,0001 |
| Control vs. S5 | -83,83 | -91,63 to -7 | Yes | **** | <0,0001 |
| Control vs. S7 | -84,8 | -92,61 to -7 | Yes | **** | <0,0001 |
| M145 vs. ΔSCO0713 | -2,755 | -10,56 to 5, | No | ns | 0,8915 |
| M145 vs. S3 | -38,91 | -46,71 to -3 | Yes | **** | <0,0001 |
| M145 vs. S5 | -81,59 | -89,39 to -7 | Yes | **** | <0,0001 |
| M145 vs. S7 | -82,57 | -90,37 to -7 | Yes | **** | <0,0001 |
| ΔSCO0713 vs. S3 | -36,15 | -43,96 to -2 | Yes | **** | <0,0001 |
| ΔSCO0713 vs. S5 | -78,83 | -86,64 to -7 | Yes | **** | <0,0001 |
| ΔSCO0713 vs. S7 | -79,81 | -87,62 to -7 | Yes | **** | <0,0001 |
| S3 vs. S5 | -42,68 | -50,49 to -3 | Yes | **** | <0,0001 |
| S3 vs. S7 | -43,66 | -51,47 to -3 | Yes | **** | <0,0001 |
| S5 vs. S7 | -0,9767 | -8,782 to 6, | No | ns | 0,9989 |
| BHET | | | | | |
| Control vs. M145 | -0,73 | -8,535 to 7, | No | ns | 0,9997 |

| | | | | | |
|---|---|---|---|---|---|
| Control vs. ΔSCO0713 | 4,717 | -3,088 to 1: | No | ns | 0,4646 |
| Control vs. S3 | 37,38 | 29,58 to 45 | Yes | **** | <0,0001 |
| Control vs. S5 | 81,2 | 73,39 to 89 | Yes | **** | <0,0001 |
| Control vs. S7 | 81,93 | 70,89 to 92 | Yes | **** | <0,0001 |
| M145 vs. ΔSCO0713 | 5,447 | -2,358 to 1: | No | ns | 0,3082 |
| M145 vs. S3 | 38,11 | 30,31 to 45 | Yes | **** | <0,0001 |
| M145 vs. S5 | 81,93 | 74,12 to 89 | Yes | **** | <0,0001 |
| M145 vs. S7 | 82,66 | 71,62 to 93 | Yes | **** | <0,0001 |
| ΔSCO0713 vs. S3 | 32,67 | 24,86 to 40 | Yes | **** | <0,0001 |
| ΔSCO0713 vs. S5 | 76,48 | 68,68 to 84 | Yes | **** | <0,0001 |
| ΔSCO0713 vs. S7 | 77,21 | 66,17 to 88 | Yes | **** | <0,0001 |
| S3 vs. S5 | 43,81 | 36,01 to 51 | Yes | **** | <0,0001 |
| S3 vs. S7 | 44,54 | 33,51 to 55 | Yes | **** | <0,0001 |
| S5 vs. S7 | 0,7297 | -10,31 to 1: | No | ns | >0,9999 |

| Test details | Predicted (LS) mear | Predicted (I | Predicted (I | SE of diff, | N1 | N2 | q | DF |
|---|---|---|---|---|---|---|---|---|
| TPA | | | | | | | | |
| Control vs. M145 | -1,421E-14 | -3,2E-14 | 1,78E-14 | 2,586 | 3 | 3 | 9,72E-15 | 34 |
| Control vs. ΔSCO0713 | -1,421E-14 | -2,84E-14 | 1,42E-14 | 2,586 | 3 | 3 | 7,77E-15 | 34 |
| Control vs. S3 | -1,421E-14 | 0 | -1,42E-14 | 2,586 | 3 | 3 | 7,77E-15 | 34 |
| Control vs. S5 | -1,421E-14 | 0 | -1,42E-14 | 2,586 | 3 | 3 | 7,77E-15 | 34 |
| Control vs. S7 | -1,421E-14 | 2,13E-14 | -3,55E-14 | 2,586 | 3 | 3 | 1,94E-14 | 34 |
| M145 vs. ΔSCO0713 | -3,197E-14 | -2,84E-14 | -3,55E-15 | 2,586 | 3 | 3 | 1,94E-15 | 34 |
| M145 vs. S3 | -3,197E-14 | 0 | -3,2E-14 | 2,586 | 3 | 3 | 1,75E-14 | 34 |
| M145 vs. S5 | -3,197E-14 | 0 | -3,2E-14 | 2,586 | 3 | 3 | 1,75E-14 | 34 |
| M145 vs. S7 | -3,197E-14 | 2,13E-14 | -5,33E-14 | 2,586 | 3 | 3 | 2,91E-14 | 34 |
| ΔSCO0713 vs. S3 | -2,842E-14 | 0 | -2,84E-14 | 2,586 | 3 | 3 | 1,55E-14 | 34 |
| ΔSCO0713 vs. S5 | -2,842E-14 | 0 | -2,84E-14 | 2,586 | 3 | 3 | 1,55E-14 | 34 |
| ΔSCO0713 vs. S7 | -2,842E-14 | 2,13E-14 | -4,97E-14 | 2,586 | 3 | 3 | 2,72E-14 | 34 |
| S3 vs. S5 | 0 | 0 | 0 | 2,586 | 3 | 3 | 0 | 34 |
| S3 vs. S7 | 0 | 2,13E-14 | -2,13E-14 | 2,586 | 3 | 3 | 1,17E-14 | 34 |

| | | | | | | | | |
|---|---|---|---|---|---|---|---|---|
| S5 vs. S7 | 0 | 2,13E-14 | -2,13E-14 | 2,586 | 3 | 3 | 1,17E-14 | 34 |
| MHET | | | | | | | | |
| Control vs. M145 | 7,727 | 9,965 | -2,238 | 2,586 | 3 | 3 | 1,224 | 34 |
| Control vs. ΔSCO0713 | 7,727 | 12,72 | -4,993 | 2,586 | 3 | 3 | 2,731 | 34 |
| Control vs. S3 | 7,727 | 48,87 | -41,14 | 2,586 | 3 | 3 | 22,5 | 34 |
| Control vs. S5 | 7,727 | 91,55 | -83,83 | 2,586 | 3 | 3 | 45,84 | 34 |
| Control vs. S7 | 7,727 | 92,53 | -84,8 | 2,586 | 3 | 3 | 46,38 | 34 |
| M145 vs. ΔSCO0713 | 9,965 | 12,72 | -2,755 | 2,586 | 3 | 3 | 1,507 | 34 |
| M145 vs. S3 | 9,965 | 48,87 | -38,91 | 2,586 | 3 | 3 | 21,28 | 34 |
| M145 vs. S5 | 9,965 | 91,55 | -81,59 | 2,586 | 3 | 3 | 44,62 | 34 |
| M145 vs. S7 | 9,965 | 92,53 | -82,57 | 2,586 | 3 | 3 | 45,15 | 34 |
| ΔSCO0713 vs. S3 | 12,72 | 48,87 | -36,15 | 2,586 | 3 | 3 | 19,77 | 34 |
| ΔSCO0713 vs. S5 | 12,72 | 91,55 | -78,83 | 2,586 | 3 | 3 | 43,11 | 34 |
| ΔSCO0713 vs. S7 | 12,72 | 92,53 | -79,81 | 2,586 | 3 | 3 | 43,65 | 34 |
| S3 vs. S5 | 48,87 | 91,55 | -42,68 | 2,586 | 3 | 3 | 23,34 | 34 |
| S3 vs. S7 | 48,87 | 92,53 | -43,66 | 2,586 | 3 | 3 | 23,88 | 34 |
| S5 vs. S7 | 91,55 | 92,53 | -0,9767 | 2,586 | 3 | 3 | 0,5341 | 34 |
| BHET | | | | | | | | |
| Control vs. M145 | 83 | 83,73 | -0,73 | 2,586 | 3 | 3 | 0,3992 | 34 |
| Control vs. ΔSCO0713 | 83 | 78,29 | 4,717 | 2,586 | 3 | 3 | 2,579 | 34 |
| Control vs. S3 | 83 | 45,62 | 37,38 | 2,586 | 3 | 3 | 20,44 | 34 |
| Control vs. S5 | 83 | 1,806 | 81,2 | 2,586 | 3 | 3 | 44,41 | 34 |
| Control vs. S7 | 83 | 1,076 | 81,93 | 3,657 | 3 | 1 | 31,68 | 34 |
| M145 vs. ΔSCO0713 | 83,73 | 78,29 | 5,447 | 2,586 | 3 | 3 | 2,979 | 34 |
| M145 vs. S3 | 83,73 | 45,62 | 38,11 | 2,586 | 3 | 3 | 20,84 | 34 |
| M145 vs. S5 | 83,73 | 1,806 | 81,93 | 2,586 | 3 | 3 | 44,8 | 34 |
| M145 vs. S7 | 83,73 | 1,076 | 82,66 | 3,657 | 3 | 1 | 31,96 | 34 |
| ΔSCO0713 vs. S3 | 78,29 | 45,62 | 32,67 | 2,586 | 3 | 3 | 17,86 | 34 |
| ΔSCO0713 vs. S5 | 78,29 | 1,806 | 76,48 | 2,586 | 3 | 3 | 41,83 | 34 |
| ΔSCO0713 vs. S7 | 78,29 | 1,076 | 77,21 | 3,657 | 3 | 1 | 29,86 | 34 |
| S3 vs. S5 | 45,62 | 1,806 | 43,81 | 2,586 | 3 | 3 | 23,96 | 34 |

| | | | | | | | | |
|---|---|---|---|---|---|---|---|---|
| S3 vs. S7 | 45,62 | 1,076 | 44,54 | 3,657 | 3 | 1 | 17,23 | 34 |
| S5 vs. S7 | 1,806 | 1,076 | 0,7297 | 3,657 | 3 | 1 | 0,2822 | 34 |

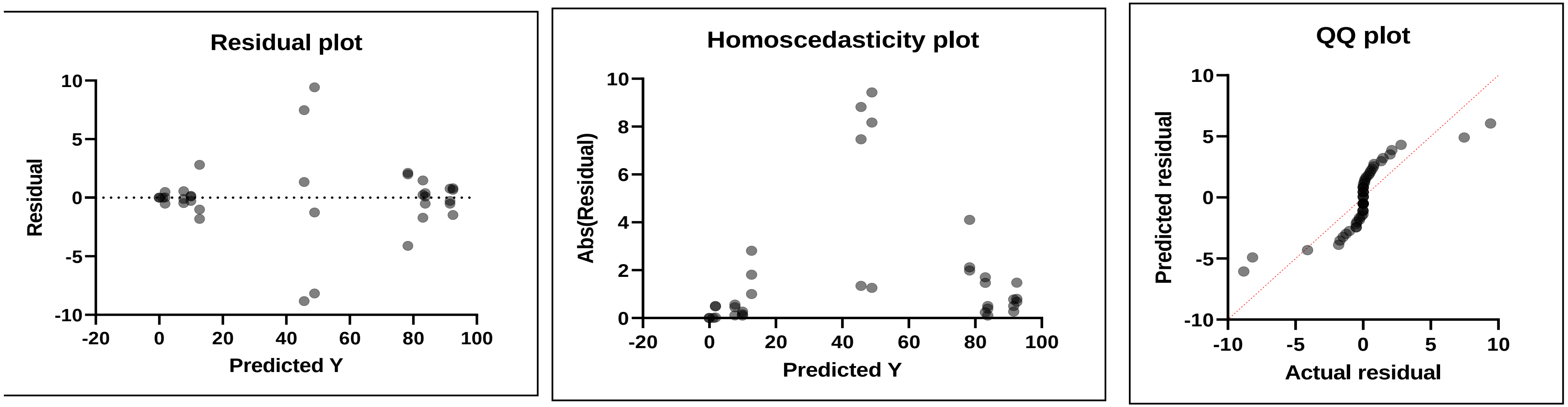

### All_TSBS_72h

Within each column, compare rows (simple effects within columns)

| | |
|---|---|
| Number of families | 3 |
| Number of comparisons per family | 15 |
| Alpha | 0,05 |

| Tukey's multiple comparisons test | Mean Diff, | 95,00% CI of diff, | Below thres | Summary | Adjusted P Value |
|---|---|---|---|---|---|
| TPA | | | | | |
| Control vs. M145 | 0 | -2,498 to 2,498 | No | ns | >0,9999 |
| Control vs. ΔSCO0713 | 0 | -2,498 to 2,498 | No | ns | >0,9999 |
| Control vs. S3 | -1,093 | -3,591 to 1,405 | No | ns | 0,774 |
| Control vs. S5 | -7,507 | -10,01 to -5,009 | Yes | **** | <0,0001 |
| Control vs. S7 | -11,37 | -13,87 to -8,872 | Yes | **** | <0,0001 |
| M145 vs. ΔSCO0713 | 0 | -2,498 to 2,498 | No | ns | >0,9999 |
| M145 vs. S3 | -1,093 | -3,591 to 1,405 | No | ns | 0,774 |
| M145 vs. S5 | -7,507 | -10,01 to -5,009 | Yes | **** | <0,0001 |
| M145 vs. S7 | -11,37 | -13,87 to -8,872 | Yes | **** | <0,0001 |
| ΔSCO0713 vs. S3 | -1,093 | -3,591 to 1,405 | No | ns | 0,774 |
| ΔSCO0713 vs. S5 | -7,507 | -10,01 to -5,009 | Yes | **** | <0,0001 |
| ΔSCO0713 vs. S7 | -11,37 | -13,87 to -8,872 | Yes | **** | <0,0001 |
| S3 vs. S5 | -6,414 | -8,912 to -3,916 | Yes | **** | <0,0001 |
| S3 vs. S7 | -10,28 | -12,77 to -7,779 | Yes | **** | <0,0001 |
| S5 vs. S7 | -3,863 | -6,361 to -1,365 | Yes | *** | 0,0006 |
| MHET | | | | | |
| Control vs. M145 | -23,53 | -26,03 to -21,04 | Yes | **** | <0,0001 |
| Control vs. ΔSCO0713 | -15,77 | -18,27 to -13,28 | Yes | **** | <0,0001 |
| Control vs. S3 | -83,55 | -86,04 to -81,05 | Yes | **** | <0,0001 |
| Control vs. S5 | -73,77 | -76,26 to -71,27 | Yes | **** | <0,0001 |
| Control vs. S7 | -68,47 | -70,97 to -65,98 | Yes | **** | <0,0001 |
| M145 vs. ΔSCO0713 | 7,76 | 5,262 to 10,26 | Yes | **** | <0,0001 |
| M145 vs. S3 | -60,01 | -62,51 to -57,52 | Yes | **** | <0,0001 |

| | | | | | |
|---|---|---|---|---|---|
| M145 vs. S5 | -50,23 | -52,73 to -47,74 | Yes | **** | <0,0001 |
| M145 vs. S7 | -44,94 | -47,44 to -42,44 | Yes | **** | <0,0001 |
| ΔSCO0713 vs. S3 | -67,77 | -70,27 to -65,28 | Yes | **** | <0,0001 |
| ΔSCO0713 vs. S5 | -57,99 | -60,49 to -55,50 | Yes | **** | <0,0001 |
| ΔSCO0713 vs. S7 | -52,7 | -55,20 to -50,20 | Yes | **** | <0,0001 |
| S3 vs. S5 | 9,78 | 7,282 to 12,28 | Yes | **** | <0,0001 |
| S3 vs. S7 | 15,07 | 12,58 to 17,57 | Yes | **** | <0,0001 |
| S5 vs. S7 | 5,293 | 2,795 to 7,791 | Yes | **** | <0,0001 |
| BHET | | | | | |
| Control vs. M145 | 21,43 | 18,93 to 23,92 | Yes | **** | <0,0001 |
| Control vs. ΔSCO0713 | 16,09 | 13,59 to 18,58 | Yes | **** | <0,0001 |
| Control vs. S3 | 81,89 | 79,39 to 84,38 | Yes | **** | <0,0001 |
| Control vs. S5 | 81,89 | 79,39 to 84,38 | Yes | **** | <0,0001 |
| Control vs. S7 | 81,89 | 79,39 to 84,38 | Yes | **** | <0,0001 |
| M145 vs. ΔSCO0713 | -5,34 | -7,838 to -2,842 | Yes | **** | <0,0001 |
| M145 vs. S3 | 60,46 | 57,96 to 62,96 | Yes | **** | <0,0001 |
| M145 vs. S5 | 60,46 | 57,96 to 62,96 | Yes | **** | <0,0001 |
| M145 vs. S7 | 60,46 | 57,96 to 62,96 | Yes | **** | <0,0001 |
| ΔSCO0713 vs. S3 | 65,8 | 63,30 to 68,30 | Yes | **** | <0,0001 |
| ΔSCO0713 vs. S5 | 65,8 | 63,30 to 68,30 | Yes | **** | <0,0001 |
| ΔSCO0713 vs. S7 | 65,8 | 63,30 to 68,30 | Yes | **** | <0,0001 |
| S3 vs. S5 | 0 | -2,498 to 2,498 | No | ns | >0,9999 |
| S3 vs. S7 | 0 | -2,498 to 2,498 | No | ns | >0,9999 |
| S5 vs. S7 | 0 | -2,498 to 2,498 | No | ns | >0,9999 |

| Test details | Mean 1 | Mean 2 | Mean Diff, | SE of diff, | N1 | N2 | q | DF |
|---|---|---|---|---|---|---|---|---|
| TPA | | | | | | | | |
| Control vs. M145 | 0 | 0 | 0 | 0,8303 | 3 | 3 | 0 | 36 |
| Control vs. ΔSCO0713 | 0 | 0 | 0 | 0,8303 | 3 | 3 | 0 | 36 |
| Control vs. S3 | 0 | 1,093 | -1,093 | 0,8303 | 3 | 3 | 1,862 | 36 |

| | | | | | | | | |
|---|---|---|---|---|---|---|---|---|
| Control vs. S5 | 0 | 7,507 | -7,507 | 0,8303 | 3 | 3 | 12,79 | 36 |
| Control vs. S7 | 0 | 11,37 | -11,37 | 0,8303 | 3 | 3 | 19,37 | 36 |
| M145 vs. ΔSCO0713 | 0 | 0 | 0 | 0,8303 | 3 | 3 | 0 | 36 |
| M145 vs. S3 | 0 | 1,093 | -1,093 | 0,8303 | 3 | 3 | 1,862 | 36 |
| M145 vs. S5 | 0 | 7,507 | -7,507 | 0,8303 | 3 | 3 | 12,79 | 36 |
| M145 vs. S7 | 0 | 11,37 | -11,37 | 0,8303 | 3 | 3 | 19,37 | 36 |
| ΔSCO0713 vs. S3 | 0 | 1,093 | -1,093 | 0,8303 | 3 | 3 | 1,862 | 36 |
| ΔSCO0713 vs. S5 | 0 | 7,507 | -7,507 | 0,8303 | 3 | 3 | 12,79 | 36 |
| ΔSCO0713 vs. S7 | 0 | 11,37 | -11,37 | 0,8303 | 3 | 3 | 19,37 | 36 |
| S3 vs. S5 | 1,093 | 7,507 | -6,414 | 0,8303 | 3 | 3 | 10,93 | 36 |
| S3 vs. S7 | 1,093 | 11,37 | -10,28 | 0,8303 | 3 | 3 | 17,5 | 36 |
| S5 vs. S7 | 7,507 | 11,37 | -3,863 | 0,8303 | 3 | 3 | 6,579 | 36 |
| MHET | | | | | | | | |
| Control vs. M145 | 8,557 | 32,09 | -23,53 | 0,8303 | 3 | 3 | 40,08 | 36 |
| Control vs. ΔSCO0713 | 8,557 | 24,33 | -15,77 | 0,8303 | 3 | 3 | 26,87 | 36 |
| Control vs. S3 | 8,557 | 92,1 | -83,55 | 0,8303 | 3 | 3 | 142,3 | 36 |
| Control vs. S5 | 8,557 | 82,32 | -73,77 | 0,8303 | 3 | 3 | 125,6 | 36 |
| Control vs. S7 | 8,557 | 77,03 | -68,47 | 0,8303 | 3 | 3 | 116,6 | 36 |
| M145 vs. ΔSCO0713 | 32,09 | 24,33 | 7,76 | 0,8303 | 3 | 3 | 13,22 | 36 |
| M145 vs. S3 | 32,09 | 92,1 | -60,01 | 0,8303 | 3 | 3 | 102,2 | 36 |
| M145 vs. S5 | 32,09 | 82,32 | -50,23 | 0,8303 | 3 | 3 | 85,56 | 36 |
| M145 vs. S7 | 32,09 | 77,03 | -44,94 | 0,8303 | 3 | 3 | 76,55 | 36 |
| ΔSCO0713 vs. S3 | 24,33 | 92,1 | -67,77 | 0,8303 | 3 | 3 | 115,4 | 36 |
| ΔSCO0713 vs. S5 | 24,33 | 82,32 | -57,99 | 0,8303 | 3 | 3 | 98,78 | 36 |
| ΔSCO0713 vs. S7 | 24,33 | 77,03 | -52,7 | 0,8303 | 3 | 3 | 89,76 | 36 |
| S3 vs. S5 | 92,1 | 82,32 | 9,78 | 0,8303 | 3 | 3 | 16,66 | 36 |
| S3 vs. S7 | 92,1 | 77,03 | 15,07 | 0,8303 | 3 | 3 | 25,67 | 36 |
| S5 vs. S7 | 82,32 | 77,03 | 5,293 | 0,8303 | 3 | 3 | 9,016 | 36 |
| BHET | | | | | | | | |
| Control vs. M145 | 81,89 | 60,46 | 21,43 | 0,8303 | 3 | 3 | 36,5 | 36 |
| Control vs. ΔSCO0713 | 81,89 | 65,8 | 16,09 | 0,8303 | 3 | 3 | 27,4 | 36 |

| | | | | | | | | |
|---|---|---|---|---|---|---|---|---|
| Control vs. S3 | 81,89 | 0 | 81,89 | 0,8303 | 3 | 3 | 139,5 | 36 |
| Control vs. S5 | 81,89 | 0 | 81,89 | 0,8303 | 3 | 3 | 139,5 | 36 |
| Control vs. S7 | 81,89 | 0 | 81,89 | 0,8303 | 3 | 3 | 139,5 | 36 |
| M145 vs. ΔSCO0713 | 60,46 | 65,8 | -5,34 | 0,8303 | 3 | 3 | 9,096 | 36 |
| M145 vs. S3 | 60,46 | 0 | 60,46 | 0,8303 | 3 | 3 | 103 | 36 |
| M145 vs. S5 | 60,46 | 0 | 60,46 | 0,8303 | 3 | 3 | 103 | 36 |
| M145 vs. S7 | 60,46 | 0 | 60,46 | 0,8303 | 3 | 3 | 103 | 36 |
| ΔSCO0713 vs. S3 | 65,8 | 0 | 65,8 | 0,8303 | 3 | 3 | 112,1 | 36 |
| ΔSCO0713 vs. S5 | 65,8 | 0 | 65,8 | 0,8303 | 3 | 3 | 112,1 | 36 |
| ΔSCO0713 vs. S7 | 65,8 | 0 | 65,8 | 0,8303 | 3 | 3 | 112,1 | 36 |
| S3 vs. S5 | 0 | 0 | 0 | 0,8303 | 3 | 3 | 0 | 36 |
| S3 vs. S7 | 0 | 0 | 0 | 0,8303 | 3 | 3 | 0 | 36 |
| S5 vs. S7 | 0 | 0 | 0 | 0,8303 | 3 | 3 | 0 | 36 |

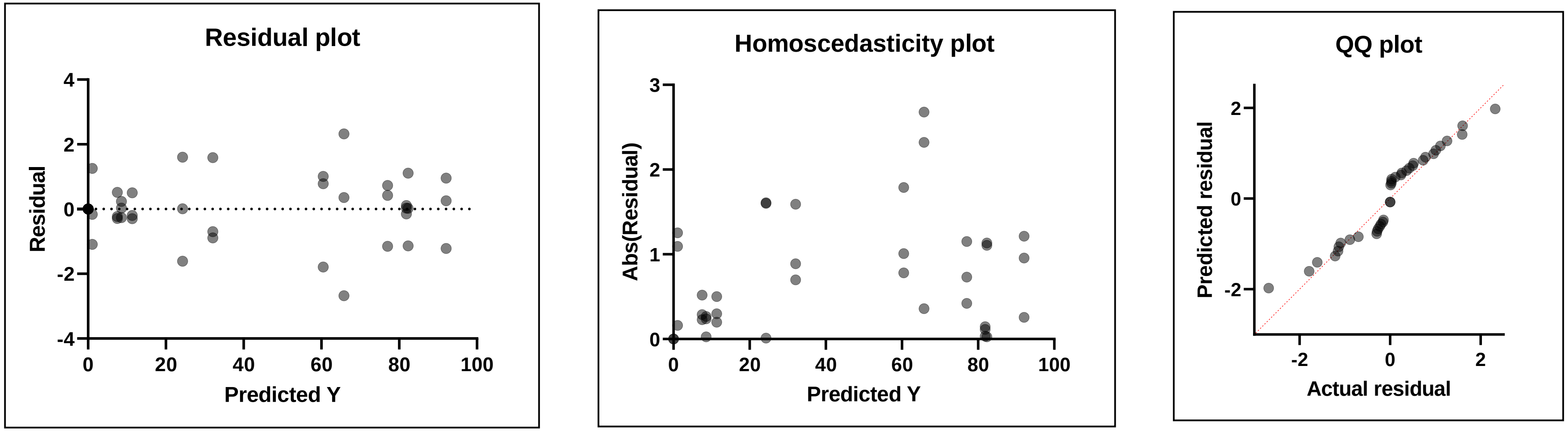

### All_NMM_24h

Within each column, compare rows (simple effects within columns)

| | |
|---|---|
| Number of families | 3 |
| Number of comparisons per f | 15 |
| Alpha | 0,05 |

| Tukey's multiple comparisons | Mean Diff, | 95,00% CI of diff, | Below threshold? | Summary | Adjusted P Value |
|---|---|---|---|---|---|
| TPA | | | | | |
| Control vs. M145 | 0 | -0,3577 to 0,3577 | No | ns | >0,9999 |
| Control vs. ΔSCO0713 | 0 | -0,3577 to 0,3577 | No | ns | >0,9999 |
| Control vs. S3 | 0 | -0,3577 to 0,3577 | No | ns | >0,9999 |
| Control vs. S5 | 0 | -0,3577 to 0,3577 | No | ns | >0,9999 |
| Control vs. S7 | 0 | -0,3577 to 0,3577 | No | ns | >0,9999 |
| M145 vs. ΔSCO0713 | 0 | -0,3577 to 0,3577 | No | ns | >0,9999 |
| M145 vs. S3 | 0 | -0,3577 to 0,3577 | No | ns | >0,9999 |
| M145 vs. S5 | 0 | -0,3577 to 0,3577 | No | ns | >0,9999 |
| M145 vs. S7 | 0 | -0,3577 to 0,3577 | No | ns | >0,9999 |
| ΔSCO0713 vs. S3 | 0 | -0,3577 to 0,3577 | No | ns | >0,9999 |
| ΔSCO0713 vs. S5 | 0 | -0,3577 to 0,3577 | No | ns | >0,9999 |
| ΔSCO0713 vs. S7 | 0 | -0,3577 to 0,3577 | No | ns | >0,9999 |
| S3 vs. S5 | 0 | -0,3577 to 0,3577 | No | ns | >0,9999 |
| S3 vs. S7 | 0 | -0,3577 to 0,3577 | No | ns | >0,9999 |
| S5 vs. S7 | 0 | -0,3577 to 0,3577 | No | ns | >0,9999 |
| MHET | | | | | |
| Control vs. M145 | -0,0402 | -0,3979 to 0,3175 | No | ns | 0,9994 |
| Control vs. ΔSCO0713 | -0,7505 | -1,108 to -0,3928 | Yes | **** | <0,0001 |
| Control vs. S3 | -0,8812 | -1,239 to -0,5235 | Yes | **** | <0,0001 |
| Control vs. S5 | 0,02597 | -0,3317 to 0,3837 | No | ns | >0,9999 |
| Control vs. S7 | -0,0427 | -0,4004 to 0,3150 | No | ns | 0,9991 |
| M145 vs. ΔSCO0713 | -0,7103 | -1,068 to -0,3526 | Yes | **** | <0,0001 |
| M145 vs. S3 | -0,841 | -1,199 to -0,4833 | Yes | **** | <0,0001 |

| | | | | | |
|---|---|---|---|---|---|
| M145 vs. S5 | 0,06617 | -0,2915 to 0,4239 | No | ns | 0,9932 |
| M145 vs. S7 | -0,0025 | -0,3602 to 0,3552 | No | ns | >0,9999 |
| ΔSCO0713 vs. S3 | -0,1307 | -0,4884 to 0,2270 | No | ns | 0,8785 |
| ΔSCO0713 vs. S5 | 0,7765 | 0,4188 to 1,134 | Yes | **** | <0,0001 |
| ΔSCO0713 vs. S7 | 0,7078 | 0,3501 to 1,066 | Yes | **** | <0,0001 |
| S3 vs. S5 | 0,9072 | 0,5495 to 1,265 | Yes | **** | <0,0001 |
| S3 vs. S7 | 0,8385 | 0,4808 to 1,196 | Yes | **** | <0,0001 |
| S5 vs. S7 | -0,06867 | -0,4264 to 0,2890 | No | ns | 0,9919 |
| BHET | | | | | |
| Control vs. M145 | 0,04 | -0,3177 to 0,3977 | No | ns | 0,9994 |
| Control vs. ΔSCO0713 | 0,7533 | 0,3956 to 1,111 | Yes | **** | <0,0001 |
| Control vs. S3 | 0,8833 | 0,5256 to 1,241 | Yes | **** | <0,0001 |
| Control vs. S5 | -0,02333 | -0,3810 to 0,3344 | No | ns | >0,9999 |
| Control vs. S7 | 0,04333 | -0,3144 to 0,4010 | No | ns | 0,9991 |
| M145 vs. ΔSCO0713 | 0,7133 | 0,3556 to 1,071 | Yes | **** | <0,0001 |
| M145 vs. S3 | 0,8433 | 0,4856 to 1,201 | Yes | **** | <0,0001 |
| M145 vs. S5 | -0,06333 | -0,4210 to 0,2944 | No | ns | 0,9944 |
| M145 vs. S7 | 0,003333 | -0,3544 to 0,3610 | No | ns | >0,9999 |
| ΔSCO0713 vs. S3 | 0,13 | -0,2277 to 0,4877 | No | ns | 0,8808 |
| ΔSCO0713 vs. S5 | -0,7767 | -1,134 to -0,4190 | Yes | **** | <0,0001 |
| ΔSCO0713 vs. S7 | -0,71 | -1,068 to -0,3523 | Yes | **** | <0,0001 |
| S3 vs. S5 | -0,9067 | -1,264 to -0,5490 | Yes | **** | <0,0001 |
| S3 vs. S7 | -0,84 | -1,198 to -0,4823 | Yes | **** | <0,0001 |
| S5 vs. S7 | 0,06667 | -0,2910 to 0,4244 | No | ns | 0,9929 |

| Test details | Mean 1 | Mean 2 | Mean Diff, | SE of diff, | N1 | N2 | q | DF |
|---|---|---|---|---|---|---|---|---|
| TPA | | | | | | | | |
| Control vs. M145 | 0 | 0 | 0 | 0,1189 | 3 | 3 | 0 | 36 |
| Control vs. ΔSCO0713 | 0 | 0 | 0 | 0,1189 | 3 | 3 | 0 | 36 |
| Control vs. S3 | 0 | 0 | 0 | 0,1189 | 3 | 3 | 0 | 36 |

| | | | | | | | | |
|---|---|---|---|---|---|---|---|---|
| Control vs. S5 | 0 | 0 | 0 | 0,1189 | 3 | 3 | 0 | 36 |
| Control vs. S7 | 0 | 0 | 0 | 0,1189 | 3 | 3 | 0 | 36 |
| M145 vs. ΔSCO0713 | 0 | 0 | 0 | 0,1189 | 3 | 3 | 0 | 36 |
| M145 vs. S3 | 0 | 0 | 0 | 0,1189 | 3 | 3 | 0 | 36 |
| M145 vs. S5 | 0 | 0 | 0 | 0,1189 | 3 | 3 | 0 | 36 |
| M145 vs. S7 | 0 | 0 | 0 | 0,1189 | 3 | 3 | 0 | 36 |
| ΔSCO0713 vs. S3 | 0 | 0 | 0 | 0,1189 | 3 | 3 | 0 | 36 |
| ΔSCO0713 vs. S5 | 0 | 0 | 0 | 0,1189 | 3 | 3 | 0 | 36 |
| ΔSCO0713 vs. S7 | 0 | 0 | 0 | 0,1189 | 3 | 3 | 0 | 36 |
| S3 vs. S5 | 0 | 0 | 0 | 0,1189 | 3 | 3 | 0 | 36 |
| S3 vs. S7 | 0 | 0 | 0 | 0,1189 | 3 | 3 | 0 | 36 |
| S5 vs. S7 | 0 | 0 | 0 | 0,1189 | 3 | 3 | 0 | 36 |
| MHET | | | | | | | | |
| Control vs. M145 | 1,048 | 1,089 | -0,0402 | 0,1189 | 3 | 3 | 0,4782 | 36 |
| Control vs. ΔSCO0713 | 1,048 | 1,799 | -0,7505 | 0,1189 | 3 | 3 | 8,928 | 36 |
| Control vs. S3 | 1,048 | 1,93 | -0,8812 | 0,1189 | 3 | 3 | 10,48 | 36 |
| Control vs. S5 | 1,048 | 1,023 | 0,02597 | 0,1189 | 3 | 3 | 0,3089 | 36 |
| Control vs. S7 | 1,048 | 1,091 | -0,0427 | 0,1189 | 3 | 3 | 0,5079 | 36 |
| M145 vs. ΔSCO0713 | 1,089 | 1,799 | -0,7103 | 0,1189 | 3 | 3 | 8,45 | 36 |
| M145 vs. S3 | 1,089 | 1,93 | -0,841 | 0,1189 | 3 | 3 | 10 | 36 |
| M145 vs. S5 | 1,089 | 1,023 | 0,06617 | 0,1189 | 3 | 3 | 0,7871 | 36 |
| M145 vs. S7 | 1,089 | 1,091 | -0,0025 | 0,1189 | 3 | 3 | 0,02974 | 36 |
| ΔSCO0713 vs. S3 | 1,799 | 1,93 | -0,1307 | 0,1189 | 3 | 3 | 1,554 | 36 |
| ΔSCO0713 vs. S5 | 1,799 | 1,023 | 0,7765 | 0,1189 | 3 | 3 | 9,237 | 36 |
| ΔSCO0713 vs. S7 | 1,799 | 1,091 | 0,7078 | 0,1189 | 3 | 3 | 8,42 | 36 |
| S3 vs. S5 | 1,93 | 1,023 | 0,9072 | 0,1189 | 3 | 3 | 10,79 | 36 |
| S3 vs. S7 | 1,93 | 1,091 | 0,8385 | 0,1189 | 3 | 3 | 9,974 | 36 |
| S5 vs. S7 | 1,023 | 1,091 | -0,06867 | 0,1189 | 3 | 3 | 0,8168 | 36 |
| BHET | | | | | | | | |
| Control vs. M145 | 98,95 | 98,91 | 0,04 | 0,1189 | 3 | 3 | 0,4758 | 36 |
| Control vs. ΔSCO0713 | 98,95 | 98,2 | 0,7533 | 0,1189 | 3 | 3 | 8,961 | 36 |

| | | | | | | | | |
|---|---|---|---|---|---|---|---|---|
| Control vs. S3 | 98,95 | 98,07 | 0,8833 | 0,1189 | 3 | 3 | 10,51 | 36 |
| Control vs. S5 | 98,95 | 98,98 | -0,02333 | 0,1189 | 3 | 3 | 0,2776 | 36 |
| Control vs. S7 | 98,95 | 98,91 | 0,04333 | 0,1189 | 3 | 3 | 0,5155 | 36 |
| M145 vs. ΔSCO0713 | 98,91 | 98,2 | 0,7133 | 0,1189 | 3 | 3 | 8,485 | 36 |
| M145 vs. S3 | 98,91 | 98,07 | 0,8433 | 0,1189 | 3 | 3 | 10,03 | 36 |
| M145 vs. S5 | 98,91 | 98,98 | -0,06333 | 0,1189 | 3 | 3 | 0,7534 | 36 |
| M145 vs. S7 | 98,91 | 98,91 | 0,003333 | 0,1189 | 3 | 3 | 0,03965 | 36 |
| ΔSCO0713 vs. S3 | 98,2 | 98,07 | 0,13 | 0,1189 | 3 | 3 | 1,546 | 36 |
| ΔSCO0713 vs. S5 | 98,2 | 98,98 | -0,7767 | 0,1189 | 3 | 3 | 9,239 | 36 |
| ΔSCO0713 vs. S7 | 98,2 | 98,91 | -0,71 | 0,1189 | 3 | 3 | 8,446 | 36 |
| S3 vs. S5 | 98,07 | 98,98 | -0,9067 | 0,1189 | 3 | 3 | 10,78 | 36 |
| S3 vs. S7 | 98,07 | 98,91 | -0,84 | 0,1189 | 3 | 3 | 9,992 | 36 |
| S5 vs. S7 | 98,98 | 98,91 | 0,06667 | 0,1189 | 3 | 3 | 0,793 | 36 |

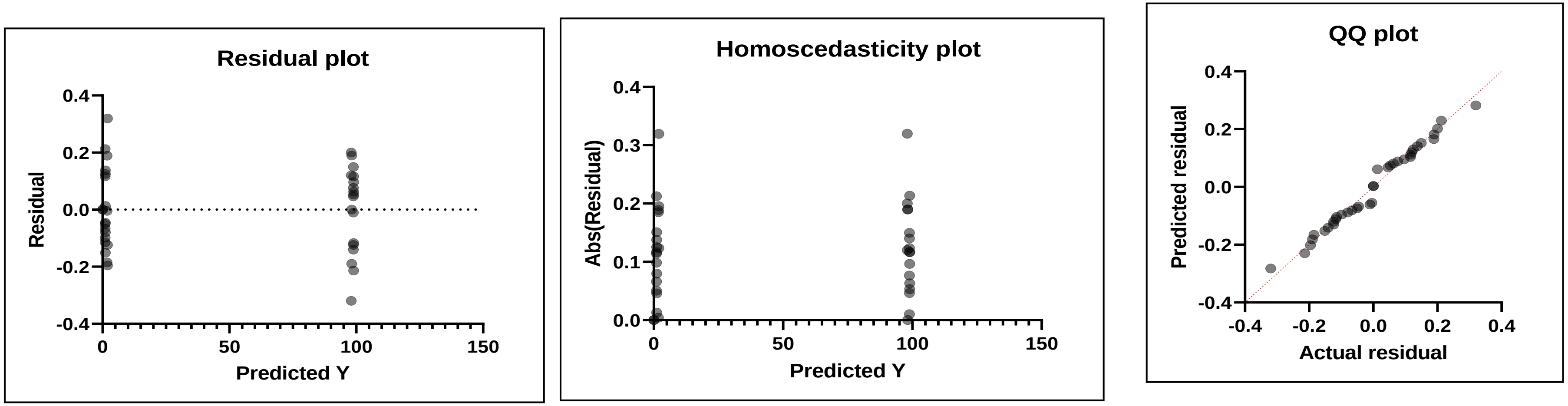

### All_NMM_48h

Within each column, compare rows (simple effects within columns)

| | |
|---|---|
| Number of families | 3 |
| Number of comparisons per family | 15 |
| Alpha | 0,05 |

| Tukey's multiple comparisons test | Mean Diff, | 95,00% CI of diff, | Below threshold? | Summary | Adjusted P Value |
|---|---|---|---|---|---|
| TPA | | | | | |
| Control vs. M145 | 0 | -3,595 to 3,595 | No | ns | >0,9999 |
| Control vs. ΔSCO0713 | 0 | -3,595 to 3,595 | No | ns | >0,9999 |
| Control vs. S3 | 0 | -3,595 to 3,595 | No | ns | >0,9999 |
| Control vs. S5 | 0 | -3,595 to 3,595 | No | ns | >0,9999 |
| Control vs. S7 | 0 | -3,595 to 3,595 | No | ns | >0,9999 |
| M145 vs. ΔSCO0713 | 0 | -3,595 to 3,595 | No | ns | >0,9999 |
| M145 vs. S3 | 0 | -3,595 to 3,595 | No | ns | >0,9999 |
| M145 vs. S5 | 0 | -3,595 to 3,595 | No | ns | >0,9999 |
| M145 vs. S7 | 0 | -3,595 to 3,595 | No | ns | >0,9999 |
| ΔSCO0713 vs. S3 | 0 | -3,595 to 3,595 | No | ns | >0,9999 |
| ΔSCO0713 vs. S5 | 0 | -3,595 to 3,595 | No | ns | >0,9999 |
| ΔSCO0713 vs. S7 | 0 | -3,595 to 3,595 | No | ns | >0,9999 |
| S3 vs. S5 | 0 | -3,595 to 3,595 | No | ns | >0,9999 |
| S3 vs. S7 | 0 | -3,595 to 3,595 | No | ns | >0,9999 |
| S5 vs. S7 | 0 | -3,595 to 3,595 | No | ns | >0,9999 |
| MHET | | | | | |
| Control vs. M145 | -9,337 | -12,93 to -5,742 | Yes | **** | <0,0001 |
| Control vs. ΔSCO0713 | -2,732 | -6,326 to 0,8629 | No | ns | 0,2259 |
| Control vs. S3 | -5,781 | -9,376 to -2,187 | Yes | *** | 0,0003 |
| Control vs. S5 | -5,21 | -8,805 to -1,616 | Yes | ** | 0,0014 |
| Control vs. S7 | -7,062 | -10,66 to -3,467 | Yes | **** | <0,0001 |
| M145 vs. ΔSCO0713 | 6,605 | 3,010 to 10,20 | Yes | **** | <0,0001 |
| M145 vs. S3 | 3,555 | -0,03919 to 7,150 | No | ns | 0,054 |

| | | | | | |
|---|---|---|---|---|---|
| M145 vs. S5 | 4,126 | 0,5318 to 7,721 | Yes | * | 0,0166 |
| M145 vs. S7 | 2,275 | -1,320 to 5,869 | No | ns | 0,4163 |
| ΔSCO0713 vs. S3 | -3,05 | -6,644 to 0,5449 | No | ns | 0,136 |
| ΔSCO0713 vs. S5 | -2,479 | -6,073 to 1,116 | No | ns | 0,3229 |
| ΔSCO0713 vs. S7 | -4,33 | -7,925 to -0,7358 | Yes | * | 0,0106 |
| S3 vs. S5 | 0,571 | -3,024 to 4,166 | No | ns | 0,9967 |
| S3 vs. S7 | -1,281 | -4,875 to 2,314 | No | ns | 0,8892 |
| S5 vs. S7 | -1,852 | -5,446 to 1,743 | No | ns | 0,6354 |
| BHET | | | | | |
| Control vs. M145 | 9,337 | 5,742 to 12,93 | Yes | **** | <0,0001 |
| Control vs. ΔSCO0713 | 2,733 | -0,8612 to 6,328 | No | ns | 0,2254 |
| Control vs. S3 | 5,783 | 2,189 to 9,378 | Yes | *** | 0,0003 |
| Control vs. S5 | 5,213 | 1,619 to 8,808 | Yes | ** | 0,0013 |
| Control vs. S7 | 7,063 | 3,469 to 10,66 | Yes | **** | <0,0001 |
| M145 vs. ΔSCO0713 | -6,603 | -10,20 to -3,009 | Yes | **** | <0,0001 |
| M145 vs. S3 | -3,553 | -7,148 to 0,04119 | No | ns | 0,0542 |
| M145 vs. S5 | -4,123 | -7,718 to -0,5288 | Yes | * | 0,0167 |
| M145 vs. S7 | -2,273 | -5,868 to 1,321 | No | ns | 0,417 |
| ΔSCO0713 vs. S3 | 3,05 | -0,5445 to 6,645 | No | ns | 0,1359 |
| ΔSCO0713 vs. S5 | 2,48 | -1,115 to 6,075 | No | ns | 0,3223 |
| ΔSCO0713 vs. S7 | 4,33 | 0,7355 to 7,925 | Yes | * | 0,0106 |
| S3 vs. S5 | -0,57 | -4,165 to 3,025 | No | ns | 0,9967 |
| S3 vs. S7 | 1,28 | -2,315 to 4,875 | No | ns | 0,8894 |
| S5 vs. S7 | 1,85 | -1,745 to 5,445 | No | ns | 0,6363 |

| Test details | Mean 1 | Mean 2 | Mean Diff, | SE of diff, | N1 | N2 | q | DF |
|---|---|---|---|---|---|---|---|---|
| TPA | | | | | | | | |
| Control vs. M145 | 0 | 0 | 0 | 1,195 | 3 | 3 | 0 | 36 |
| Control vs. ΔSCO0713 | 0 | 0 | 0 | 1,195 | 3 | 3 | 0 | 36 |
| Control vs. S3 | 0 | 0 | 0 | 1,195 | 3 | 3 | 0 | 36 |

| | | | | | | | | |
|---|---|---|---|---|---|---|---|---|
| Control vs. S5 | 0 | 0 | 0 | 1,195 | 3 | 3 | 0 | 36 |
| Control vs. S7 | 0 | 0 | 0 | 1,195 | 3 | 3 | 0 | 36 |
| M145 vs. ΔSCO0713 | 0 | 0 | 0 | 1,195 | 3 | 3 | 0 | 36 |
| M145 vs. S3 | 0 | 0 | 0 | 1,195 | 3 | 3 | 0 | 36 |
| M145 vs. S5 | 0 | 0 | 0 | 1,195 | 3 | 3 | 0 | 36 |
| M145 vs. S7 | 0 | 0 | 0 | 1,195 | 3 | 3 | 0 | 36 |
| ΔSCO0713 vs. S3 | 0 | 0 | 0 | 1,195 | 3 | 3 | 0 | 36 |
| ΔSCO0713 vs. S5 | 0 | 0 | 0 | 1,195 | 3 | 3 | 0 | 36 |
| ΔSCO0713 vs. S7 | 0 | 0 | 0 | 1,195 | 3 | 3 | 0 | 36 |
| S3 vs. S5 | 0 | 0 | 0 | 1,195 | 3 | 3 | 0 | 36 |
| S3 vs. S7 | 0 | 0 | 0 | 1,195 | 3 | 3 | 0 | 36 |
| S5 vs. S7 | 0 | 0 | 0 | 1,195 | 3 | 3 | 0 | 36 |
| MHET | | | | | | | | |
| Control vs. M145 | 2,577 | 11,91 | -9,337 | 1,195 | 3 | 3 | 11,05 | 36 |
| Control vs. ΔSCO0713 | 2,577 | 5,308 | -2,732 | 1,195 | 3 | 3 | 3,233 | 36 |
| Control vs. S3 | 2,577 | 8,358 | -5,781 | 1,195 | 3 | 3 | 6,843 | 36 |
| Control vs. S5 | 2,577 | 7,787 | -5,21 | 1,195 | 3 | 3 | 6,167 | 36 |
| Control vs. S7 | 2,577 | 9,639 | -7,062 | 1,195 | 3 | 3 | 8,359 | 36 |
| M145 vs. ΔSCO0713 | 11,91 | 5,308 | 6,605 | 1,195 | 3 | 3 | 7,818 | 36 |
| M145 vs. S3 | 11,91 | 8,358 | 3,555 | 1,195 | 3 | 3 | 4,208 | 36 |
| M145 vs. S5 | 11,91 | 7,787 | 4,126 | 1,195 | 3 | 3 | 4,884 | 36 |
| M145 vs. S7 | 11,91 | 9,639 | 2,275 | 1,195 | 3 | 3 | 2,692 | 36 |
| ΔSCO0713 vs. S3 | 5,308 | 8,358 | -3,05 | 1,195 | 3 | 3 | 3,61 | 36 |
| ΔSCO0713 vs. S5 | 5,308 | 7,787 | -2,479 | 1,195 | 3 | 3 | 2,934 | 36 |
| ΔSCO0713 vs. S7 | 5,308 | 9,639 | -4,33 | 1,195 | 3 | 3 | 5,126 | 36 |
| S3 vs. S5 | 8,358 | 7,787 | 0,571 | 1,195 | 3 | 3 | 0,6759 | 36 |
| S3 vs. S7 | 8,358 | 9,639 | -1,281 | 1,195 | 3 | 3 | 1,516 | 36 |
| S5 vs. S7 | 7,787 | 9,639 | -1,852 | 1,195 | 3 | 3 | 2,192 | 36 |
| BHET | | | | | | | | |
| Control vs. M145 | 97,42 | 88,09 | 9,337 | 1,195 | 3 | 3 | 11,05 | 36 |
| Control vs. ΔSCO0713 | 97,42 | 94,69 | 2,733 | 1,195 | 3 | 3 | 3,235 | 36 |

| | | | | | | | | |
|---|---|---|---|---|---|---|---|---|
| Control vs. S3 | 97,42 | 91,64 | 5,783 | 1,195 | 3 | 3 | 6,846 | 36 |
| Control vs. S5 | 97,42 | 92,21 | 5,213 | 1,195 | 3 | 3 | 6,171 | 36 |
| Control vs. S7 | 97,42 | 90,36 | 7,063 | 1,195 | 3 | 3 | 8,361 | 36 |
| M145 vs. ΔSCO0713 | 88,09 | 94,69 | -6,603 | 1,195 | 3 | 3 | 7,816 | 36 |
| M145 vs. S3 | 88,09 | 91,64 | -3,553 | 1,195 | 3 | 3 | 4,206 | 36 |
| M145 vs. S5 | 88,09 | 92,21 | -4,123 | 1,195 | 3 | 3 | 4,881 | 36 |
| M145 vs. S7 | 88,09 | 90,36 | -2,273 | 1,195 | 3 | 3 | 2,691 | 36 |
| ΔSCO0713 vs. S3 | 94,69 | 91,64 | 3,05 | 1,195 | 3 | 3 | 3,61 | 36 |
| ΔSCO0713 vs. S5 | 94,69 | 92,21 | 2,48 | 1,195 | 3 | 3 | 2,936 | 36 |
| ΔSCO0713 vs. S7 | 94,69 | 90,36 | 4,33 | 1,195 | 3 | 3 | 5,125 | 36 |
| S3 vs. S5 | 91,64 | 92,21 | -0,57 | 1,195 | 3 | 3 | 0,6747 | 36 |
| S3 vs. S7 | 91,64 | 90,36 | 1,28 | 1,195 | 3 | 3 | 1,515 | 36 |
| S5 vs. S7 | 92,21 | 90,36 | 1,85 | 1,195 | 3 | 3 | 2,19 | 36 |

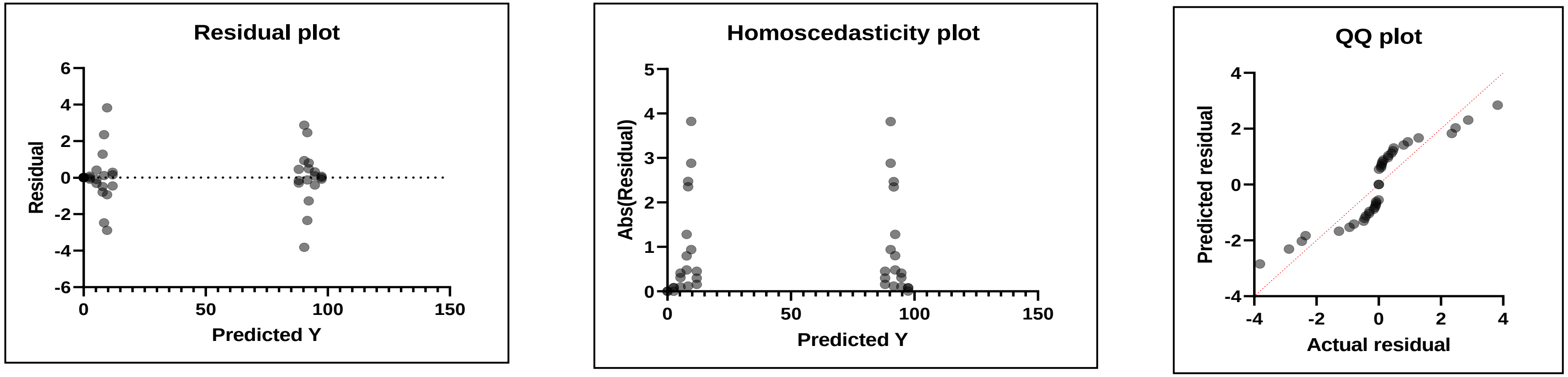

### All_NMM_72h

Within each column, compare rows (simple effects within columns)

| | |
|---|---|
| Number of families | 3 |
| Number of comparisons | 15 |
| Alpha | 0,05 |

| Tukey's multiple compari | Mean Diff, | 95,00% CI of diff, | Below threshold? | Summary | Adjusted P Value |
|---|---|---|---|---|---|
| TPA | | | | | |
| Control vs. M145 | 0 | -15,67 to 15,67 | No | ns | >0,9999 |
| Control vs. ΔSCO0713 | 0 | -15,67 to 15,67 | No | ns | >0,9999 |
| Control vs. S3 | 0 | -15,67 to 15,67 | No | ns | >0,9999 |
| Control vs. S5 | 0 | -15,67 to 15,67 | No | ns | >0,9999 |
| Control vs. S7 | 0 | -15,67 to 15,67 | No | ns | >0,9999 |
| M145 vs. ΔSCO0713 | 0 | -15,67 to 15,67 | No | ns | >0,9999 |
| M145 vs. S3 | 0 | -15,67 to 15,67 | No | ns | >0,9999 |
| M145 vs. S5 | 0 | -15,67 to 15,67 | No | ns | >0,9999 |
| M145 vs. S7 | 0 | -15,67 to 15,67 | No | ns | >0,9999 |
| ΔSCO0713 vs. S3 | 0 | -15,67 to 15,67 | No | ns | >0,9999 |
| ΔSCO0713 vs. S5 | 0 | -15,67 to 15,67 | No | ns | >0,9999 |
| ΔSCO0713 vs. S7 | 0 | -15,67 to 15,67 | No | ns | >0,9999 |
| S3 vs. S5 | 0 | -15,67 to 15,67 | No | ns | >0,9999 |
| S3 vs. S7 | 0 | -15,67 to 15,67 | No | ns | >0,9999 |
| S5 vs. S7 | 0 | -15,67 to 15,67 | No | ns | >0,9999 |
| MHET | | | | | |
| Control vs. M145 | -49,67 | -65,34 to -34,00 | Yes | **** | <0,0001 |
| Control vs. ΔSCO0713 | -17,05 | -32,72 to -1,382 | Yes | * | 0,0262 |
| Control vs. S3 | -39,06 | -54,73 to -23,39 | Yes | **** | <0,0001 |
| Control vs. S5 | -44,81 | -60,48 to -29,14 | Yes | **** | <0,0001 |
| Control vs. S7 | -49,73 | -65,40 to -34,06 | Yes | **** | <0,0001 |
| M145 vs. ΔSCO0713 | 32,62 | 16,95 to 48,29 | Yes | **** | <0,0001 |
| M145 vs. S3 | 10,61 | -5,057 to 26,28 | No | ns | 0,342 |

| | | | | | |
|---|---|---|---|---|---|
| M145 vs. S5 | 4,863 | -10,81 to 20,53 | No | ns | 0,935 |
| M145 vs. S7 | -0,05667 | -15,73 to 15,61 | No | ns | >0,9999 |
| ΔSCO0713 vs. S3 | -22,01 | -37,68 to -6,337 | Yes | ** | 0,002 |
| ΔSCO0713 vs. S5 | -27,76 | -43,43 to -12,09 | Yes | **** | <0,0001 |
| ΔSCO0713 vs. S7 | -32,68 | -48,35 to -17,01 | Yes | **** | <0,0001 |
| S3 vs. S5 | -5,75 | -21,42 to 9,920 | No | ns | 0,8765 |
| S3 vs. S7 | -10,67 | -26,34 to 5,000 | No | ns | 0,3363 |
| S5 vs. S7 | -4,92 | -20,59 to 10,75 | No | ns | 0,9319 |
| BHET | | | | | |
| Control vs. M145 | 49,67 | 34,00 to 65,34 | Yes | **** | <0,0001 |
| Control vs. ΔSCO0713 | 17,05 | 1,380 to 32,72 | Yes | * | 0,0263 |
| Control vs. S3 | 39,06 | 23,39 to 54,73 | Yes | **** | <0,0001 |
| Control vs. S5 | 44,81 | 29,14 to 60,48 | Yes | **** | <0,0001 |
| Control vs. S7 | 49,73 | 34,06 to 65,40 | Yes | **** | <0,0001 |
| M145 vs. ΔSCO0713 | -32,62 | -48,29 to -16,95 | Yes | **** | <0,0001 |
| M145 vs. S3 | -10,61 | -26,28 to 5,057 | No | ns | 0,342 |
| M145 vs. S5 | -4,863 | -20,53 to 10,81 | No | ns | 0,935 |
| M145 vs. S7 | 0,05667 | -15,61 to 15,73 | No | ns | >0,9999 |
| ΔSCO0713 vs. S3 | 22,01 | 6,337 to 37,68 | Yes | ** | 0,002 |
| ΔSCO0713 vs. S5 | 27,76 | 12,09 to 43,43 | Yes | **** | <0,0001 |
| ΔSCO0713 vs. S7 | 32,68 | 17,01 to 48,35 | Yes | **** | <0,0001 |
| S3 vs. S5 | 5,75 | -9,920 to 21,42 | No | ns | 0,8765 |
| S3 vs. S7 | 10,67 | -5,000 to 26,34 | No | ns | 0,3363 |
| S5 vs. S7 | 4,92 | -10,75 to 20,59 | No | ns | 0,9319 |

| Test details | Mean 1 | Mean 2 | Mean Diff, | SE of diff, | N1 | N2 | q | DF |
|---|---|---|---|---|---|---|---|---|
| TPA | | | | | | | | |
| Control vs. M145 | 0 | 0 | 0 | 5,208 | 3 | 3 | 0 | 36 |
| Control vs. ΔSCO0713 | 0 | 0 | 0 | 5,208 | 3 | 3 | 0 | 36 |
| Control vs. S3 | 0 | 0 | 0 | 5,208 | 3 | 3 | 0 | 36 |

| | | | | | | | | |
|---|---|---|---|---|---|---|---|---|
| Control vs. S5 | 0 | 0 | 0 | 5,208 | 3 | 3 | 0 | 36 |
| Control vs. S7 | 0 | 0 | 0 | 5,208 | 3 | 3 | 0 | 36 |
| M145 vs. ΔSCO0713 | 0 | 0 | 0 | 5,208 | 3 | 3 | 0 | 36 |
| M145 vs. S3 | 0 | 0 | 0 | 5,208 | 3 | 3 | 0 | 36 |
| M145 vs. S5 | 0 | 0 | 0 | 5,208 | 3 | 3 | 0 | 36 |
| M145 vs. S7 | 0 | 0 | 0 | 5,208 | 3 | 3 | 0 | 36 |
| ΔSCO0713 vs. S3 | 0 | 0 | 0 | 5,208 | 3 | 3 | 0 | 36 |
| ΔSCO0713 vs. S5 | 0 | 0 | 0 | 5,208 | 3 | 3 | 0 | 36 |
| ΔSCO0713 vs. S7 | 0 | 0 | 0 | 5,208 | 3 | 3 | 0 | 36 |
| S3 vs. S5 | 0 | 0 | 0 | 5,208 | 3 | 3 | 0 | 36 |
| S3 vs. S7 | 0 | 0 | 0 | 5,208 | 3 | 3 | 0 | 36 |
| S5 vs. S7 | 0 | 0 | 0 | 5,208 | 3 | 3 | 0 | 36 |
| MHET | | | | | | | | |
| Control vs. M145 | 3,748 | 53,42 | -49,67 | 5,208 | 3 | 3 | 13,49 | 36 |
| Control vs. ΔSCO0713 | 3,748 | 20,8 | -17,05 | 5,208 | 3 | 3 | 4,63 | 36 |
| Control vs. S3 | 3,748 | 42,81 | -39,06 | 5,208 | 3 | 3 | 10,61 | 36 |
| Control vs. S5 | 3,748 | 48,56 | -44,81 | 5,208 | 3 | 3 | 12,17 | 36 |
| Control vs. S7 | 3,748 | 53,48 | -49,73 | 5,208 | 3 | 3 | 13,5 | 36 |
| M145 vs. ΔSCO0713 | 53,42 | 20,8 | 32,62 | 5,208 | 3 | 3 | 8,857 | 36 |
| M145 vs. S3 | 53,42 | 42,81 | 10,61 | 5,208 | 3 | 3 | 2,882 | 36 |
| M145 vs. S5 | 53,42 | 48,56 | 4,863 | 5,208 | 3 | 3 | 1,321 | 36 |
| M145 vs. S7 | 53,42 | 53,48 | -0,05667 | 5,208 | 3 | 3 | 0,01539 | 36 |
| ΔSCO0713 vs. S3 | 20,8 | 42,81 | -22,01 | 5,208 | 3 | 3 | 5,975 | 36 |
| ΔSCO0713 vs. S5 | 20,8 | 48,56 | -27,76 | 5,208 | 3 | 3 | 7,537 | 36 |
| ΔSCO0713 vs. S7 | 20,8 | 53,48 | -32,68 | 5,208 | 3 | 3 | 8,873 | 36 |
| S3 vs. S5 | 42,81 | 48,56 | -5,75 | 5,208 | 3 | 3 | 1,561 | 36 |
| S3 vs. S7 | 42,81 | 53,48 | -10,67 | 5,208 | 3 | 3 | 2,897 | 36 |
| S5 vs. S7 | 48,56 | 53,48 | -4,92 | 5,208 | 3 | 3 | 1,336 | 36 |
| BHET | | | | | | | | |
| Control vs. M145 | 96,25 | 46,58 | 49,67 | 5,208 | 3 | 3 | 13,49 | 36 |
| Control vs. ΔSCO0713 | 96,25 | 79,2 | 17,05 | 5,208 | 3 | 3 | 4,63 | 36 |

| | | | | | | | | |
|---|---|---|---|---|---|---|---|---|
| Control vs. S3 | 96,25 | 57,19 | 39,06 | 5,208 | 3 | 3 | 10,6 | 36 |
| Control vs. S5 | 96,25 | 51,44 | 44,81 | 5,208 | 3 | 3 | 12,17 | 36 |
| Control vs. S7 | 96,25 | 46,52 | 49,73 | 5,208 | 3 | 3 | 13,5 | 36 |
| M145 vs. ΔSCO0713 | 46,58 | 79,2 | -32,62 | 5,208 | 3 | 3 | 8,857 | 36 |
| M145 vs. S3 | 46,58 | 57,19 | -10,61 | 5,208 | 3 | 3 | 2,882 | 36 |
| M145 vs. S5 | 46,58 | 51,44 | -4,863 | 5,208 | 3 | 3 | 1,321 | 36 |
| M145 vs. S7 | 46,58 | 46,52 | 0,05667 | 5,208 | 3 | 3 | 0,01539 | 36 |
| ΔSCO0713 vs. S3 | 79,2 | 57,19 | 22,01 | 5,208 | 3 | 3 | 5,975 | 36 |
| ΔSCO0713 vs. S5 | 79,2 | 51,44 | 27,76 | 5,208 | 3 | 3 | 7,537 | 36 |
| ΔSCO0713 vs. S7 | 79,2 | 46,52 | 32,68 | 5,208 | 3 | 3 | 8,873 | 36 |
| S3 vs. S5 | 57,19 | 51,44 | 5,75 | 5,208 | 3 | 3 | 1,561 | 36 |
| S3 vs. S7 | 57,19 | 46,52 | 10,67 | 5,208 | 3 | 3 | 2,897 | 36 |
| S5 vs. S7 | 51,44 | 46,52 | 4,92 | 5,208 | 3 | 3 | 1,336 | 36 |

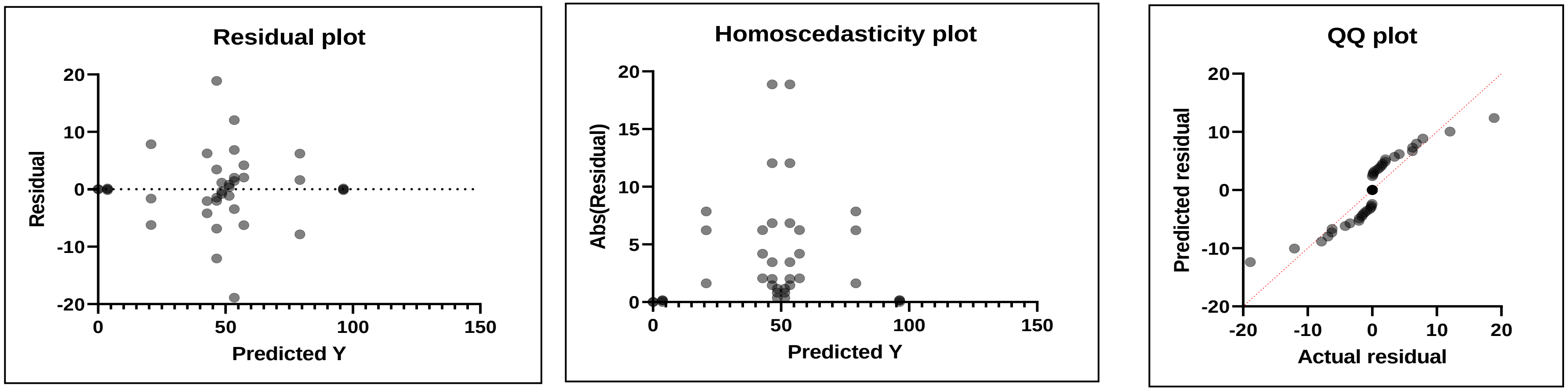

#### S9. Statistical analysis of Enzyme assays

| | | | | | |
|---|---|---|---|---|---|
| Table Analyzed | **24h Data BHET color** | | | | |
| Data sets analyzed | A-F | | | | |
| ANOVA summary | | | | | |
| F | 667,9 | | | | |
| P value | <0,0001 | | | | |
| P value summary | **** | | | | |
| Significant diff. among me | Yes | | | | |
| R squared | 0,9967 | | | | |
| Brown-Forsythe test | | | | | |
| F (DFn, DFd) | 0,2800 (5, 11) | | | | |
| P value | 0,9145 | | | | |
| P value summary | ns | | | | |
| Are SDs significantly diffe | No | | | | |
| Bartlett's test | | | | | |
| Bartlett's statistic (corrected) | | | | | |
| P value | | | | | |
| P value summary | | | | | |
| Are SDs significantly different (P < 0.05)? | | | | | |
| ANOVA table | SS | DF | MS | F (DFn, DFd) | P value |
| Treatment (between colur | 1,81 | 5 | 0,3621 | F (5, 11) = 667,9 | P<0,0001 |
| Residual (within columns) | 0,005963 | 11 | 0,0005421 | | |
| Total | 1,816 | 16 | | | |
| Data summary | | | | | |
| Number of treatments (co | 6 | | | | |
| Number of values (total) | 17 | | | | |
| Number of families | 1 | | | | |
| Number of comparisons p | 15 | | | | |

| | |
|---|---|
| Alpha | 0,05 |

| Tukey's multiple comparis | Mean Diff, | 95,00% CI of diff, | Below threshold? | Summary | Adjusted P Value | |
|---|---|---|---|---|---|---|
| IsPETase vs. TfCut2 | 0,4331 | 0,3683 to 0,4980 | Yes | **** | <0,0001 | A-B |
| IsPETase vs. 2LipA | 0,3045 | 0,2397 to 0,3694 | Yes | **** | <0,0001 | A-C |
| IsPETase vs. 92LipA | 0,9079 | 0,8354 to 0,9804 | Yes | **** | <0,0001 | A-D |
| IsPETase vs. ScLipA | 0,8151 | 0,7503 to 0,8799 | Yes | **** | <0,0001 | A-E |
| IsPETase vs. Buffer | 0,8232 | 0,7583 to 0,8880 | Yes | **** | <0,0001 | A-F |
| TfCut2 vs. 2LipA | -0,1286 | -0,1934 to -0,06376 | Yes | *** | 0,0003 | B-C |
| TfCut2 vs. 92LipA | 0,4747 | 0,4023 to 0,5472 | Yes | **** | <0,0001 | B-D |
| TfCut2 vs. ScLipA | 0,382 | 0,3171 to 0,4468 | Yes | **** | <0,0001 | B-E |
| TfCut2 vs. Buffer | 0,39 | 0,3252 to 0,4549 | Yes | **** | <0,0001 | B-F |
| 2LipA vs. 92LipA | 0,6033 | 0,5308 to 0,6758 | Yes | **** | <0,0001 | C-D |
| 2LipA vs. ScLipA | 0,5106 | 0,4457 to 0,5754 | Yes | **** | <0,0001 | C-E |
| 2LipA vs. Buffer | 0,5186 | 0,4538 to 0,5835 | Yes | **** | <0,0001 | C-F |
| 92LipA vs. ScLipA | -0,09278 | -0,1653 to -0,02030 | Yes | * | 0,0109 | D-E |
| 92LipA vs. Buffer | -0,08469 | -0,1572 to -0,01220 | Yes | * | 0,0199 | D-F |
| ScLipA vs. Buffer | 0,008092 | -0,05674 to 0,07293 | No | ns | 0,9977 | E-F |

| Test details | Mean 1 | Mean 2 | Mean Diff, | SE of diff, | n1 | n2 | q | DF |
|---|---|---|---|---|---|---|---|---|
| IsPETase vs. TfCut2 | 0,9693 | 0,5362 | 0,4331 | 0,01901 | 3 | 3 | 32,22 | 11 |
| IsPETase vs. 2LipA | 0,9693 | 0,6648 | 0,3045 | 0,01901 | 3 | 3 | 22,65 | 11 |
| IsPETase vs. 92LipA | 0,9693 | 0,06144 | 0,9079 | 0,02125 | 3 | 2 | 60,41 | 11 |
| IsPETase vs. ScLipA | 0,9693 | 0,1542 | 0,8151 | 0,01901 | 3 | 3 | 60,63 | 11 |
| IsPETase vs. Buffer | 0,9693 | 0,1461 | 0,8232 | 0,01901 | 3 | 3 | 61,24 | 11 |
| TfCut2 vs. 2LipA | 0,5362 | 0,6648 | -0,1286 | 0,01901 | 3 | 3 | 9,566 | 11 |
| TfCut2 vs. 92LipA | 0,5362 | 0,06144 | 0,4747 | 0,02125 | 3 | 2 | 31,59 | 11 |
| TfCut2 vs. ScLipA | 0,5362 | 0,1542 | 0,382 | 0,01901 | 3 | 3 | 28,41 | 11 |
| TfCut2 vs. Buffer | 0,5362 | 0,1461 | 0,39 | 0,01901 | 3 | 3 | 29,02 | 11 |
| 2LipA vs. 92LipA | 0,6648 | 0,06144 | 0,6033 | 0,02125 | 3 | 2 | 40,14 | 11 |
| 2LipA vs. ScLipA | 0,6648 | 0,1542 | 0,5106 | 0,01901 | 3 | 3 | 37,98 | 11 |
| 2LipA vs. Buffer | 0,6648 | 0,1461 | 0,5186 | 0,01901 | 3 | 3 | 38,58 | 11 |
| 92LipA vs. ScLipA | 0,06144 | 0,1542 | -0,09278 | 0,02125 | 2 | 3 | 6,173 | 11 |

| | | | | | | | | |
|---|---|---|---|---|---|---|---|---|
| 92LipA vs. Buffer | 0,06144 | 0,1461 | -0,08469 | 0,02125 | 2 | 3 | 5,635 | 11 |
| ScLipA vs. Buffer | 0,1542 | 0,1461 | 0,008092 | 0,01901 | 3 | 3 | 0,6019 | 11 |

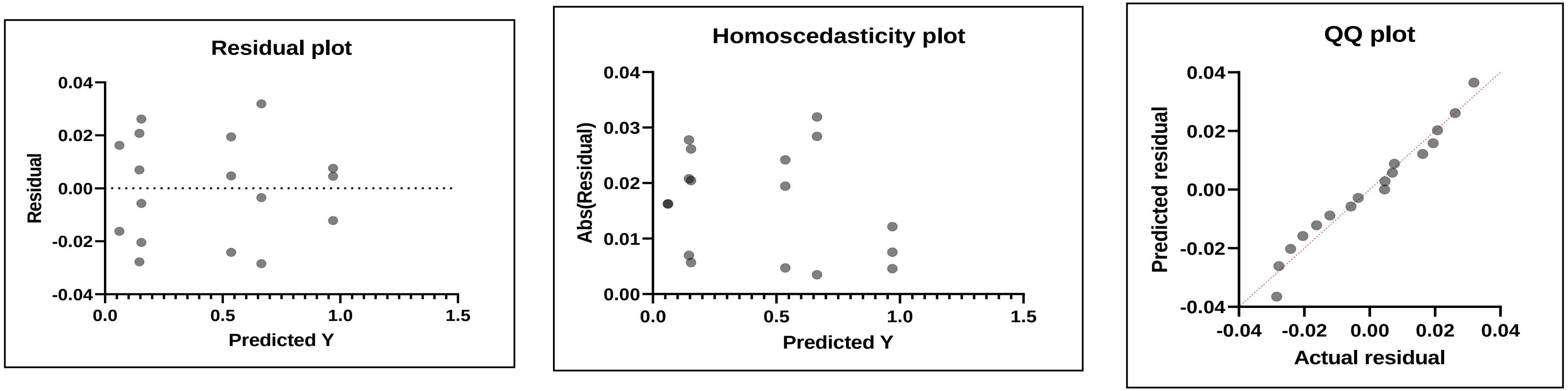

| | | | | | |
|---|---|---|---|---|---|
| Table Analyzed | 48h Data BHET color | | | | |
| Data sets analyzed | A-F | | | | |
| ANOVA summary | | | | | |
| F | 161,6 | | | | |
| P value | <0,0001 | | | | |
| P value summary | **** | | | | |
| Significant diff. among means (P < 0.05)? | Yes | | | | |
| R squared | 0,9878 | | | | |
| Brown-Forsythe test | | | | | |
| F (DFn, DFd) | | | | | |
| P value | | | | | |
| P value summary | | | | | |
| Are SDs significantly different (P < 0.05)? | | | | | |
| Bartlett's test | | | | | |
| Bartlett's statistic (corrected) | | | | | |
| P value | | | | | |
| P value summary | | | | | |
| Are SDs significantly different (P < 0.05)? | | | | | |
| ANOVA table | SS | DF | MS | F (DFn, DF | P value |
| Treatment (between columns) | 1,42 | 5 | 0,284 | F (5, 10) = | P<0,0001 |
| Residual (within columns) | 0,01757 | 10 | 0,001757 | | |
| Total | 1,438 | 15 | | | |
| Data summary | | | | | |
| Number of treatments (columns) | 6 | | | | |
| Number of values (total) | 16 | | | | |
| Number of families | 1 | | | | |

| | |
|---|---|
| Number of comparisons per family | 15 |
| Alpha | 0,05 |

| Tukey's multiple comparisons test | Mean Diff, | 95,00% CI of diff, | Below thres | Summary | Adjusted P | Value |
|---|---|---|---|---|---|---|
| IsPETase vs. TfCut2 | 0,3708 | 0,2519 to 0,4897 | Yes | **** | <0,0001 | A-B |
| IsPETase vs. 2LipA | 0,2565 | 0,1377 to 0,3754 | Yes | *** | 0,0002 | A-C |
| IsPETase vs. 92LipA | 0,8551 | 0,6869 to 1,023 | Yes | **** | <0,0001 | A-D |
| IsPETase vs. ScLipA | 0,7232 | 0,6043 to 0,8421 | Yes | **** | <0,0001 | A-E |
| IsPETase vs. Buffer | 0,7628 | 0,6439 to 0,8816 | Yes | **** | <0,0001 | A-F |
| TfCut2 vs. 2LipA | -0,1143 | -0,2332 to 0,004589 | No | ns | 0,0613 | B-C |
| TfCut2 vs. 92LipA | 0,4842 | 0,3161 to 0,6524 | Yes | **** | <0,0001 | B-D |
| TfCut2 vs. ScLipA | 0,3524 | 0,2335 to 0,4713 | Yes | **** | <0,0001 | B-E |
| TfCut2 vs. Buffer | 0,3919 | 0,2730 to 0,5108 | Yes | **** | <0,0001 | B-F |
| 2LipA vs. 92LipA | 0,5985 | 0,4304 to 0,7667 | Yes | **** | <0,0001 | C-D |
| 2LipA vs. ScLipA | 0,4667 | 0,3478 to 0,5856 | Yes | **** | <0,0001 | C-E |
| 2LipA vs. Buffer | 0,5062 | 0,3873 to 0,6251 | Yes | **** | <0,0001 | C-F |
| 92LipA vs. ScLipA | -0,1319 | -0,3000 to 0,03628 | No | ns | 0,1536 | D-E |
| 92LipA vs. Buffer | -0,09231 | -0,2604 to 0,07583 | No | ns | 0,45 | D-F |
| ScLipA vs. Buffer | 0,03955 | -0,07934 to 0,1584 | No | ns | 0,8476 | E-F |

| Test details | Mean 1 | Mean 2 | Mean Diff, | SE of diff, | n1 | n2 | q | DF |
|---|---|---|---|---|---|---|---|---|
| IsPETase vs. TfCut2 | 0,8553 | 0,4845 | 0,3708 | 0,03423 | 3 | 3 | 15,32 | 10 |
| IsPETase vs. 2LipA | 0,8553 | 0,5988 | 0,2565 | 0,03423 | 3 | 3 | 10,6 | 10 |
| IsPETase vs. 92LipA | 0,8553 | 0,0002843 | 0,8551 | 0,04841 | 3 | 1 | 24,98 | 10 |
| IsPETase vs. ScLipA | 0,8553 | 0,1321 | 0,7232 | 0,03423 | 3 | 3 | 29,88 | 10 |
| IsPETase vs. Buffer | 0,8553 | 0,09259 | 0,7628 | 0,03423 | 3 | 3 | 31,51 | 10 |
| TfCut2 vs. 2LipA | 0,4845 | 0,5988 | -0,1143 | 0,03423 | 3 | 3 | 4,722 | 10 |
| TfCut2 vs. 92LipA | 0,4845 | 0,0002843 | 0,4842 | 0,04841 | 3 | 1 | 14,15 | 10 |
| TfCut2 vs. ScLipA | 0,4845 | 0,1321 | 0,3524 | 0,03423 | 3 | 3 | 14,56 | 10 |
| TfCut2 vs. Buffer | 0,4845 | 0,09259 | 0,3919 | 0,03423 | 3 | 3 | 16,19 | 10 |
| 2LipA vs. 92LipA | 0,5988 | 0,0002843 | 0,5985 | 0,04841 | 3 | 1 | 17,49 | 10 |
| 2LipA vs. ScLipA | 0,5988 | 0,1321 | 0,4667 | 0,03423 | 3 | 3 | 19,28 | 10 |
| 2LipA vs. Buffer | 0,5988 | 0,09259 | 0,5062 | 0,03423 | 3 | 3 | 20,92 | 10 |

| | | | | | | | | |
|---|---|---|---|---|---|---|---|---|
| 92LipA vs. ScLipA | 0,0002843 | 0,1321 | -0,1319 | 0,04841 | 1 | 3 | 3,852 | 10 |
| 92LipA vs. Buffer | 0,0002843 | 0,09259 | -0,09231 | 0,04841 | 1 | 3 | 2,697 | 10 |
| ScLipA vs. Buffer | 0,1321 | 0,09259 | 0,03955 | 0,03423 | 3 | 3 | 1,634 | 10 |

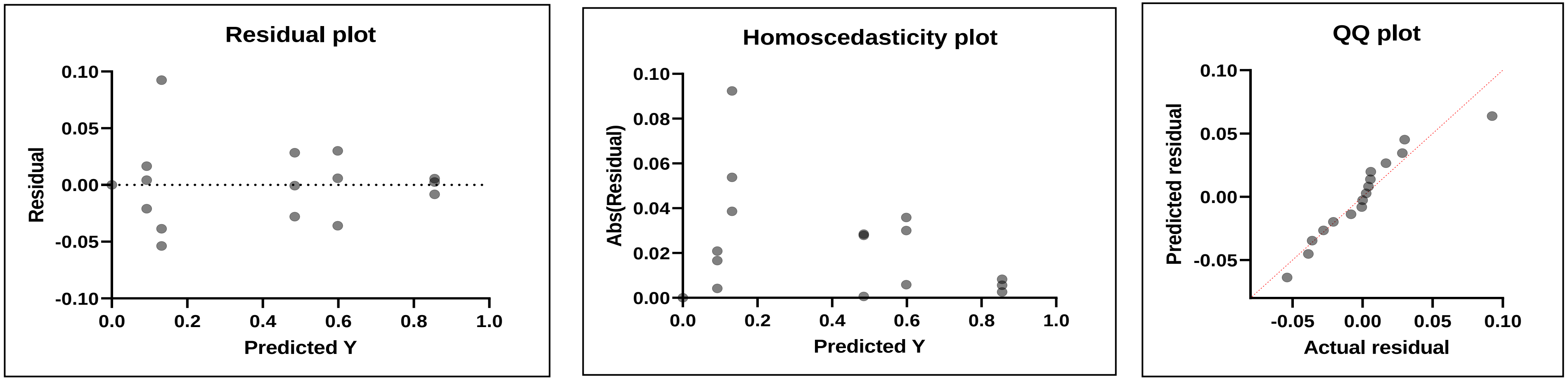

| | | | | | |
|---|---|---|---|---|---|
| Table Analyzed | 72hf Data BHET color | | | | |
| Data sets analyzed | A-C, E, F | | | | |
| ANOVA summary | | | | | |
| F | 370,4 | | | | |
| P value | <0,0001 | | | | |
| P value summary | **** | | | | |
| Significant diff. among means (P < 0.05)? | Yes | | | | |
| R squared | 0,9933 | | | | |
| Brown-Forsythe test | | | | | |
| F (DFn, DFd) | 0,6063 (4, 10) | | | | |
| P value | 0,6672 | | | | |
| P value summary | ns | | | | |
| Are SDs significantly different (P < 0.05)? | No | | | | |
| Bartlett's test | | | | | |
| Bartlett's statistic (corrected) | | | | | |
| P value | | | | | |
| P value summary | | | | | |
| Are SDs significantly different (P < 0.05)? | | | | | |
| ANOVA table | SS | DF | MS | F (DFn, DF | P value |
| Treatment (between columns) | 0,9227 | 4 | 0,2307 | F (4, 10) = | P<0,0001 |
| Residual (within columns) | 0,006228 | 10 | 0,000623 | | |
| Total | 0,9289 | 14 | | | |
| Data summary | | | | | |
| Number of treatments (columns) | 5 | | | | |
| Number of values (total) | 15 | | | | |
| Number of families | 1 | | | | |
| Number of comparisons per family | 10 | | | | |

| | |
|---|---|
| Alpha | 0,05 |

| Tukey's multiple comparisons test | Mean Diff, | 95,00% CI of diff, | Below thres | Summary | Adjusted P | Value |
|---|---|---|---|---|---|---|
| IsPETase vs. TfCut2 | 0,3847 | 0,3177 to 0,4518 | Yes | **** | <0,0001 | A-B |
| IsPETase vs. 2LipA | 0,2525 | 0,1854 to 0,3195 | Yes | **** | <0,0001 | A-C |
| IsPETase vs. ScLipA | 0,3992 | 0,3321 to 0,4662 | Yes | **** | <0,0001 | A-E |
| IsPETase vs. Buffer | 0,7653 | 0,6982 to 0,8323 | Yes | **** | <0,0001 | A-F |
| TfCut2 vs. 2LipA | -0,1323 | -0,1993 to -0,06520 | Yes | *** | 0,0005 | B-C |
| TfCut2 vs. ScLipA | 0,01444 | -0,05262 to 0,08150 | No | ns | 0,9498 | B-E |
| TfCut2 vs. Buffer | 0,3805 | 0,3135 to 0,4476 | Yes | **** | <0,0001 | B-F |
| 2LipA vs. ScLipA | 0,1467 | 0,07964 to 0,2138 | Yes | *** | 0,0002 | C-E |
| 2LipA vs. Buffer | 0,5128 | 0,4457 to 0,5799 | Yes | **** | <0,0001 | C-F |
| ScLipA vs. Buffer | 0,3661 | 0,2990 to 0,4332 | Yes | **** | <0,0001 | E-F |

| Test details | Mean 1 | Mean 2 | Mean Diff, | SE of diff, | n1 | n2 | q | DF |
|---|---|---|---|---|---|---|---|---|
| IsPETase vs. TfCut2 | 0,8296 | 0,4448 | 0,3847 | 0,02038 | 3 | 3 | 26,7 | 10 |
| IsPETase vs. 2LipA | 0,8296 | 0,5771 | 0,2525 | 0,02038 | 3 | 3 | 17,52 | 10 |
| IsPETase vs. ScLipA | 0,8296 | 0,4304 | 0,3992 | 0,02038 | 3 | 3 | 27,7 | 10 |
| IsPETase vs. Buffer | 0,8296 | 0,06429 | 0,7653 | 0,02038 | 3 | 3 | 53,11 | 10 |
| TfCut2 vs. 2LipA | 0,4448 | 0,5771 | -0,1323 | 0,02038 | 3 | 3 | 9,18 | 10 |
| TfCut2 vs. ScLipA | 0,4448 | 0,4304 | 0,01444 | 0,02038 | 3 | 3 | 1,002 | 10 |
| TfCut2 vs. Buffer | 0,4448 | 0,06429 | 0,3805 | 0,02038 | 3 | 3 | 26,41 | 10 |
| 2LipA vs. ScLipA | 0,5771 | 0,4304 | 0,1467 | 0,02038 | 3 | 3 | 10,18 | 10 |
| 2LipA vs. Buffer | 0,5771 | 0,06429 | 0,5128 | 0,02038 | 3 | 3 | 35,59 | 10 |
| ScLipA vs. Buffer | 0,4304 | 0,06429 | 0,3661 | 0,02038 | 3 | 3 | 25,41 | 10 |

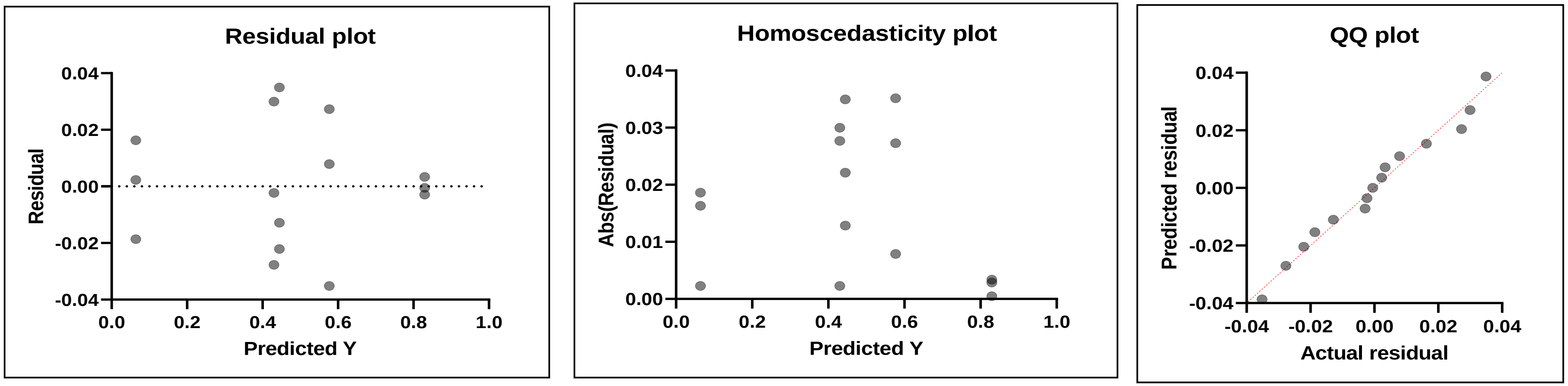
Residual plot
Residual
0.04
0.02
0.00
-0.02
-0.04
0.0
0.2
0.4
0.6
0.8
1.0
Predicted Y
Homoscedasticity plot
Abs(Residual)
0.04
0.03
0.02
0.01
0.00
0.0
0.2
0.4
0.6
0.8
1.0
Predicted Y
QQ plot
Predicted residual
0.04
0.02
0.00
-0.02
-0.04
-0.04
-0.02
0.00
0.02
0.04
Actual residual

#### S11. AlphaFold multimer

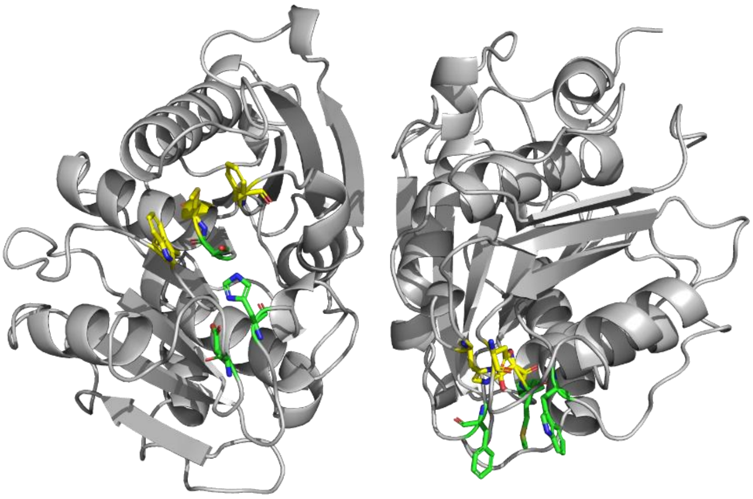

*Figure 3: AlphaFold multimer prediction of dimer complex.*
